# Orthologs of the *C. elegans* heterochronic genes have divergent functions in *C. briggsae*

**DOI:** 10.1101/2023.05.23.542001

**Authors:** Maria Ivanova, Eric G. Moss

**Affiliations:** Department of Molecular Biology, Graduate School of Biomedical Sciences, Rowan University, Stratford, NJ 08084

## Abstract

The heterochronic genes of *C. elegans* comprise the best-studied pathway controlling the timing of tissue and organ formation in an animal. To begin to understand the evolution of this pathway and the significance of the relationships among its components, we characterized 11 *C. briggsae* orthologs of *C. elegans* heterochronic genes. Using CRISPR/Cas9, we made a variety of alleles and found that several mutant phenotypes differ in significant ways from those of *C. elegans*. Although most mutant orthologs displayed defects in developmental timing, their phenotypes could differ in which stages were affected, the penetrance and expressivity of the phenotypes, or by having additional pleiotropies that were not obviously connected to developmental timing. However, when examining pairwise epistasis and synergistic relationships, we found those paralleled the known relationships between their *C. elegans* orthologs, suggesting that the arrangements of these genes in functional modules is conserved, but the modules’ relationships to each other and/or to their targets has drifted since the time of the species’ last common ancestor. Furthermore, our investigation has revealed a relationship to this pathway to other aspects of the animal’s growth and development, including gonad development, that is relevant to both species.

## Introduction

A developmental regulatory system performs its function in part due to the specific activities of its components and in part due to the manner in which these components interact. It has been found through comparative analysis that as these systems evolve, components may be replaced or their relationships may change. Such investigations can illuminate important features of a developmental regulatory system and how it performs its function (True and Haag 2001; Hill *et al*. 2006; Sommer 2012; Ellis 2022).

The heterochronic pathway of the nematode *Caenorahbditis elegans* is the most thoroughly characterized developmental regulatory system controlling timing of tissue and organ development in an animal (Rougvie and Moss 2013). The components of the core pathway include transcription factors, RNA-binding proteins, and several microRNAs (miRNAs). This is the pathway in which miRNAs were discovered, and they play an important role in how it works: several regulatory factors are down-regulated at specific times during larval development by the miRNAs. Furthermore, the transcription and processing of miRNAs are temporally regulated and, in some cases, under the control of other heterochronic regulators.

Mutations in heterochronic genes alter the relative timing of diverse developmental events independent of spatial or cell type specific regulation. Similar animal-wide timing pathways have not been characterized in other species. The core heterochronic pathway includes the protein-coding genes *lin-14, lin-28, lin-29, lin-41, lin-46* and *hbl-1*, and the miRNA-encoding *lin-4, let-7*, *mir-241, mir-28,* and *mir-84* (Rougvie and Moss 2013). (Several other genes with heterochronic effects are not considered here.)

Most of the proteins encoded by heterochronic genes are expressed in the beginning of postembryonic development whereas the miRNAs are not. The levels of the miRNAs rise during the larval stages and block the expression of proteins whose activities promote stage-specific developmental events. In general, when the miRNAs are missing or defective, developing mutant animals repeat some stage-specific events and postpone later events, which is called a reiterative phenotype. Mutations that delete miRNA binding sites from the 3’UTRs of their heterochronic gene targets also cause reiterative phenotypes. By contrast, when target genes are defective due to loss-of-function mutations, stage-specific events are skipped, which is called a precocious phenotype.

The core heterochronic genes of *C. elegans* have one-to-one orthologs in *C. briggsae*. Some have orthologs in other phyla, such as the miRNAs and *lin-28*, and others belong to conserved gene families, such as *hbl-1*, *lin-29*, and *lin-41* (Rougvie and Moss 2013). *lin-14* and *lin-46* are found only in the *Caenorhabditis* genus of nematodes. The degree of conservation does not correlate with how important a gene is in the regulation of development: *lin-14* has a key role in controlling L1 and L2 cell fates, and in many ways it is the paradigmatic heterochronic gene (Ambros and Horvitz 1987).

*C. elegans* and *C. briggsae* are remarkably similar nematodes despite being separated evolutionarily by 5 to 30 million years (Cutter 2008). They both occupy the same ecological niche and have nearly identical development down to their cell lineages (Zhao *et al*. 2008; Félix and Duveau 2012). Although they are nearly indistinguishable anatomically, only 60% of their loci are clear orthologs (Stein *et al*. 2003).

Comparative developmental studies—especially of the sex determination pathway in *C. elegans*, *C. briggsae*, and other *Caenorhabditis* species—have revealed that many alterations, shifts, and substitutions of components and their relationships are possible while preserving morphology, life history, and behavior (Ellis 2017, 2022). Such evolution in developmental pathways while the resulting morphology remains unchanged is a phenomenon called developmental systems drift (True and Haag 2001). Random mutations that do not dramatically decrease fitness may linger for several generations and become suppressed by other mutations. Over time, many genetic differences can accumulate, causing the roles of individual genes to change and components of developmental pathways to be replaced or change their relationships to their targets.

Our goal was to investigate the functional organization of the heterochronic pathway by seeing how much of its composition and arrangement are the same across a short evolutionary time—short enough so that orthologs are identifiable for all components, but where sufficient time has passed for developmental systems drift to have occurred. We began by mutating each *C. briggsae* ortholog of the 11 core heterochronic genes and characterizing their phenotypes, as well as comparing some well-characterized epistasis relationships. To study genes expected to have early embryonic lethal, pleiotropic, or infertile null phenotypes we used the auxin-inducible degron system in *C. briggsae* (Zhang *et al*. 2015; Hills-Muckey *et al*. 2021).

## Results

### Wild-type seam and intestinal cell fates are similar in C. briggsae and C. elegans

Heterochronic phenotypes of *C. elegans* can be reliably observed in the postembryonic lineages of the lateral hypodermal seam cells (Ambros and Horvitz 1984, 1987; Ambros 1989). Seam cells are located along each side of the newly hatched larva, dividing at each larval stage and differentiating at adulthood (Sulston and Horvitz 1977). We counted seam cells at each larval stage and observed their divisions in wildtype *C. briggsae* to see if the lineage patterns resembled those of *C. elegans*.

We found that *C. briggsae* L1 larvae had 10 seam cells within 3 hours of hatching (n = 10, Fig. S1). Seam cells were observed to divide in L1 larvae within 6 hours after hatching, with one of the daughters staying at the midline while the other moved dorsal or ventral to join the hypodermal syncytium. As a result, we saw that molting L1 larvae still had 10 seam cells (n = 10). Seam cells H1, V1-V4, and V6 divided symmetrically in L2 larvae, and late L2 or early L3 larvae had 15.5+/-0.5 seam cells (n = 16). Both L3 and L4 larvae still had asymmetrical divisions like those in the L1. L4 larvae and adult worms had 15-16 seam cells (Fig. 1A). All seam cells aligned along the midline and produced cuticular alae at adulthood. These observations indicate that the numbers and division patterns of seam cells in *C. briggsae* are like those of *C. elegans* at each larval stage: 10 in the L1 and 16 in the L2 and later.

**Fig. 1.**
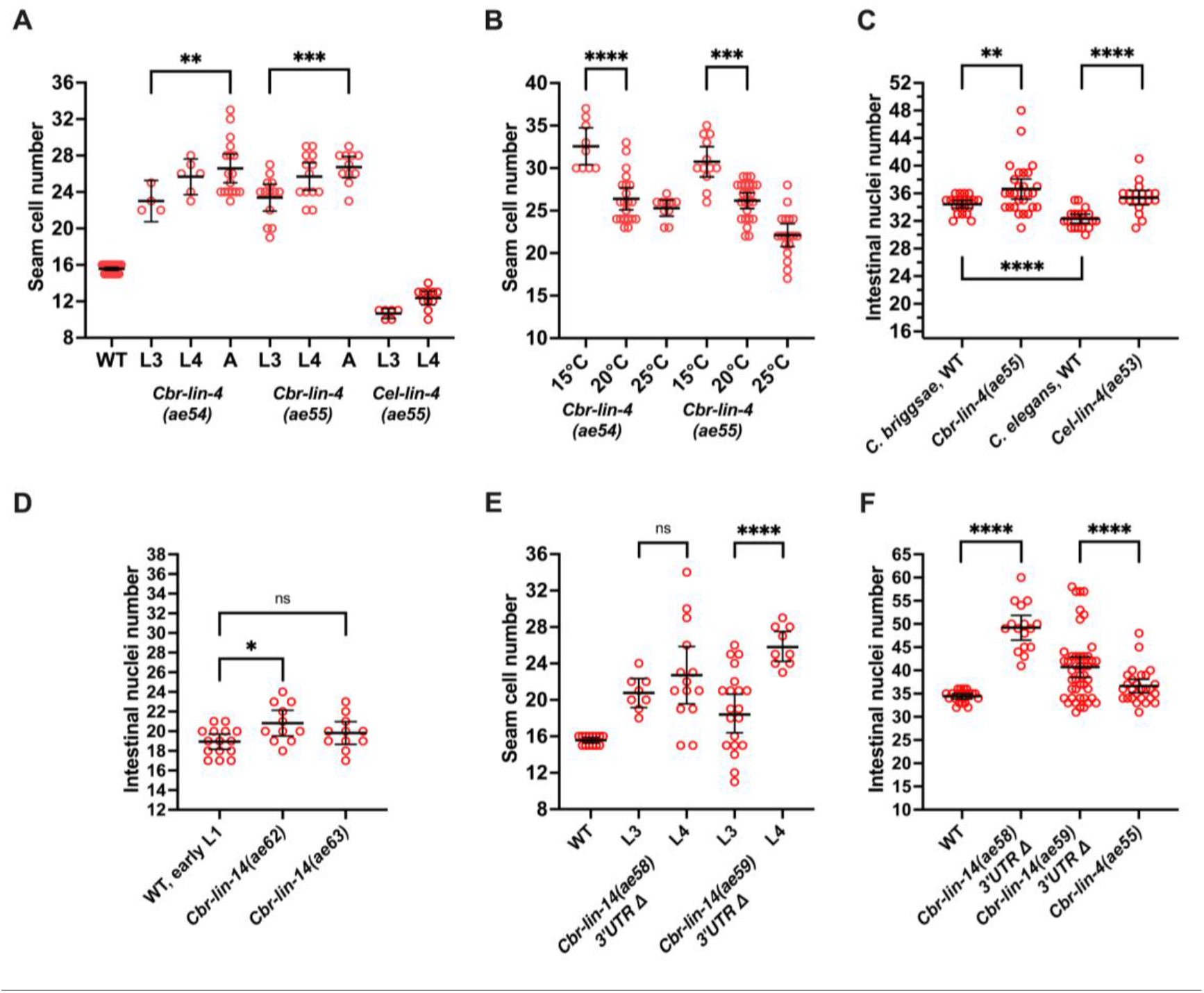
Seam cell and intestinal nuclei changes in *Cbr-lin-4(0)* and *Cbr-lin-14(0)* or *(gf)* mutants. Plots show seam cell and intestinal nuclei counts in L4 and young adults (unless otherwise specified) at 20°C (unless otherwise specified) for the strains indicated. (A) The number of seam cells is higher in *Cbr-lin-4(0)* mutants than in the wild type, and increases between the L3 and adult stages. On the contrary, *Cel-lin-4(0)* mutants have a reduced number of seam cells. (B) The number of seam cells in *Cbr-lin-4(0)* mutants increases at 15°C. (C) Intestinal nuclei numbers increase in *Cbr-lin-4(0)* mutants. (D) Intestinal nuclei numbers in the *Cbr-lin-14(0)* mutants are similar to those in early L1 larvae. (E) *Cbr-lin-14* 3’UTR deletion mutants have an increased number of seam cells. (F) *Cbr-lin-14* 3’UTR deletion mutants have even higher numbers of intestinal nuclei than *Cbr-lin-4(0)* mutants. The data set for *Cbr-lin-4(ae55)* is the same as in panel C. The reason for the difference between *ae58* and *ae59* is unclear as both deletions had approximately the same length and location. Statistical analysis is described in Materials and Methods.

Also like *C. elegans*, newly hatched *C. briggsae* had 20 intestinal nuclei. In *C. elegans*, 10 to 14 of 20 intestinal nuclei divide at the beginning of L1 lethargus (Sulston and Horvitz 1977; Hedgecock and White 1985). In *C. briggsae,* some intestinal cell nuclei also divided during the L1 lethargus and molt, with some divisions coinciding with the first round of the L2 seam cell divisions. In addition, we observed a slight but statistically significant increase in the average number of intestinal nuclei between the L3 and L4 stage. It is possible that some intestinal nuclei in *C. briggsae* divide during stages after the L1, which does not occur in *C. elegans* (Fig. S2).

We used seam cell and intestinal nuclei, as well as vulval lineages, which have been previously documented (Brown 2001), to characterize developmental timing phenotypes in *C. briggsae* mutants. However, we also observed phenotypic changes in other tissues, as described below.

### Cbr-lin-4(0) mutants reiterate L2 stages

*C. elegans lin-4* is the first heterochronic gene to be identified and encodes the first miRNA to be discovered (Chalfie *et al*. 1981; Ambros and Horvitz 1984; Lee *et al*. 1993). *Cel-lin-4(0)* mutants have a profound reiterative phenotype where L1-specific events are repeated causing adult animals to lack both vulvae and alae.

We isolated two *Cbr-lin-4(0)* mutant alleles with 6 (allele *ae54*) and 8 bp (allele *ae55*) deletions that remove most or all of the miRNA seed sequence (Table S1). Grossly, these animals were egg-laying defective for lacking a vulva (Fig. S3A and B).

To learn if *Cbr-lin-4(0)* mutants reiterated stage-specific fates like *C. elegans*, we examined seam cells, molts, adult alae, and intestinal nuclei. Based on seam cell counts, *Cbr-lin-4(0)* mutants appeared to re-iterate the symmetric divisions characteristic of the L2 stage during later larval stages (Fig. 1A, Fig. S4). Seam cells divided symmetrically at L2, and most larvae had 15.2+/-0.7 seam cells (n = 30) after the L2 divisions. Thus, L1 stages were not reiterated by most seam cells. This effect on seam cell lineages was also cold-sensitive: L4 larvae and young adults had more seam cells at 15°C than at 20°C or 25°C (Fig. 1B).

In contrast, both the *C. elegans* reference allele *Cel-lin-4(e912)* and *Cel-lin-4(ae53)*, a mutant generated using the same single guide RNA used for the *Cbr-lin-4(0)* mutant alleles, reiterated mostly L1 stage seam cell fates. As a result, these animals had 10.7+/-0.5 seam cells prior to the L4 stage (n = 6) and 12.7+/-1.7 seam cells after the L4 (n = 13).

Like *C. elegans lin-4(0)* mutants, *Cbr-lin-4(0)* mutant animals also have extra molts. Five L4 larvae from the *Cbr-lin-4(ae55)* strain were placed together on a plate at 20°C, and the next day, 7 shed cuticles were found in the bacterial lawn. The animals were adults (carried eggs), and 4 of them were stuck while shedding extra cuticles. Adult egg-producing wild-type *C. briggsae* were never observed shedding cuticles. Thus, *Cbr-lin-4(0)* mutants have at least one additional molt after reaching adulthood, and possibly as many as two. Unlike *Cel-lin-4(0)* mutants, whose cuticles are devoid of adult alae, *Cbr-lin-4(0)* mutants developed partial adult alae at the end of the development, most young adults (adults without embryos) had alae "patches", meaning that more than 0% but less than 50% of their seam cells generated adult alae (50%, n = 12), while most adult worms with embryos had “gapped” alae, meaning that more than 50%, but less than 100%, of the seam cells formed alae (78%, n = 18, Fig. S3C, Table S2). Thus, most seam cells differentiate at the end of the development in *C. briggsae lin-4* mutants, while they fail to differentiate in *C. elegans*.

In wild-type *C. elegans* and *C. briggsae*, dauer larvae represent an alternative developmental stage that forms in unfavorable environmental conditions like overcrowding or lack of nutrients. When heterochronic mutants reiterate L1 stages and do not transition to L2 they cannot form dauer larvae, which was observed for *Cel-lin-4(0)* mutants (Liu and Ambros 1989). Surprisingly, *Cbr-lin-4(0)* mutants could enter the dauer developmental pathway, reinforcing the conclusion that they enter the L2 stage. However, *Cbr-lin-4(0)* dauers had gapped dauer alae and some segments of their bodies looked expanded that was not observed in wild-type dauers (Fig. S5). We suspect that those worms transitioned to the dauer stage incompletely.

Interestingly, despite these differences, both *Cbr-lin-4(0)* and *Cel-lin-4(0)* mutants had a slightly increased number of intestinal nuclei compared to wild-type (Fig. 1C). The number of intestinal nuclei in *Cbr-lin-4(ae55)* mutant was not significantly different (*p* > 0.05, unpaired Welch’s t-test) at 15°C (36.9+/-3.1, n = 28) compared to the same mutant at 20°C. So, whereas the phenotype of *Cel-lin-4(0)* mutants is interpreted as a reiteration of L1 stage-specific fates, we find some ambiguity with *Cbr-lin-4(0)* mutants, since they appear to reiterate L1 fates in the intestine and L2 fates in the hypodermis.

### C. briggsae lin-14(0) *mutants resemble* C. elegans lin-14(0) *mutants*

*C. elegans lin-14* encodes a transcription factor unique to this genus (Ruvkun and Giusto 1989; Hristova *et al*. 2005). Alleles are of two general types: loss-of-function (lf) and null (0), which cause a precocious phenotype, and gain-of-function (gf), which lack miRNA binding sites in the 3’UTR and cause a reiterative phenotype (Ambros and Horvitz 1987; Wightman *et al*. 1991).

We made mutant alleles *ae62* and *ae63* with frameshift mutations that create premature stop codons in *Cbr-lin-14* by targeting the first exon shared by all isoforms (Table S1, Fig. S6). The mutants were often infertile, so we balanced them with a mutation that caused a visible phenotype: *Cbr-dpy-8(v262)* (Wei *et al*. 2014). *Cbr-lin-14(0)* progeny from the balanced strain resembled *Cel-lin-14(0)* mutants in several key features, including a protruding vulva, and shared a similar overall morphology (Fig. S7).

To see if these mutants displayed a precocious phenotype, we examined the L4 cuticle, seam cell divisions, and intestinal nuclei number. *Cbr-lin-14(0)* mutants developed full adult alae by the L4 stage (100% had precocious alae, n = 20). As occurs in *C. elegans lin-14(0)* mutants, seam cell counts of *Cbr-lin-14(0)* mutants are close to albeit slightly below the wild-type number by the L4 (Fig. S8). This is consistent with most seam cells of *Cbr-lin-14(0)* animals dividing symmetrically during the L1 stage, as occurs in *C. elegans lin-14(0)* mutants (Ambros and Horvitz, 1984). Also like *Cel-lin-14(0)*, the number of intestinal nuclei in later development was reduced in *Cbr-lin-14(0)* mutants, although some divisions did occur (Fig. 1D). Overall, our observations suggest *Cbr-lin-14* is required for the stage-appropriate expression of L1-specific fates and that the gene function is largely conserved between the two species.

### The Cbr-lin-4(0) reiterative phenotype requires functional Cbr-lin-14

In *C. elegans*, *lin-4* mutations that lead to a reiterative phenotype do so because they cause prolonged expression of *lin-14* (Ruvkun and Giusto 1989; Lee *et al*. 1993; Wightman *et al*. 1993). As a result, the phenotype of a loss-of-function *Cel-lin-14(0)* mutation is completely epistatic to that of *Cel-lin-4(0)* (Ambros 1989).

To test whether the temporal fate reiterations that occur in *Cbr-lin-4(0)* mutants occurred due to the elevated *Cbr-lin-14* function, we made *Cbr-lin-4(0); Cbr-lin-14(0)* double mutants by disrupting the *Cbr-lin-4* gene in a *Cbr-lin-14(ae62)* balanced strain using CRISPR/Cas9, and then isolating double homozygotes from among the progeny. We made strains with two different *Cbr-lin-4* alleles, *ae71* and *ae72* (Table S1).

The *Cbr-lin-4(0); Cbr-lin-14(0)* double mutant phenotype mostly resembled the *Cbr-lin-14(0)* single mutant phenotype—the number of seam cells by the L4 was 14.7+/- 0.9 (n = 16, combined data from two strains) and most of the intestinal nuclei did not divide; after the L1 stage, the number of intestinal nuclei was 19.6+/-1.5 (n = 16, combined data from two strains). Surprisingly, however, precocious alae were not always observed in double mutant worms: 2 of observed L4 larvae did not have precocious alae and other 2 had full precocious alae. This differs significantly from the *C. elegans* double mutant, in which alae appear precociously in all worms (Ambros 1989). This observation indicates a difference between the species in the relationships of *Cbr-lin-4* and *Cbr-lin-14* to downstream regulators controlling the timing of alae formation. But as in *C. elegans*, *Cbr-lin-14* appears to be required for the reiterative and vulvaless phenotypes of *Cbr-lin-4(0)*.

### Cbr-lin-14(gf) *mutants resemble weak* Cel-lin-14(gf) *alleles*

In *C. elegans*, two mutants with deletions in the 3’UTR of *lin-14* display a reiterative phenotype: an allele with nearly all miRNA sites removed reiterates L1 stages and lacks a vulva and adult alae, closely resembling *lin-4(0)*, and a weaker allele with a few intact miRNA sites reiterates both L1 and L2 stage events, also lacks a vulva, and develops some alae (Ambros and Horvitz 1987; Wightman *et al*. 1993).

*Cbr-lin-14* 3’UTR deletions (alleles *ae58* and *ae59*) were generated with CRISPR/Cas9, removing approximately 1.3 kb that includes all predicted miRNA binding sites (Table S1, Fig. S6). As adults, the mutants lacked vulvae, had extra seam cells and intestinal nuclei, and had alae patches (52%, n = 21) or lacked alae completely (48%, Table S2). The patches of adult alae were more transparent and thinner than wild-type alae.

The number of seam cells in *Cbr-lin-14(gf)* mutants was slightly lower and more variable at late stages than in *Cbr-lin-4(0)* mutants (Fig. 1A, E). Additionally, late L2 and early L3 larvae had fewer seam cells than expected if *Cbr-lin-14(gf)* phenocopied *Cbr-lin-4(0)* (mean = 11+/-1, n = 11). This difference would be explained by most seam cells reiterating L1 cell fates before they reiterate L2 fates. Also, the number of intestinal nuclei was higher than in *Cbr-lin-4(0)* mutants—a sign of reiteration of L1 fates in this tissue—which supports the interpretation that these animals reiterate L1 stage events to some degree. Thus, *Cbr-lin-14(gf)* mutants resemble the weaker *Cel-lin-14(gf)* allele.

### *Cbr-lin-28(0)* mutants have minor heterochronic defects and arrest development at the L4 stage

*C. elegans lin-28* encodes an RNA-binding protein that is widely conserved in animals (Moss *et al*. 1997; Moss and Tang 2003; Vadla *et al*. 2012). *Cel-lin-28(0)* mutants display a completely penetrant precocious phenotype where they skip cell fates of the L2, and a less penetrant defect of skipping L3 fates (Ambros and Horvitz 1984; Vadla *et al*. 2012). They also show an incompletely penetrant fertility problem as a result of spermathecal defects (Choi and Ambros 2019).

We made *Cbr-lin-28* mutant alleles *ae25*, *ae35*, and *ae39* by targeting the second exon to generate frameshifts with premature stop codons (Table S1, Fig. S6). *Cbr-lin-28* mutant animals were strikingly different from *Cel-lin-28(0)* animals: many arrested their development during the late L4 stage, did not undergo the final molt and lacked adult alae. These arrested animals retained features characteristic of mid- to late-wild-type L4 animals: the reflexed gonad arms stopped developing toward each other, and the vulva ceased development during morphogenesis (Fig. 2A). Observing a synchronized population, we found that *Cbr-lin-28* mutants developed at the same rate and produced oocytes at the same time as wild-type animals, except that wild type proceeded to the last molt and developed a vulva and alae (n≈20).

**Fig. 2.**
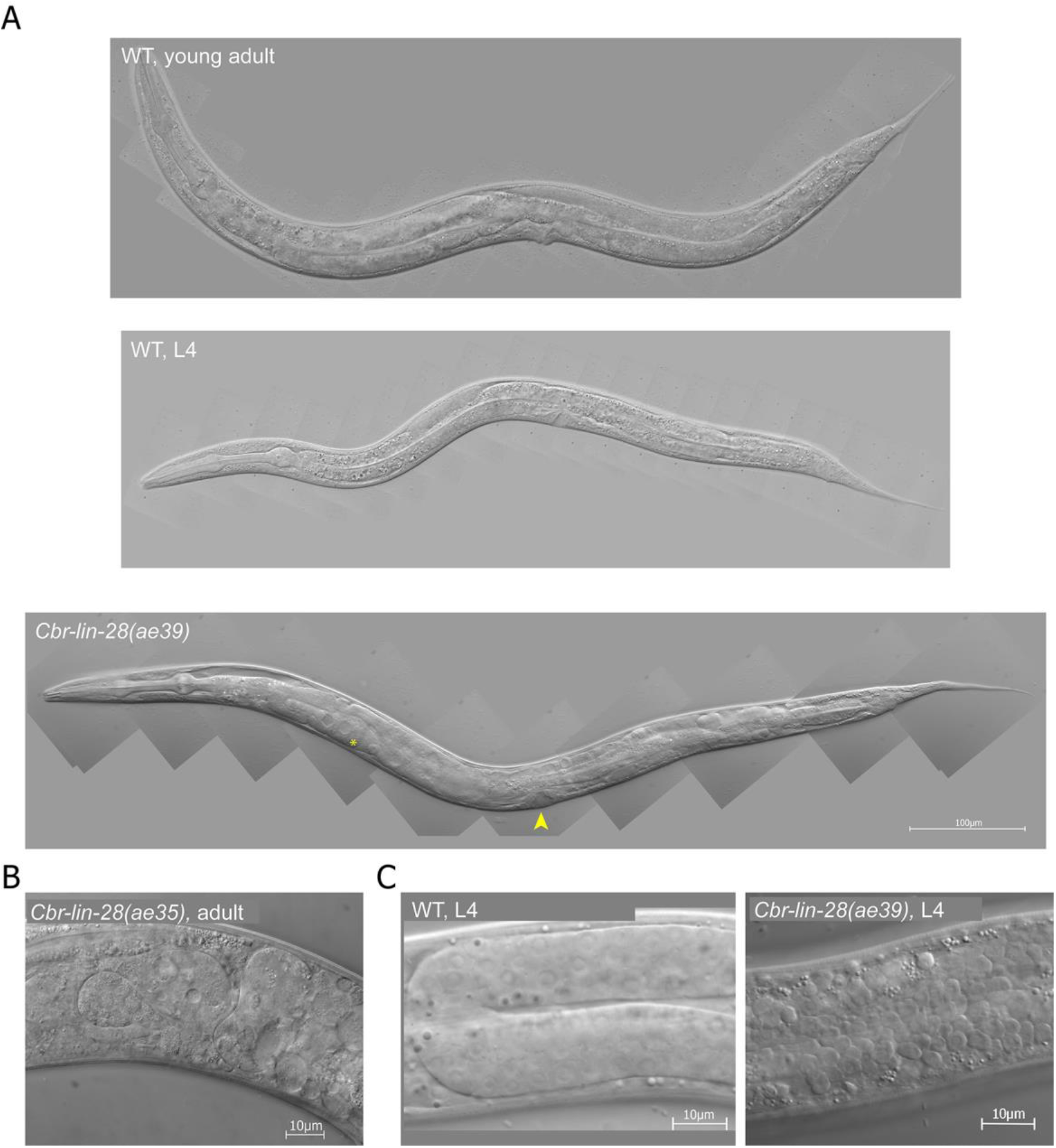
Developmental timing defects in *Cbr-lin-28(0)* mutants cause an L4 arrest and gonad disintegration. DIC photomicrographs of animals grown at 20°C. (A) Wild-type (AF16) L4 larva and young adult compared to *Cbr-lin-28(ae39)* with “arrested L4” phenotype. Arrowhead indicates L4-like vulva (larval trait) and asterisk indicates oocytes (adult trait). (B) A closer view at the disorganized gonad of a *Cbr-lin-28(ae35)* adult. (C) An earlier stage gonad disorganization that occurs in some *Cbr-lin-28(ae39)* mutant L4 larvae compared to a normal gonad. All animals are oriented anterior end left, dorsal side up.

The gonads of *Cbr-lin-28* mutants became disorganized or disintegrated after animals arrested in L4 or reached adulthood. Gonad contents leaked into the pseudocoelom and sometimes the gonad fell apart into separate cells (Fig. 2B). In some cases, gonad disorganization was not visible at first but manifested later (Fig. S9). The underlying defect is unknown. Some mutants developed through the L4 stage and had normal vulvae and adult alae, but became stuck in the L4 cuticle during molting. Those few that completed the L4 molt (“successfully molted”) were not morphologically different from the wild type, although sometimes they had disorganized gonads.

In contrast to what happens in *C. elegans*, *Cbr-lin-28* mutants had only weak heterochronic defects: most animals had a small patch of precocious alae near the pharynx at the L4 stage, and the number of seam cells was similar to the wild type (Fig. 3A). Undergoing dauer development suppressed precocious alae, but did not suppress L4 arrest (Fig. 3B and C). By contrast, dauer development completely suppresses *C. elegans lin-28(0)* heterochronic defects (Liu and Ambros 1991). Some other observed phenotypes included rolling (less than 10%), protruding vulvae (10-20%), larvae stuck at L2 and L3 molts, incompletely shed cuticles (“belts”), and sometimes gonads losing their structure during the L4 stage (Fig. 2C).

**Fig. 3.**
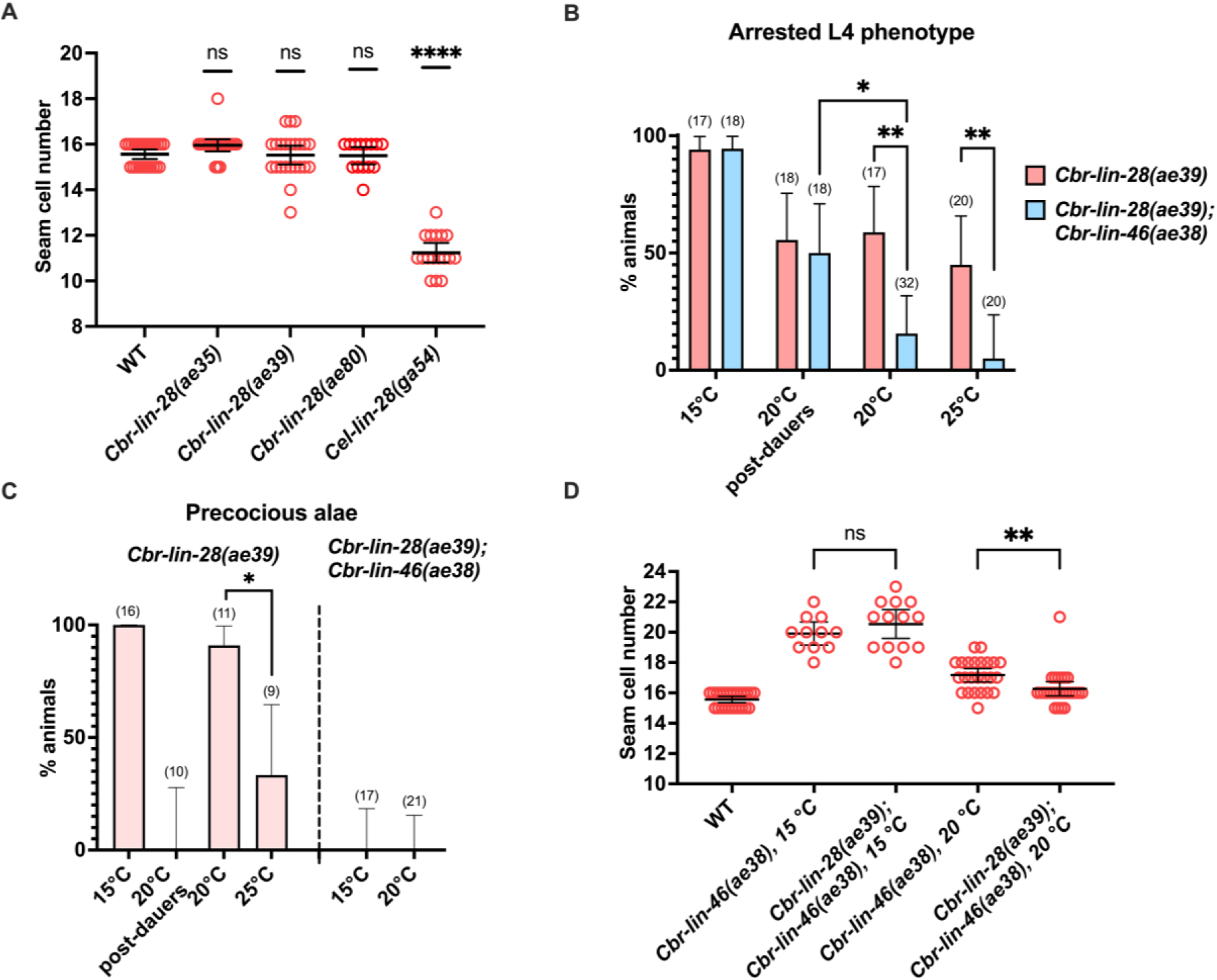
*Cbr-lin-28(0)* mutant phenotypes and their suppression by *Cbr-lin-46(0)* allele. (A) *Cbr-lin-28(0)* mutants do not have a reduced number of seam cells at 20°C, unlike *Cel-lin-28(0).* (B) L4 developmental arrest was more penetrant in the *Cbr-lin-28(0)* mutant at 15°C than at 20°C (both dauers and post-dauers, *p* ≤ 0.05). It also was not suppressed by *Cbr-lin-46(0)* mutation at 15°C in contrast to 20°C and 25°C. Those animals that did not arrest their development at L4, had “successfully molted” or intermediate phenotypes. Sample sizes are specified in parenthesis above the bars. (C) Precocious alae of *Cbr-lin-28(0)* occurred less often at 25°C and were suppressed by dauer pathway and *Cbr-lin-46(0)* mutation at 15°C and 20°C. (D) Increased seam cell number of *Cbr-lin-46(0)* mutant was suppressed by *Cbr-lin-28(0)* mutation at 20°C but not at 15°C. Statistical analysis is described in Materials and Methods.

*Cbr-lin-28*’s mutant phenotype was cold-sensitive. At 15°C, most animals became arrested L4s and all had precocious alae (n = 16) (Fig. 3B and C). Moreover, patches of precocious alae were longer and some animals had complete precocious alae. Additionally, the strain could not be maintained at 15°C. Some animals produced eggs at this temperature but they did not hatch, although eggs placed at 15°C after the mothers were grown at 25°C were viable and did hatch, suggesting a maternal effect embryonic problem at cold temperatures.

To confirm that the *Cbr-lin-28* mutant phenotypes that we observed are those of null alleles, we generated a deletion (*ae80*) that removed 78% of the 206 amino-acid coding region. This deletion starts within the CSD RNA-binding domain and deletes both CCHC zinc knuckles, so that the 46 remaining amino acids contain none of *Cbr-lin-28*’s known functional domains.

*Cbr-lin-28(ae80)* L4 larvae had 15.5+/-0.7 (n = 14) seam cells, and 42% (n = 12) had precocious alae patches. After 24 hours, all of 30 L4 larvae picked to a separate plate had vulvae arrested at early L4 stages (L4.2-L4.3; Mok *et al*., 2015), produced oocytes (most of them also had embryos), had L4 cuticles (60% had precocious alae patches, n = 15), and disorganized gonads.

Therefore, the phenotypes of *Cbr-lin-28(ae80)* worms were not significantly different from those of *ae35* and *ae39*. Heterochronic defects remained weak, and L2 seam cell divisions were not skipped. The penetrance of the "arrested L4" phenotype might be higher in this strain since no "successfully molted" worms were observed among the 30 isolated L4, although some advanced-stage worms (with adult-like vulvaе) were rarely observed on plates as well as worms with protruding vulvaе.

Overall, *Cbr-lin-28(0)* has only a minor resemblance to *Cel-lin-28(0)*, pleiotropic effects, and variable penetrance and expressivity for most phenotypes.

### Cbr-lin-28 is expressed at all stages in C. briggsae and down-regulated in seam cells after the L1 and L3 stages

In *C. elegans, lin-28* shows a characteristic “on early, off late” expression pattern that parallels its function in controlling L2 fates, and this temporal down-regulation is a consequence of miRNAs acting via its 3’UTR (Moss *et al*. 1997; Tsialikas *et al*. 2017). Because the phenotype of mutant *Cbr-lin-28* differed from that of its *C. elegans* ortholog, we examined the expression of *Cbr-lin-28* to see if that was different as well.

We employed a transgenic approach that had been successfully used in *C. elegans* which creates multicopy extrachromosomal arrays of plasmids (Stinchcomb *et al*. 1985; Mello *et al*. 1991). A full-length translational fusion with GFP that included intact 5’ and 3’ regulatory regions was constructed (Fig. S10) The construct was injected into wild-type *C. briggsae,* producing a stable extrachromosomal array *aeEx45*.

In transgenic animals, GFP fluorescence was observed in head and tail neurons, motor neurons, muscles (including the pharynx), intestinal cells, and seam cells (Fig. 4), which is similar to *C. elegans*, except that *Cel-lin-28::GFP* had obvious expression throughout the hypodermis (Moss *et al*. 1997). In contrast to *C. elegans*, most GFP expression did not appreciably decline with age.

**Fig. 4.**
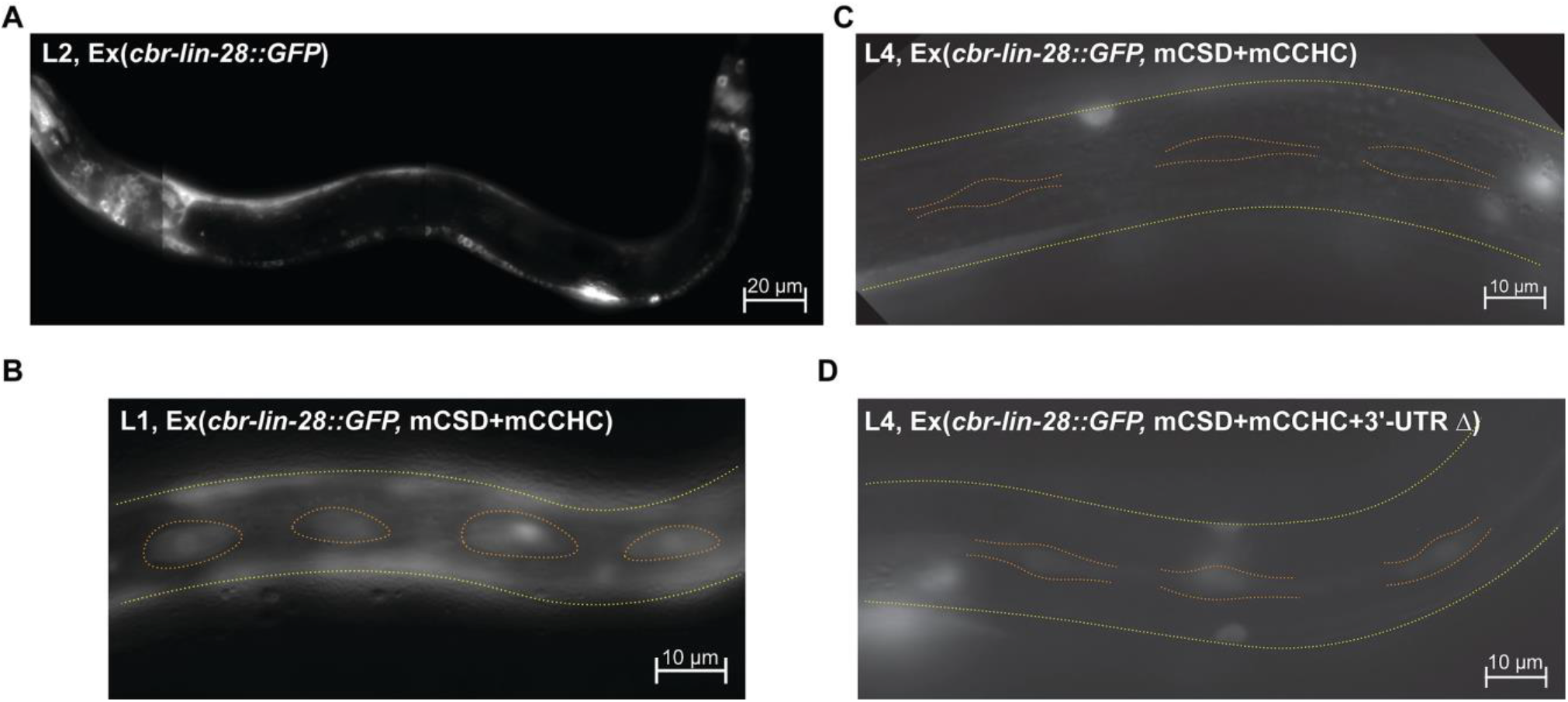
The expression of LIN-28 is down-regulated during *C. briggsae* development. Fluorescent microscopy images, green channel. All animals are oriented anterior end left, dorsal side up. Yellow dotted line indicates sides of animals, orange dotted line outlines seam cells. (A) An AF16 L2 larva from a brood of animals carrying *Cbr-lin-28::GFP* on an extrachromosomal array. The expression is visible in neurons, pharynx, and P-cells. (B) Fluorescing seam cells in an L1 larva expressing *Cbr-lin-28::GFP* with mutated CSD and CCHC domains. (C) Seam cells not glowing in an L4 larva expressing *Cbr-lin-28::GFP* array with mutated CSD and CCHC domains and an intact 3’UTR. (D) Seam cells glowing in an L4 larva with *Cbr-lin-28::GFP* array with mutated CSD and CCHC domains and a 3’UTR deletion.

About 30% of fluorescing animals also had alae gaps. Extrachromosomal arrays might exceed wild-type levels of expression since stable transgenes contain several copies of the gene (Mello *et al*. 1991). Thus, it is possible that our construct resulted in overexpression of *Cbr-lin-28—*possibly by overcoming miRNA repression—to cause a weak reiterative phenotype. Furthermore, we observed that embryos showing very bright GFP fluorescence failed to hatch or died soon after hatching; only 2 larvae hatched out of 34 brightly fluorescing eggs, suggesting that very high *Cbr-lin-28* expression causes an embryonic lethal phenotype.

To address the possibility that overexpression of the transgene affected *Cbr-lin-28* regulation, we generated *aeEx46* transgene with mutations in the *lin-28* protein’s two functional domains, the CSD (Y35A, F37A) and the CCHC zinc fingers (C127A, C137A) (Fig. S10). The GFP fluorescence was still observed at all stages and did not decline except for seam cells. In seam cells, robust fluorescence was observed mostly at the L1 stage. Weaker fluorescence was observed at L2 and L3 stages and occurred less often than at the L1 stage, and no fluorescence was observed at the L4 stage (Table 1).

**Table 1:** Percents of animals with fluorescent seam cells in strains with a *Cbr-lin-28::GFP* extrachromosomal array.

| Array |  | Stage |  |  |  |  |
| --- | --- | --- | --- | --- | --- | --- |
|  |  | L1 | L2 | L3 | L4 | young adult |
| *Cbr-lin-28-GFP*<br>*mCSD+mCCHC* | 1 | 73% (15) | 42% (12) | 28% (18) | 0% (24) | 0% (11) |
|  | 2 | 70% (20) | 18% (17) | 11% (19) | 0% (19) | 0% (5) |
|  | 3 | 40% (15) | 41% (17) | 31% (13) | 0% (28) | 0% (2) |
| *Cbr-lin-28-GFP*<br>*mCSD+mCCHC+3'-UTR del* |  | 75% (16) | 50% (14) | 35% (20) | 17% (23) | 11% (18) |

We further modified the transgene construct to contain a deletion in the 3’UTR to remove miRNA sites (Fig. S10). Animals with *aeEx48* transgene encoding a mutant protein and a 3’UTR deletion did not show any differences in the place and timing of *Cbr-lin-28* expression, except for seam cells. Fluorescing seam cells were observed less often after the L1 stage, similar to arrays with an intact 3’UTR. Nevertheless, they were observed in L4 and young adult animals in this strain, in contrast to 3 independently generated strains bearing the CSD/CCHC mutant with an intact 3’UTR (Table 1, Fig. 4B, C, and D). Thus, *Cbr-lin-28* is down-regulated in seam cells in late larval development in part via its 3’UTR.

Overall, the marked difference in expression between *C. elegans* and *C. briggsae* parallels the differences in phenotype, where it seems *Cbr-lin-28* has a broader role in the animal than *Cel-lin-28*. Nevertheless, 3’UTR-dependent down-regulation in seam cells occurs in both species.

### Cbr-lin-46(0) mutants are similar to Cel-lin-46(0) mutants

In *C. elegans, lin-46* was discovered as a suppressor of *lin-28(0)* phenotype (Pepper *et al*. 2004). *lin-46(0); lin-28(0)* double mutants appear mostly as wild type, whereas *lin-46(0)* single mutants have alae gaps and an increased number of seam cells, with both defects being cold-sensitive. *lin-46* encodes an unusual protein with protein-protein interaction activity.

A *Cbr-lin-46(0)* allele was generated by targeting the second exon: the *Cbr-lin-46(ae38)* mutation is an insertion causing a frameshift that results in a premature stop codon. A second allele, *Cbr-lin-46(ae44)*, has a deletion removing the start-codon (Table S1; Fig. S6).

The null mutant has a weak reiterative phenotype similar to *Cel-lin-46(0)* mutants. In *C. briggsae*, around 65% of the animals had alae gaps at either 20°C or 25°C, and over 80% at 15°C (Fig. 5B). The penetrance at higher temperatures is higher than is seen for *Cel-lin-46(0)* animals, but the cold sensitivity is shared (Pepper *et al*. 2004). In both species, the number of seam cells was significantly higher at 15°C (Fig. 5C). There were also slight egg-laying defects, and vulvae were often abnormally shaped in *C. briggsae* (Fig. 5A, Fig. S11). Thus, *Cbr-lin-46(0)* mutants have similar defects and cold sensitivity as *Cel-lin-46(0)* mutants, suggesting that the orthologs have similar functions. Furthermore, the fact that null alleles are cold sensitive in both species implies that a process that is exposed by the loss of *lin-46* has inherent cold sensitivity.

**Fig. 5.**
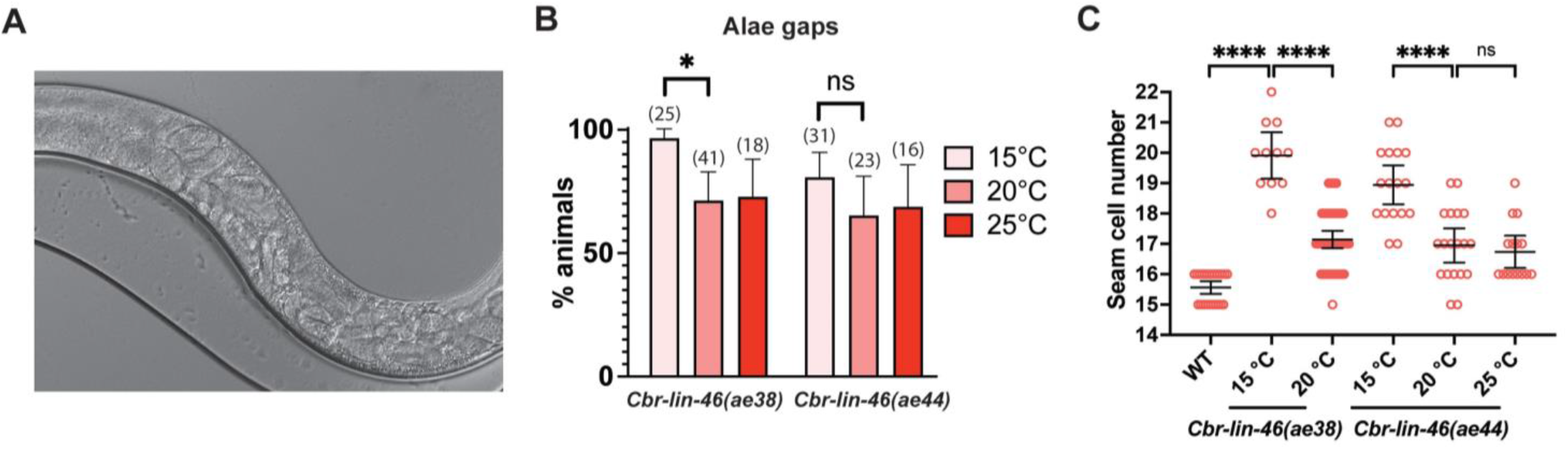
Null mutants of *Cbr-lin-46* have a reiterative phenotype. (A) A DIC photomicrograph of a *cbr-lin-46(ae38)* adult animal with an egg-laying defect. and accumulated late stage embryos at 20°C can be seen. The worm is oriented anterior end left, dorsal side up. (B) Alae gaps in *Cbr-lin-46(0)* adults are slightly more frequent at 15°C. Sample sizes are specified in parenthesis above the bars.(C) *Cbr-lin-46(0)* L4 and young adult animals have increased numbers of seam cells that are also cold-sensitive.

### Cbr-lin-46(0) partially suppresses the Cbr-lin-28(0) phenotype

To see whether the relationship between *Cbr-lin-28* and *Cbr-lin-46* is conserved despite the drift in *lin-28’s* role, we constructed a *Cbr-lin-46(ae38)*; *Cbr-lin-28(ae39)* double null mutant. Surprisingly, we found that *Cbr-lin-46(0)* suppressed not only the precocious alae defect of *Cbr-lin-28(0)* mutants, but that the L4 developmental arrest was partly suppressed (although not at 15°C), and that the gonad disorganization was partly suppressed at all temperatures (Fig. 3B and C, Fig. S9, Fig. S12).

Interestingly, *Cbr-lin-46(0)* did not suppress the L4 arrest phenotype when the double-mutant passed through dauer: the fraction of L4-arrested animals at 20°C in *Cbr-lin-28(0); Cbr-lin-46(0)* double mutants that had passed through dauer was comparable to the fraction of “arrested L4” animals in *Cbr-lin-28(0)* post-dauers at 20°C (Fig. 3B). This suggests that different downstream effectors exist for *Cbr-lin-28* in continuous development and dauer development.

We also observed some reciprocal suppression: the increased number of seam cells in *Cbr-lin-46(0)* mutants was suppressed by the *Cbr-lin-28(0)* mutation at 20°C, although not at 15°C (Fig. 3D). By contrast, successfully molted double mutants had alae gaps at comparable rates to the *Cbr-lin-46(0)* single mutants (63.1%, n = 19 at 20°C and 61.5%, n = 13 at 25°C, Fig. 5B). Thus, some of the reiterative traits of *Cbr-lin-46(0)* were not suppressed by *Cbr-lin-28(0)*, which is in contrast to what occurs in *C. elegans* (Pepper *et al*. 2004).

### The Cbr-lin-46 5’UTR mutant phenotype differs from that of a Cbr-lin-28(0) mutant

In *C. elegans*, the *lin-46* 5’UTR is a regulatory region through which *lin-28* acts to inhibit *lin-46* expression; small deletions in this sequence causes a phenotype that resembles the *Cel-lin-28(lf)* phenotype (Ilbay *et al*. 2021). This 36-nt 5’UTR is conserved among all species of *Caenorhabditis* and is identical between *C. elegans* and *C. briggsae*. We created a 6-bp deletion (allele *ae43*) in this region using CRISPR/Cas9 (Table S1, Fig. S13A). The *Cbr-lin-46* 5’UTR mutants had protruding vulvae and either full or gapped precocious alae at the L4 stage; however, they had the same number of seam cells as the wild type (Fig. S13B, C, and D). This phenotype resembles the heterochronic traits of *Cbr-lin-28(0)*, however, the penetrance and expressivity of the alae defect are more severe in the *Cbr-lin-46(gf)* mutant. Significantly, the *Cbr-lin-46* 5’UTR mutant lacks the larval-arrest and gonad disintegration defects of the *Cbr-lin-28(0)* mutants. Thus, the role of *Cbr-lin-46* in developmental timing resembles that of *Cel-lin-46*, but the fact that the phenotype of the *Cbr-lin-46* 5’UTR deletion differs substantially from that of *Cbr-lin-28(0)* suggests that the relationship between *lin-28* and *lin-46* has drifted as these species evolved.

### mir-241, mir-48, *and* mir-84 *have a conserved function in* C. elegans *and* C. briggsae

In *C. elegans*, three *let-7*-family miRNAs, *mir-48*, *mir-84*, and *mir-241* (the “*3let-7s*”), redundantly control *hbl-1* and *lin-28*: a strong phenotype appears when all 3 are knocked out, causing reiteration of L2-specific cell fates, whereas single mutants have little or no effect (Abbott *et al*. 2005; Tsialikas *et al*. 2017). By contrast, in *C. briggsae*, a *Cbr-mir-48(0)* mutation alone yielded a strong phenotype: *Cbr-mir-48(ae65)* mutant burst at the vulva at the end of the L4 molt, had an increased number of seam cells, and incomplete adult alae (Table S1, Fig. 6A and B). We saw that 95% of mutant adults had less than half of the normal amount of adult alae, and 11% lacked alae entirely (n = 44).

**Fig. 6.**
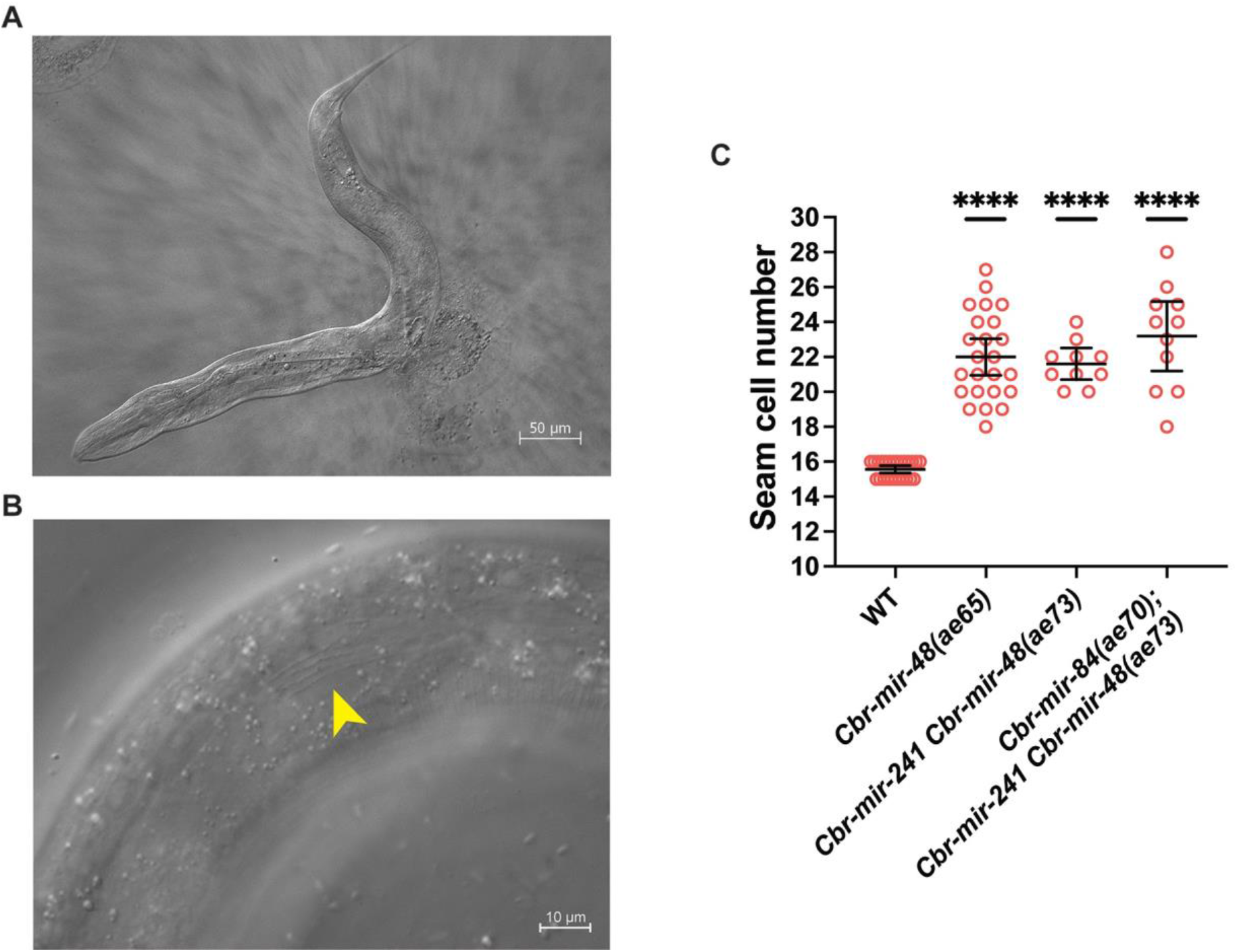
*Cbr-3let-7s* mutants have reiterative phenotypes. (A) DIC micrograph of a *Cbr-mir-48(ae65)* mutant that has burst at vulva after reaching adulthood. (B) *Cbr-mir-48(ae65)* young adults develop alae patches (indicated by an arrowhead). (C) *Cbr-mir-48(ae65)* have an increased number of seam cells, but adding *Cbr-mir-241(0)* and *Cbr-mir-84(0)* mutations does not cause further increase in the seam cell number. The difference between mutant groups was not statistically significant (unpaired Welch’s t-test, *p* > 0.05).

The *Cbr-mir-48* and *Cbr-241* genes are within 3kb of each other on linkage group V, and we obtained a deletion allele that removed both (Table S1; *ae73*). Deletion of both miRNAs resulted in a more severe phenotype than *Cbr-mir-48(ae65)* alone: 92% of adult animals lacked alae altogether (n = 13), compared to 11% for the single mutant. However, the double mutants that had developed through the dauer pathway did not burst at the vulva and appeared wild-type, indicating that these mutations, like their *C. elegans* counterparts, are suppressed by the dauer developmental pathway (data not shown).

Deleting all three miRNAs resulted in animals that could not be maintained as homozygotes because most were sterile. However, the frequency of alae patches in *Cbr-mir-48 Cbr-mir-241(ae73); Cbr-mir-84(ae70)* animals segregating from heterozygotes was similar to that of the double mutant: 92% lacked alae (n = 13). Furthermore, this triple mutant did not show an increase in the number of seam cells compared to the *Cbr-mir-48(0)* single mutant (Fig. 6C).

In other aspects, the *Cbr-mir-241(0)* and *Cbr-mir-84(0)* single null mutants and the *Cbr-mir-241(0); Cbr-mir-84(0)* double null mutants had few differences from the wild type. There were no alae gaps and the number of seam cells was close to normal at both 20°C and 15°C (Fig. S14). A few animals had egg-laying defects and abnormal vulvae (Fig. S15), and some of these egg-laying defective animals appeared to be stuck in lethargus (not pumping). Finally, a small percentage of sterile animals was observed in these strains (Table 2).

**Table 2:**
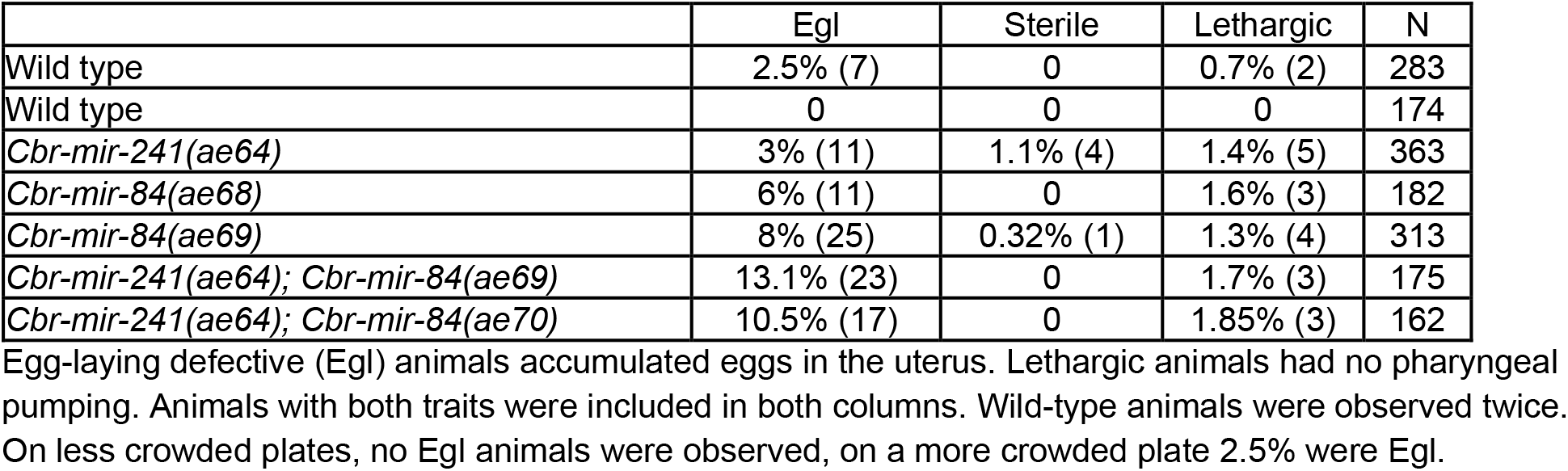
Phenotypes of *Cbr-mir-241(0)* and *Cbr-mir-84(0)* mutants.

|  | Egl | Sterile | Lethargic | N |
| --- | --- | --- | --- | --- |
| Wild type | 2.5% (7) | 0 | 0.7% (2) | 283 |
| Wild type | 0 | 0 | 0 | 174 |
| *Cbr-mir-241(ae64)* | 3% (11) | 1.1% (4) | 1.4% (5) | 363 |
| *Cbr-mir-84(ae68)* | 6% (11) | 0 | 1.6% (3) | 182 |
| *Cbr-mir-84(ae69)* | 8% (25) | 0.32% (1) | 1.3% (4) | 313 |
| *Cbr-mir-241(ae64); Cbr-mir-84(ae69)* | 13.1% (23) | 0 | 1.7% (3) | 175 |
| *Cbr-mir-241(ae64); Cbr-mir-84(ae70)* | 10.5% (17) | 0 | 1.85% (3) | 162 |

Egg-laying defective (Egl) animals accumulated eggs in the uterus. Lethargic animals had no pharyngeal pumping. Animals with both traits were included in both columns. Wild-type animals were observed twice. On less crowded plates, no Egl animals were observed, on a more crowded plate 2.5% were Egl.

In *C. elegans*, mutations in *lin-4* and the three *let-7*-related miRNAs *mir-48*, *mir-84*, and *mir-241*, have distinct phenotypes: deletion of *lin-4* causes reiteration of L1-specific fates and deletion of all of the *3let-7s* causes reiteration of L2-specific fates (Chalfie *et al*. 1981; Abbott *et al*. 2005). All four of these miRNA genes have been shown to act together in stage-specifically down-regulating *lin-28* to ensure appropriate expression of L2 fates (Tsialikas *et al*. 2017). Because we found that mutations in *Cbr-lin-4* and *Cbr-3let-7s* both cause reiteration of L2 fates, we tested whether these mutations enhanced one another, leading to a reiteration of the earlier L1 fates. To our knowledge, the equivalent mutant of *C. elegans* has not been reported. A *Cbr-lin-4* deletion was introduced into *Cbr-mir-241(0), mir-48(0), mir-84(0)* mutant background (also containing *Cbr-hbl-1::AID; TIR1(F79G)*; see below). The quadruple mutants were vulvaless with gapped alae at adulthood (data not shown). However, in contrast to *Cbr-lin-4(0)* mutants, late L2 larvae of the quadruple mutant had a lower number of seam cells (mean = 12.4+/-1.3, n = 14), suggesting that some seam cells reiterated L1 fates. The phenotype resembles the *Cbr-lin-14* 3’UTR deletion, suggesting that *Cbr-3let-7s* participate in the down-regulation of *Cbr-lin-14*, which contains let-7 sites in its 3’UTR.

### Auxin-inducible degron system in C. briggsae

In *C. elegans*, certain heterochronic genes have pleiotropic phenotypes that include embryonic lethality or infertility. Anticipating that these genes might have similar pleiotropies in *C. briggsae*, we used the auxin-inducible degron system to generate conditional alleles. We produced lines of *C. briggsae* expressing *TIR1(F79G)* from extrachromosomal (*aeEx44)* and attached (*aeIs15)* arrays (Hills-Muckey *et al*. 2021). The attached transgene (*aeIs15*) was located by crossing with marker strains and found to be on LGII.

Unexpectedly, both the extrachromosomal and attached *TIR1(F79G)* arrays caused a reduction in the number of intestinal nuclei (Table S3). However, no other abnormalities were observed and the animals appeared to develop normally and be healthy. The reason for this reduction in intestinal nuclei is unclear. The intestinal nuclei glowed brightly, indicative of high array expression.

### Cbr-hbl-1(lf) causes a precocious phenotype like Cel-hbl-1(lf)

In *C. elegans*, *hbl-1* encodes an Ikaros-family transcription factor involved in hypodermis development where null alleles are embryonic lethal and weak alleles have a heterochronic phenotype with reduced number of seam cells, precocious alae, and a protruding vulva, resembling *lin-28(0)* (Fay *et al*. 1999; Lin *et al*. 2003; Abrahante *et al*. 2003). To study *Cbr-hbl-1* loss-of-function while avoiding potential embryonic lethality, the locus was tagged with an auxin-inducible degron (AID) using CRISPR/Cas9 (Table S1). The *hbl-1::AID* strain was then crossed with lines bearing the *TIR1(F59G)* transgene to generate lines in which *Cbr-hbl-1* activity could be reduced in response to the auxin analog 5-Ph-IAA (Hills-Muckey *et al*. 2021).

Adult animals were placed on plates with 0.01-0.02 µmol of 5-Ph-IAA, and the phenotypes of the next generation were characterized. We saw that 47% of animals on those plates had fully precocious alae, and the remainder had gapped precocious alae (n = 30). There was no embryonic lethality. However, 82% of adult animals remained stuck in the L4 molt, and 27% of mutants had a precocious vulval differentiation (reaching the “christmas tree”, or L4.4-L4.5 according to Mok, *et al*., 2015, stage of morphogenesis by the end of the L3 stage) or a protruding vulva (Fig. 7B). As in *C. elegans hbl-1(lf)*, the protruding vulva developed during the L4 stage. Other animals had normal vulval development and a functional vulva. There was a slight reduction in seam cell number in some *Cbr-hbl-1(lf)* animals. However, the reduction was not as significant as in *Cel-hbl-1::AID* under similar conditions (Fig. 7C). These observations show that the functions of *hbl-1* are at least partly conserved between *C. elegans* and *C. briggsae*.

**Fig. 7.**
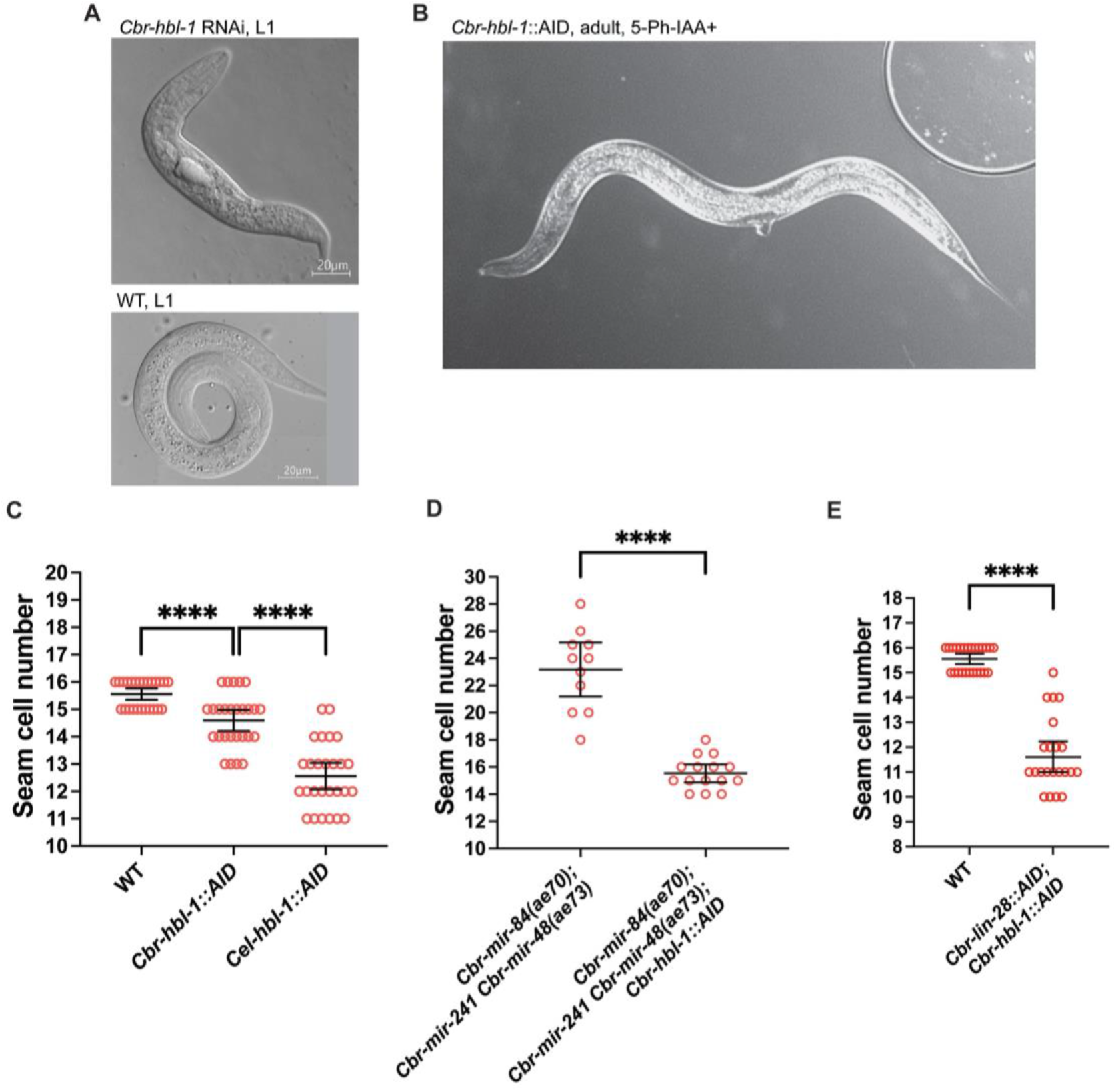
*Cbr-hbl-1* is required for early development and promotes L2 seam cell fates. (A) DIC micrograph of a wild-type L1 larva and deformed L1 larva from *Cbr-hbl-1* dsRNA injections. These animals do not survive and develop into adults. (B) A *Cbr-hbl-1::AID* young adult grown on 5-Ph-IAA. Notice a protruding vulva. (C) Some *Cbr-hbl-1::AID* mutants have a slightly reduced number of seam cells in the presence of 5-Ph-IAA, but the reduction is not as large as for *Cel-hbl-1::AID*, which may due to differences in *hbl-1* orthologs functions or a difference in the degradation efficiency. (D) *Cbr-hbl-1* depletion on 5-Ph-IAA suppresses increased seam cell numbers in the *Cbr-3let-7s* triple mutant. *Cbr-3let-7s* seam cell counts are the same data set as in Fig. 6,C. (E) Combined depletion of *Cbr-lin-28::AID* and *Cbr-hbl-1::AID* on 5-Ph-IAA causes a reduced number of seam cells.

To test whether a more drastic reduction of *Cbr-hbl-1* activity could cause embryonic lethality like that seen in *C. elegans*, we performed RNAi. A dsRNA representing 668bp from exon 4 of the *Cbr-hbl-1* ORF was injected into wildtype *C. briggsae*, and we observed dead L1 larvae in the next generation (Fig. 7A). Their terminal phenotype appeared to be more developmentally advanced than that observed in similar *C. elegans* experiments with none of the eggs failing to hatch (Fay *et al*. 1999). No older larvae or adults with heterochronic phenotypes were observed, potentially indicating that all animals receiving RNAi failed to proceed with development after hatching. This result suggests that *Cbr-hbl-1* may have a role in embryonic development but one that differs slightly from *Cel-hbl-1*.

### Reiterative phenotype of Cbr-3let-7s is suppressed by Cbr-lin-28(0) and Cbr-hbl-1(lf)

The *3let-7s* play an important role in down-regulating *lin-28* and *hbl-1* in *C. elegans*, a conclusion supported by the fact that *lin-28(0)* and *hbl-1(lf)* are epistatic to loss of the *3let-7s* (Abbott *et al*. 2005). Likewise, we found that *Cbr-lin-28(0)* and *Cbr-hbl-1(lf)* were epistatic to the reiterative phenotype caused by *Cbr-3let-7s(0)*. The *Cbr-hbl-1::AID; Cbr-3let-7s(0); TIR1(F79G)* strain had a number of seam cells close to normal (Fig. 7D), precocious alae (100%, n = 14), and protruding vulvae (43%, n = 14) when grown on 5-Ph-IAA plates. *Cbr-lin-28(ae39); Cbr-mir-48(ae65)* double mutants had a normal number of seam cells (15.8+/-0.6, n = 13), precocious alae patches (66.7%, n = 12) at L4, a nearly complete adult alae, although some had a gap (25%, n = 8), and protruding vulvae (24%, n = 21).

These observations suggest that both *Cbr-lin-28* and *Cbr-hbl-1* act downstream of the *Cbr-3let-7s* and are necessary for the reiteration of the L2 stage caused by these three mutants, as in *C. elegans*. This is surprising since *Cbr-lin-28(0)* and *Cbr-hbl-1(0)* single mutants mostly did not show a reduction in seam cell numbers.

### Simultaneous reduction of Cbr-lin-28 and Cbr-hbl-1 activities shows that Cbr-lin-28 acts in the L2

In *C. elegans*, both *lin-28* and *hbl-1* are needed for L2 fates to occur. Because *C. briggsae lin-28(0)* mutants showed no L2 defect, we investigated whether it might still be involved at this stage by testing whether mutations would enhance the precocious phenotype caused by loss of *Cbr-hbl-1* activity. To do this, we generated a *Cbr-lin-28::AID; Cbr-hbl-1::AID* strain in a *TIR1(F79G)* background. When grown on 5-Ph-IAA plates, these animals had a reduction in seam cell numbers that was more severe than *Cbr-hbl-1:aid* alone (Fig. 7E; compare with Fig. 7C). They also developed gapped alae at the L3 stage (100% of animals had some precocious alae at L3), and complete alae by the L4. Thus, the reduction of both *Cbr-lin-28* and *Cbr-hbl-1* activity resembled the *Cel-lin-28(0)* phenotype. These double mutants also had a prolonged L3 stage and became stuck in the L3 molt: 24 hours after L3 animals were selected, some still had L3 cuticles with gapped alae and non-reflexing gonads (Fig. S16). In other animals, the gonads migrated closer to the pharynx and anus than normal before reflexing, and occasionally, the distal tips cells leading the gonad arms migrated in unexpected directions. Finally, the vulvae were either protruding or stuck in an L4-like (pre-“christmas tree”, or L4.2-L4.3 according to Mok *et al*., 2015) shape. Interestingly, all of the *Cbr-lin-28::AID; Cbr-hbl-1::AID* animals grown on 5-Ph-IAA were sterile and had disorganized gonads.

These observations show that *Cbr-lin-28* is involved with *Cbr-hbl-1* in promoting L2 cell fates, as in *C. elegans*. But in *C. elegans*, both genes are necessary, and in *C. briggsae* they are partially redundant.

### *Cbr-let-7(0)* mutants have additional molts but no heterochronic defects

*C. elegans let-7(0)* mutants have delayed adult alae formation due to reiteration of L3-specific developmental events, and they burst at the vulva upon reaching adulthood (Reinhart *et al*. 2000; Vadla *et al*. 2012). Two mutant alleles of *Cbr-let-7* were generated: an insertion (*ae47*) and deletion (*ae48*), both of which eliminate *Cbr-let-7* activity (Table S1).

Both *Cbr-let-7(0)* mutants displayed egg-laying defects, slightly abnormal vulvae shapes, and at least one extra molt (Fig. S17). The vulvae of these animals were slightly protruding, and adults burst at the vulvae on microscope slides, suggesting defects in vulval development or structure. In contrast to *Cel-let-7(0)*, the *Cbr-let-7(0)* mutants developed normal adult alae at the L4 molt. The alae looked thinner than the wild type, perhaps due to a cuticle defect or the formation of overlying cuticle during an extra molt (Fig. S18). Some *Cbr-let-7(0)* mutants may have two extra molts: A plate containing four L4 larvae were placed at 20°C and the number of shed cuticles were counted the next day: 6 cuticles were found and 2 adult animals (with eggs) were stuck in cuticles (Fig. S19). The *Cbr-let-7(0)* mutants had a slightly increased number of seam cells at low temperatures, but it is unclear whether this is a heterochronic defect (Fig. S20).

### Cbr-let-7(0) *mutation suppresses later defects of* Cbr-lin-14(0) *and* Cbr-lin-28(0) mutants

Despite *Cbr-let-7(0)* mutants not reiterating late larval stage seam cell fates (as assessed by alae formation), we tested whether they could nevertheless suppress the precocious alae defect of *Cbr-lin-28(0)*, as occurs in *C. elegans* (Slack *et al*. 2000; Vadla *et al*. 2012). Examining *Cbr-lin-28(ae39); Cbr-let-7(ae47)* animals, we found that *Cbr-let-7(0)* mutation completely suppressed the “arrested L4” phenotype and gonad disorganization of the *Cbr-lin-28(0)* mutant, which implies that this aspect of the *Cbr-lin-28(0)* phenotype is in part due to the inappropriate upregulation of *Cbr-let-7* and presumably the subsequent silencing of the miRNA’s targets.

Some *Cbr-lin-28(0); Cbr-let-7(0)* animals looked wild-type, whereas others had egg-laying defects and resembled *Cbr-let-7(0)* mutants. The double mutants had slightly abnormal vulvae similar to *Cbr-let-7(0)* single mutants, and sometimes developed protruding vulvae (data not shown). The double mutants also underwent extra molts, however, unlike the *Cbr-let-7(0)* single mutants, they usually did not complete those molts and remained stuck in cuticles (Fig. S21). Four L4 larvae from the *Cbr-lin-28(0); Cbr-let-7(0)* strain were isolated on a separate plate, and cuticles were scored next day; 4 cuticles were found and all of the animals were adults stuck in the cuticle while molting. Thus, loss of *Cbr-lin-28* slightly mitigates this *Cbr-let-7(0)* phenotype, suggesting that some functions of *Cbr-let-7* depend on the activity of *Cbr-lin-28*.

Interestingly, although some *Cbr-lin-28(0); Cbr-let-7(0)* double mutants developed precocious alae at the L4 stage at both 15°C (84.2%, n = 38) and at 20°C (25%, n = 8), the frequencies were lower than in *Cbr-lin-28(0)* single mutant (compare to Fig. 3C). Alae patches were usually located on the head and just behind the pharynx, but short patches were also observed in other areas. The results suggest that precocious alae formation in *Cbr-lin-28(0)* mutants is in part caused by premature *Cbr-let-7* upregulation.

Finally, we examined a *Cbr-lin-14(ae51) Cbr-let-7(ae50)* double mutant. The *Cbr-let-7(0)* mutation restored fertility to *Cbr-lin-14(0)* mutants and suppressed precocious alae formation. This observation suggests that sterility and precocious alae occur in *Cbr-lin-14(0)* mutants because of the inappropriate *Cbr-let-7* expression. In *C. elegans*, *lin-14* acts in part through *lin-28* to control late-stage events, so the sterility and precocious alae of *Cbr-lin-14(0)* could be due to the down-regulation of *Cbr-lin-28*, although we have not tested that hypothesis here (Seggerson *et al*. 2002; Tsialikas *et al*. 2017).

### The Cbr-hbl-1(lf) phenotype is partly epistatic to that of Cbr-let-7(0)

Work in *C. elegans* suggests that *hbl-1* acts downstream of *let-7* (Lin *et al*. 2003; Abrahante *et al*. 2003; Abbott *et al*. 2005; Vadla *et al*. 2012). We tested the ability of *Cbr-hbl-1::AID* to suppress *Cbr-let-7(0)* phenotype by constructing the double mutant. Of double mutant animals grown on 5-Ph-IAA plates, 81% (n = 21) developed patches of precocious alae at the L3 molt, although no animals were stuck in the L4 molt. These results indicate that the *Cbr-hbl-1(lf)* mutant phenotype is partly epistatic to that of *Cbr-let-7(0)*, as seen in *C. elegans* (Abrahante *et al*. 2003).

### A Cbr-lin-41(lf) mutation causes a developmental arrest at the L4 stage

*C. elegans lin-41* has multiple roles in the animal: null mutants are sterile (due to a germline defect) and hypomorphs have a heterochronic phenotype (Slack *et al*. 2000; Tocchini *et al*. 2014). We chose to tag the *C. briggsae* ortholog with the auxin-inducible degron so its level of activity could be controlled. We inserted the *aid* sequence near the start codon of *Cbr-lin-41* with CRISPR/Cas9. In this process, we also generated two loss-of-function alleles, including a potential null allele (Table S1). This allele, *Cbr-lin-41(ae77),* is an incorrect *aid* insertion creating a false ORF that did not contain any *Cbr-lin-41* exons. A potential start codon that was in-frame with the remaining *Cbr-lin-41* ORF was located 181 bp downstream of the original start. Because homozygous *Cbr-lin-41(ae77)* animals were sterile, the allele was balanced with the marker *Cbr-spe-8(v142)* that is also sterile when homozygous (R. Ellis, pers. comm.). Thus, only heterozygous animals reproduced. *Cbr-lin-41(ae77)* segregating from the heterozygous strain were identified as having a shorter body (Dpy-like phenotype).

Homozygous *Cbr-lin-41(ae77)* animals had a developmental delay at the end of the L4 stage similar to that observed in *Cbr-lin-28(0)* mutants. Around 52 hours post hatching, when wild-type animals complete the L4 molt, develop alae, adult vulva, and both types of gametes, the *Cbr-lin-41(ae77)* animals also had oocytes and spermatozoa but did not have alae, and their vulvae were stuck at the pre-“christmas tree” stage of morphogenesis (Fig. 8). Sometimes they had disorganized gonads similar to *Cbr-lin-28(0)* mutants. However, older *Cbr-lin-41(ae77)* animals had alae and adult-shaped vulvae in contrast to *Cbr-lin-28(0)* mutants, which sometimes failed to continue development after the L4 arrest. This suggests that the “arrested L4” state in *Cbr-lin-28(0)* mutants cannot be due solely to *Cbr-lin-41* down-regulation. Notably, in contrast to *Cel-lin-41* mutants, *Cbr-lin-41(ae77)* animals did not have precocious alae.

**Fig. 8.**
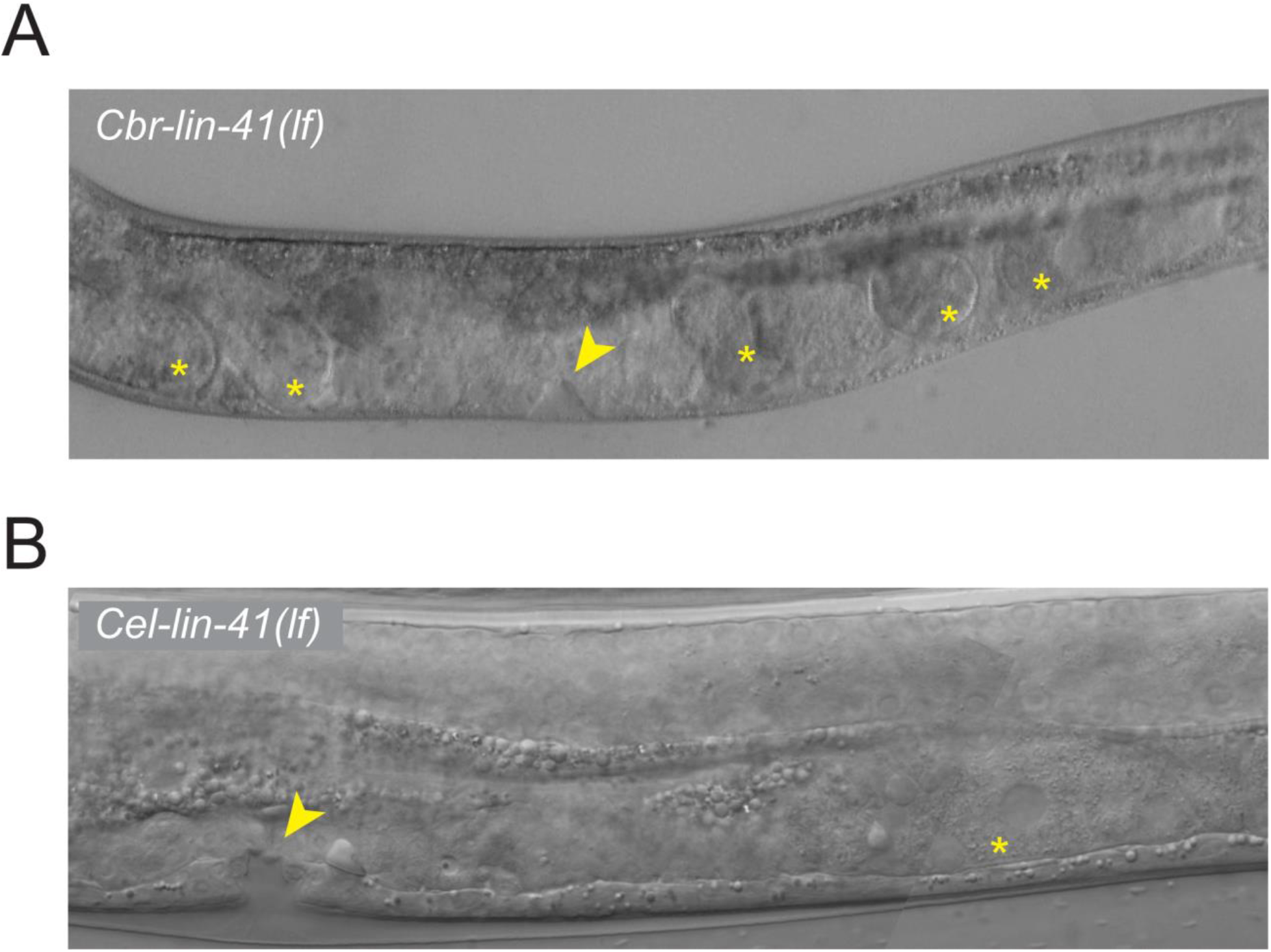
Both *Cel-lin-41(lf)* and *Cbr-lin-41(lf)* have L4 developmental delay or arrest. DIC micrographs of (top) *Cbr-lin-41(ae76)* and (bottom) *Cel-lin-41::AID(aeIs10)* animals grown on 5-Ph-IAA. In both, the vulva is arrested at an L4 stage of development (arrowheads) and the germ line is producing embryos or oocytes (asterisks). Animals are oriented anterior end left, dorsal side up.

The *Cbr-lin-41(ae76)* allele is a small insertion that creates a frameshift and early stop codon. Another in-frame start codon occurred 170 bp downstream of the original, which might allow some expression of functional protein (Table S1). This allele caused a weaker phenotype than *ae77* and could be maintained as a homozygous line. The *Cbr-lin-41(ae76)* mutants had these additional defects: 17.9% successfully molted into adults, 67.9% were stuck in the L4 molt, and 14.2% had an “arrested L4” phenotype with disorganized gonads (n = 28). None had precocious alae.

Animals with a degron-tagged *Cbr-lin-41* and *TIR1(F79G)* expressed from an attached array grown on 5-Ph-IAA produced oocytes and sperm before the soma completed development: the animals had vulvae still undergoing morphogenesis and no alae. The *Cbr-lin-41::AID* animals also had slightly abnormal vulvae, with asymmetric shapes and protrusions, and had egg-laying defects. There were no sterile animals on 5-Ph-IAA, and no precocious alae. The observations reveal similar drifts in the roles of *lin-28* and *lin-41* during the evolution of the two species, with more explicit control of developmental timing per se in *C. elegans* and less in *C. briggsae*.

### Depletion of C. elegans lin-41 causes L4 developmental arrest

Although a variety of defects caused by loss of *Cel-lin-41* activity have been observed (Slack *et al*. 2000; Tocchini *et al*. 2014), to our knowledge, L4 arrest like we observed in *Cbr-lin-41(lf)* mutants, have not been reported. To test whether *C. elegans lin-41(lf)* mutants can arrest in the L4 stage, we generated a *Cel-lin-41::AID* strain and crossed it with a strain expressing *TIR1(F79G)* allele (Hills-Muckey *et al*. 2021). Synchronized L1 larvae of wild-type *C. elegans* and *Cel-lin-41::AID* were placed on plates with or without 5-Ph-IAA. After 57 hours of development, 71% (n = 14) of wild-type animals were molting or had already shed cuticles, and the remainder were still late L4 larvae. Most molting animals had mature vulvae and were still producing sperm. Only 14% of the animals were already producing oocytes (one young adult and one molting animal). Of the *Cel-lin-41::AID* animals, 40% (n = 15) were molting, only 13% had mature vulvae (2 of the molting animals), and the others had L4-shaped vulvae. Surprisingly, 73% of the animals already had both sperm and oocytes, including animals that appeared like “late L4” larvae. Moreover, when we looked at *Cel-lin-41::AID* animals on 5-Ph-IAA at 72 hours of development, 13% of animals had an arrested L4 phenotype characterized by an L4-like vulva (“christmas tree”-like morphology), lack of adult alae, the absence of the final molt, and the presence of oocytes (Fig. 8). These observations indicate that reduction of *Cel-lin-41* activity can produce an L4 developmental delay or arrest similar to that observed for both *Cbr-lin-41(lf)* and *Cbr-lin-28(0),* resulting in asynchrony between the germline and soma.

### The Cbr-lin-41 gene acts downstream of Cbr-let-7 in the heterochronic pathway

In *C. elegans*, *lin-41* is a primary target of *let-7* in the heterochronic pathway (Slack *et al*. 2000). To test whether *Cbr-lin-41* acts downstream of *Cbr-let-7*, we made the *Cbr-lin-41(ae76)*; *Cbr-let-7(ae48)* double mutant. Of the double mutants observed (n = 33), 97% of the animals were stuck in the L4 molt and 3% had an “arrested L4” phenotype (L4-shaped vulva, no alae, produced embryos). The difference in the penetrance of the “arrested L4” phenotype was not significant compared to the *Cbr-lin-41(ae76)* single mutant (*p* > 0.05, Fisher’s exact test). Because the double mutant displayed the *Cbr-lin-41(lf)* mutant characteristics and none of the *Cbr-let-7(0)* mutant features were displayed, the *Cbr-lin-41(lf)* phenotype was epistatic to that of *Cbr-let-7(0)*, suggesting that *Cbr-lin-41* acts downstream of *Cbr-let-7*, as in *C. elegans*.

### A Cbr-lin-41(0) mutation enhances the Cbr-hbl-1(lf) phenotype

In *C. elegans*, *lin-41* and *hbl-1* both appear to control L3 cell fates, and the animals lacking the activities of both genes generate some precocious alae at the L2 molt (Abrahante *et al*. 2003; Vadla *et al*. 2012). To test whether depletion of both these genes would cause a more severe phenotype in *C. briggsae*, a *Cbr-lin-41(ae76); Cbr-hbl-1::AID* double mutant was generated. Double mutants carrying the *TIR1(F79G)* unattached array were placed on 5-Ph-IAA-containing plates, and their phenotypes were analyzed. Double mutants resembled *Cbr-lin-28::AID; Cbr-hbl-1::AID* animals grown on 5-Ph-IAA: they had an L3 developmental delay, gonads with delayed reflexion, and abnormal DTC migration after the reflection. Moreover, precocious alae patches appeared at the L2 molt in 69% of animals (n = 26), and 100% of animals had gapped (68.4%) or complete (31.6%) precocious alae at the L3 molt (n = 19). The double mutants also failed to shed L3 cuticles. The number of seam cells was slightly reduced (14.5+/-0.8, n = 26) compared to wild type (15.6+/-0.5, n = 25), but was similar to that observed in *Cbr-hbl-1::AID* strain grown on 5-Ph-IAA (14.8+/-0.8, n = 23, Fig. 7C). Thus, these counts do not show an effect of *Cbr-lin-41(lf)* on seam cell number. Our observations suggest that *Cbr-lin-41* and *Cbr-hbl-1* also control L3 hypodermal cell fates redundantly, but the fact that seam cell duplication occurs normally shows that *lin-41* does not control L2 fates.

### The Cbr-lin-29(0) phenotype resembles the Cel-lin-29(0) phenotype

The *lin-29* gene encodes a zinc finger transcription factor that directly regulates the larval-to-adult adult switch in the *C. elegans* hypodermis; in its absence, the seam cells fail to differentiate, whereas precocious alae are formed because of early *lin-29* activity (Ambros and Horvitz 1984; Ambros 1989; Rougvie and Ambros 1995; Azzi *et al*. 2020).

A *Cbr-lin-29(0)* mutant allele was made by targeting the 6th exon, where orthologous *Cel-lin-29* mutations are located (Rougvie and Ambros 1995). The deletion *ae75* is a frameshift which leads to a premature stop codon (Table S1). The *Cbr-lin-29(ae75)* mutants did not develop alae and had delayed vulval development that caused them to burst at the adult stage (n = 15, Fig. S22). Some animals retained a patch of L4 cuticle around the vulva, which occasionally prevented bursting and allowed some animals to survive and produce eggs. *Cbr-lin-29(0)* rarely produced larvae and could not be maintained as homozygotes. We therefore balanced *Cbr-lin-29(ae75)* with *Cbr-trr-1(v76)* (Guo *et al*. 2013). Overall, the phenotypes of *Cel-lin-29(0)* and *Cbr-lin-29(0)* mutants are very similar, suggesting conserved function and relationship to targets.

### Cbr-lin-28 *and* Cbr-lin-41 *act through* Cbr-lin-29

In *C. elegans*, *lin-29* acts at the end of the heterochronic gene hierarchy and is necessary for the late-stage phenotypes of earlier acting heterochronic genes (Ambros 1989). To test whether the L4 developmental arrest in particular of *Cbr-lin-28(0)* and *Cbr-lin-41(lf)* mutants required *Cbr-lin-29* activity, we made the *Cbr-lin-28(ae39); Cbr-lin-29(ae75)* and *Cbr-lin-41(ae76); Cbr-lin-29(ae75)* double mutants.

Double homozygotes of these alleles could not be maintained because they burst at the vulva at adulthood and had very few progeny. Strains that were heterozygous for *Cbr-lin-29(ae75)* and homozygous for *Cbr-lin-28(ae39) or Cbr-lin-41(ae76)* (determined by PCR genotyping) segregated mostly worms (more than 50%) that lacked developmental arrest and disorganized gonads, suggesting that loss of one copy of *Cbr-lin-29* is sufficient to suppress these phenotypes. These strains segregated worms phenotypically similar to *Cbr-lin-29(0),* which lacked adult alae and burst at the vulva (Fig. S23), as well as worms that phenotypically resembled *Cbr-lin-41(lf)* or *Cbr-lin-28(0)* respectively. Among *Cbr-lin-41(ae76); Cbr-lin-29(ae75)/+* animals, 60% of worms resembled wild type (a normal L4 molt, vulva and adult alae), 19% were stuck in the L4 molt and looked like *Cbr-lin-41(ae76)* single mutants, and 21% looked like *Cbr-lin-29(ae75)* single mutants (n = 48). *Cbr-lin-28(ae39); Cbr-lin-29(ae75)/+* animals segregated L4 larvae that had precocious alae patches (12 of 14 animals examined), and adults that had the *Cbr-lin-29(0)* phenotype (burst vulvae) lacked alae completely (n = 11). These observations suggest that loss of *Cbr-lin-29* is epistatic to loss of either *Cbr-lin-28* or *Cbr-lin-41*. Thus, *Cbr-lin-28* and *Cbr-lin-41* act through *Cbr-lin-29*, as they do in *C. elegans*, and interestingly, the arrested L4 phenotype also depends on *Cbr-lin-29*.

## Discussion

We characterized 11 *C. briggsae* orthologs of *C. elegans* heterochronic genes using a total of 35 genetic lesions and 18 double and triple mutants and found that their mutant phenotypes differ in significant ways from those of *C. elegans*. Although most orthologs displayed defects in developmental timing, some of the phenotypes differed in which stages were affected, the penetrance and expressivity of the phenotypes, or by having pleiotropic effects that were not obviously connected to developmental timing. However, when examining pairwise epistasis and synergistic relationships, we found those reflected the relationships between their *C. elegans* orthologs, suggesting that the arrangements of these genes in functional modules is conserved, but the modules’ relationships to each other and/or to their targets has drifted since the time of the species’ last common ancestor.

A previous comparison of *C. elegans* and *C. briggsae* orthologs by RNA-interference (RNAi) showed that only a small fraction (91 of 1333 orthologs) have significantly different loss-of-function phenotypes in the two species (Verster *et al*. 2014). This study included the protein-coding heterochronic gene orthologs, except *Cbr-lin-28*, but in general did not detect the degree of divergence that we observed. However, the level of analysis was limited and included only larval or embryonic lethality, growth rate, morphology, and fertility.

Our observations suggest that the level of functional divergence in the heterochronic gene orthologs is greater than previously thought. Despite this, the fundamental structure of the heterochronic pathway is largely conserved between *C. elegans* and *C. briggsae*. Since these species have nearly indistinguishable larval development and post-embryonic cell lineages, the differences in single-gene mutant phenotypes and some pairwise relationships indicate a significant degree of developmental systems drift has occurred while giving rise to essentially the same anatomy and life history.

### Conservation of key regulatory modules

As has been done in the analysis of other complex developmental pathways, the heterochronic pathway can be divided into subcircuits or modules that govern individual aspects of the phenotype (Fig. S24, Verd *et al*. 2019). The modules are defined by enhancement or epistasis relationships among genes that act either together or in opposition to control cell fates at specific larval stages. Our findings suggest that these modules are largely conserved between *C. elegans* and *C. briggsae*.

The *lin-4/lin-14* module in *C. elegans* specifies L1 cell fates and controls the transition to L2 fates (Chalfie *et al*. 1981; Ambros and Horvitz 1987; Wightman *et al*. 1993). *lin-14* acts to specify L1 cell fates then is down-regulated by *lin-4*, allowing L2 and later fates to occur. We found the same is true in *C. briggsae*: *Cbr-lin-14* has nearly the same role and is required for the reiterative mutant phenotype of mutant *Cbr-lin-4*. Taken with the fact that the miRNA sites in *Cbr-lin-14*’s 3’UTR are conserved, and their deletion also causes a reiterative phenotype, we find that the regulatory relationship between *lin-4* and *lin-14* is conserved between the two species.

The genes *lin-28, lin-46*, *3let-7s*, and *hbl-1* comprise a complex regulatory module that, in *C. elegans*, specifies L2 cell fates and the transition to the L3 (Pepper *et al*. 2004; Vadla *et al*. 2012; Tsialikas *et al*. 2017; Ilbay and Ambros 2019; Ilbay *et al*. 2021). In *C. briggsae*, the relationships among these genes is conserved, although relative roles within the module have drifted. As in *C. elegans*, in *C. briggsae* both *lin-28* and *hbl-1* are required for L2 fates, *lin-46* acts downstream of *lin-28* while *lin-28* also has *lin-46*-independent activity, the *3let-7*s act upstream of *hbl-1*, and the *3let-7*s are needed for the transition to L3 fates.

The transition from L3 to L4 and subsequently to adult cell fates is regulated by the *let-7/lin-41/lin-29* module in *C. elegans* (Slack *et al*. 2000; Vadla *et al*. 2012; Azzi *et al*. 2020). Like *lin-28*, *let-7*’s individual role has drifted, but its relationship to other genes is largely conserved: In both species, *let-7* generally acts downstream of *lin-14*, *lin-28*, and *hbl-1* and upstream of *lin-41*. *lin-41*’s role has also drifted, but it appears to be a regulatory target of *let-7* and acts to negatively regulate *lin-29* in both species. *lin-29* is the furthest downstream of all the genes in both species, at least with regard to the larva-adult transition. Thus, the core relationships in the *let-7/lin-41/lin-29* regulatory module are conserved.

It is also worth noting the high phenotypic variability that we observed, which is greater in *C. briggsae* than in *C. elegans* for some mutants. This fact speaks to the high degree of genetic buffering that these modules provide when fully intact, a condition that would accommodate considerable developmental systems drift.

### Evolutionary drift in regulatory relationships

Despite the fact that key regulatory relationships are conserved between the species, the single-gene phenotypes and some double mutant effects reveal substantial drift in the relationships of these genes to downstream targets. This drift is manifested in two ways: a shift in the role of the gene in the developmental timing of cell fates (their heterochronic roles), and the uncovering of additional roles that are not obviously related to cell fate timing, specifically the completion of larval development and gonad integrity.

Two aspects of the role of the *lin-4*/*lin-14* module in *C. briggsae* compared to *C. elegans* are significant. First, the fact that *Cbr-lin-4* primarily affects the transition from L2 to L3 suggests either that it alone is not sufficient to repress *Cbr-lin-14*, or that *Cbr-lin-14* is not sufficient to specify L1 fates. Second, the fact that *Cbr-lin-4(0)* suppresses—but is not epistatic to—the precocious *Cbr-lin-14(0)* indicates drift in *lin-4*’s relationship to its targets. Specifically, whereas in *C. elegans lin-4*’s primary target is *lin-14* and secondarily the L2 regulators *lin-28* and *hbl-1*, in *C. briggsae lin-4* may play a more significant role regulating *lin-28* and *hbl-1* than it does with *lin-14*.

In both species, both *lin-14* and *lin-28* have sites for *lin-4* and *let-7* family miRNAs in their 3’UTRs. The regulation of *lin-28* by both miRNA families was described previously and it was shown that the *lin-4/lin-14* module controls the expression of the *3let-7s* (Tsialikas *et al*. 2017). The drift we see in the role of *lin-4* may reflect differences in the relative strengths of multiple miRNAs from the *lin-4* and *let-7* families in down-regulating their multiple targets. Developmental systems drift arises due to the accumulation of small changes—in this case, perhaps, in miRNA expression, abundance, or target sensitivity—and how those changes are compensated for by other small changes that keep developmental outcomes the same (True and Haag 2001).

Such subtle shifts in relative strengths is more apparent among the *3let-7*s. In *C. elegans*, mutations in *mir-48, mir-84*, and *mir-241* individually have very weak or undetectable phenotypes but when combined cause a strong L2 reiterative defect (Li *et al*. 2005; Abbott *et al*. 2005). These miRNAs all resemble *let-7* in having the same 8-nt seed sequence and therefore have the potential to regulate the same targets (Lau *et al*. 2001; Lim *et al*. 2003). In *C. elegans*, *mir-48* is largely redundant with the other two genes, but in *C. briggsae mir-48* is mostly responsible for work of all three genes for this phenotype (loss of *mir-48* has nearly the same heterochronic effect as that of the triple mutant). Thus, small evolutionary changes in the relative levels of these miRNAs, or differences in the regulation caused by changes in 3’UTRs of their target genes, may be offset by compensatory changes in other miRNAs or target sites, leading to identical outputs of the regulatory module for the different species.

A reciprocal example of drift involving redundancy is *lin-28* and *hbl-1*. In *C. elegans*, each gene is needed to specify L2 cell fates (Ambros and Horvitz 1984; Lin *et al*. 2003; Abrahante *et al*. 2003). By our observations, the regulatory module involving these genes is conserved, but in *C. briggsae*, *lin-28*’s role in specifying L2 fates is barely detectable until *Cbr-lin-28(lf)* is combined with *Cbr-hbl-1(lf)*, indicating its redundancy with *Cbr-hbl-1*. The substantial difference in *lin-28*’s solo role in the two species may reflect a drift in two parallel components of this regulatory module, which are the negative regulators of *hbl-1*: the *3let-7*s and *lin-28’s* direct target, *lin-46*. In *C. elegans*, the effect of deleting *lin-46*’s 5’UTR is similar to, although weaker than, *lin-28*’s null phenotype (Ilbay *et al*. 2021). It is therefore surprising that deletion of the presumed *Cbr-lin-28* regulatory site in the 5’UTR of *Cbr-lin-46* does not phenocopy *Cbr-lin-28(0)* at all, but rather looks more like *Cbr-hbl-1(lf)*. Furthermore, the role of *lin-46* appears not to have drifted during the divergence of the two species. It is possible that *Cbr-lin-28* may not be the only gene acting via the 5’UTR of *Cbr-lin-46*, but that alone would not explain *Cbr-lin-28*’s redundancy with *Cbr-hbl-1*. A shift in the relative strength of *lin-28*’s regulation of *let-7* or another target may be responsible.

Significant drift was also found in the *let-7/lin-41/lin-29* module. On their own, neither *Cbr-let-7* nor *Cbr-lin-41* influence the developmental timing of cell fates, in striking contrast to what happens in *C. elegans*. However as in *C. elegans*, in *C. briggsae*, *lin-41* shows some redundancy with *hbl-1* in controlling seam cell differentiation, demonstrating that its regulatory relationships with other heterochronic regulators is conserved. *let-7*, on the other hand, does not have the same suppressor interactions with the early-acting heterochronic genes *lin-14* and *lin-28* in *C. briggsae* as it has in *C. elegans*, although it does have a role in cessation of the molting cycle in both species. This “split” phenotype is consistent with the analysis of Azzi and colleagues who showed a branching of the pathway in the control of *lin-29* isoforms that affect different aspects of terminal differentiation in the hypodermis (Azzi *et al*. 2020).

By far, the most significant difference between *C. elegans* and *C. briggsae* is *lin-28*’s phenotype, which shows only minor heterochronic defects in *C. briggsae* while at the same time displaying significant late larval arrest and gonad integrity problems, two unexpected phenomena that are not well understood even in *C. elegans*. This was surprising given *lin-28*’s broad conservation and role in timing among animals (Moss and Tang 2003; Balzer *et al*. 2010; Romer-Seibert *et al*. 2019). We have demonstrated here that a reduction in *lin-41* activity can also lead to L4 arrest in *C. elegans*. Perhaps a difference in *lin-28*’s regulatory relationship with *lin-41* (possibly via its direct regulation of *let-7*) is responsible for *lin-28*’s different influence on the completion of larval development in the two species. Additionally, we found significant differences in the time and place of *lin-28*’s expression between the two species—including different degrees of significance of 3’UTR regulation—which may also account for the drift in pleiotropies over evolutionary time.

Our comparison of the heterochronic genes of *C. elegans* to those of their orthologs in *C. briggsae* revealed less drastic changes than has been seen in the sex determination pathways. Sex determination pathways evolve rapidly: *C. elegans* and *C. briggsae* developed hermaphroditism independently, since their common ancestor was dioecious (Ellis 2017; Haag *et al*. 2018). Some sex determination genes have conserved roles in *Caenorhabditis* species, such as *tra-1*, *tra-2*, the *fem* genes, and *fog-3* whereas others have quite different functions, including genes involved in sex determination in one species but not the other. Whether there exist genes in *C. briggsae* that have primary roles in developmental timing but are not among the orthologs studied here can be determined by forward genetic screens in these species for mutants with developmental timing defects.

### Insights into the heterochronic pathway of *C. elegans*

Our investigation of the heterochronic gene orthologs of *C. briggsae* revealed new relationships between this pathway and other aspects of the animal’s growth that may be relevant to both species. In particular, we found that a *Cel-lin-41* mutation can cause a developmental arrest similar to that caused by *Cbr-lin-28* and *Cbr-lin-41* mutations. It is possible that redundancy that has not yet been uncovered in *C. elegans* could connect the heterochronic genes to completion of larval development and gonad integrity. Given the number of heterochronic gene orthologs directly involved gonad integrity—either because mutations in them cause disintegration (*Cbr-lin-28*, *Cbr-hbl-1*, *Cbr-lin-41*) or mutations suppress that disintegration (*Cbr-lin-46*, *Cbr-let-7*)—it would not be surprising to find that further investigation of heterochronic genes in *C. elegans* uncovers such a connection.

The majority of developmental systems drift in the heterochronic pathway appears to have occurred in redundant and parallel components of regulatory modules. Parallel regulatory branches may have different “weights” in determining phenotypic outcome in different species. Even in *C. elegans*, we do not yet fully understand the nature of these parallel branches—such as non-*lin-4* regulation of *lin-14*, *lin-46*-independent regulation of *hbl-1* by *lin-28*, and the different contributions of the three *let-7* miRNAs to the repression of multiple heterochronic genes. Further investigation using multiple *Caenorhabditis* species may reveal why the pathway is organized as it is and why it has evolved the ways it has.

## Materials and Methods

### Sequence analysis

*C. briggsae* orthologs of *C. elegans* heterochronic genes were identified previously in Wormbase (wormbase.org) and miRBase (mirbase.org) and were confirmed by reciprocal BLAST on each genome. Links to the database entry for each gene are given in Table S1.

### Strains and culture conditions

Nematodes were grown at 20°C on standard NGM plates seeded with *E. coli* AMA1004 unless otherwise indicated.

### Strains used

*C. elegans* strains:

RG733 (*wIs78* [pDP#MM016B (*unc-119*) + pJS191 (*ajm-1::GFP* + pMF1(*scm::GFP*) + F58E10]) (wild type for this study), HML1029 *cshIs140[rps-28pro::TIR1(F79G)_P2A mCherry-His-11; Cbr-unc-119(+)] LGII,* ME502 *cshIs140[rps-28pro::TIR1(F79G)_P2A mCherry-His-11; Cbr-unc-119(+)] LGII; hbl-1(aeIs8[hbl-1::AID]); wIs78,* ME504 *cshIs140[rps-28pro::TIR1(F79G)_P2A mCherry-His-11; Cbr-unc-119(+)] LGII; lin-41(aeIs10[lin-41::AID]); wIs78*, ME507 *cshIs140[rps-28pro::TIR1(F79G)_P2A mCherry-His-11; Cbr-unc-119(+)] LGII; lin-14(aeIs5[lin-14::AID]); wIs78*.

C. briggsae strains:

AF16 (wild type),

ME421 *Cbr-lin-28(ae25),*

ME444 *Cbr-dpy-5(v234) +/+ Cbr-lin-28(ae35),*

ME449 *Cbr-lin-46(ae38),*

ME450 *Cbr-lin-28(ae39),*

ME451 *Cbr-lin-46(ae38); Cbr-lin-28(ae39),*

ME454 *Cbr-dpy-5 (v234) +/+ Cbr-lin-28(ae39),*

ME480 *Cbr-lin-46(ae43),*

ME482 *Cbr-lin-46(ae44),*

ME486 *Cbr-let-7(ae47),*

ME487 *Cbr-let-7(ae48),*

ME489 *Cbr-lin-14(ae51) Cbr-let-7(ae50),*

ME493 *Cel-lin-4(ae53),*

ME494 *Cbr-lin-28(ae39); Cbr-let-7(ae47),*

ME497 *Cbr-lin-4(ae54),*

ME500 *Cbr-lin-4(ae55),*

ME511 *Cbr-hbl-1(aeIs12[Cbr-hbl-1::AID]),*

ME514 *Cbr-lin-14(ae58),*

ME515 *Cbr-lin-14(ae59),*

ME519 *Cbr-dpy-8(v262) + Cbr-unc-7(v271)/+ Cbr-lin-14(ae62) Cbr-unc-7(v271),*

ME520 *Cbr-dpy-8(v262) + Cbr-unc-7(v271)/+ Cbr-lin-14(ae63) Cbr-unc-7(v271),*

ME526 *Cbr-mir-241(ae64),*

ME527 *Cbr-mir-48(ae65),*

ME529 *Cbr-mir-84(ae68),*

ME530 *Cbr-mir-84(ae69),*

ME531 *Cbr-mir-241(ae64); Cbr-mir-84(ae70),*

ME533 *Cbr-mir-241(ae64); Cbr-mir-84(ae69),*

ME534 *Cbr-lin-4(ae71); Cbr-dpy-8(v262) + Cbr-unc-7(v271)/+ Cbr-lin-14(ae62) Cbr-unc-7(v271),*

ME535 *Cbr-lin-4(ae72); Cbr-dpy-8(v262) + Cbr-unc-7(v271)/+ Cbr-lin-14(ae62) Cbr-unc-7(v271),*

ME538 *Cbr-mir-241 Cbr-mir-48(ae73),*

ME541 *Cbr-mir-241 Cbr-mir-48(ae73)/+; Cbr-mir-84(ae70),*

ME544 *Cbr-lin-41(ae76),*

ME545 *Cbr-spe-8 (v142) +/+ Cbr-lin-41(ae77),*

ME547 *Cbr-lin-41(aeIs14[Cbr-lin-41::AID]),*

ME548 *Cbr-trr-1(v76) +/+ Cbr-lin-29(ae75),*

ME549 *aeIs15[rps-28pro>atTIR1(F79G)::P2A::GFP::His-11; Cbr-unc-119(+)]; Cbr-hbl-1(aeIs12[Cbr-hbl-1::AID]),*

ME550 *aeIs15[rps-28pro>atTIR1(F79G)::P2A::GFP::His-11; Cbr-unc-119(+)],*

ME552 *aeIs15[rps-28pro>atTIR1(F79G)::P2A::GFP::His-11; Cbr-unc-119(+)]; Cbr-lin-41(aeIs14[Cbr-lin-41::AID]),*

ME553 *aeIs15[rps-28pro>atTIR1(F79G)::P2A::GFP::His-11; Cbr-unc-119(+)]; Cbr-mir-241 Cbr-mir-48(ae73); Cbr-hbl-1(aeIs12[Cbr-hbl-1::AID]) Cbr-mir-84(ae70),*

ME554 *aeIs15[rps-28pro>atTIR1(F79G)::P2A::GFP::His-11; Cbr-unc-119(+)]; Cbr-lin-28(aeIs13[Cbr-lin-28::AID]); Cbr-hbl-1(aeIs12[Cbr-hbl-1::AID]),*

ME555 *aeEx44[rps-28pro>atTIR1(F79G)::P2A::GFP::His-11; Cbr-unc-119(+)],*

ME556 *aeEx45[Cbr-lin-28::GFP],*

ME558 *Cbr-lin-41(ae76); Cbr-let-7(ae48),*

ME559 *aeEx46[Cbr-lin-28::GFP(Y35R F37A C127R C137A)],*

ME561 *aeEx48[Cbr-lin-28::GFP(Y35R F37A C127R C137A) 3’-UTR deletion],*

ME562 *Cbr-lin-41(ae76); Cbr-lin-29(ae75)/+,*

ME563 *Cbr-lin-28(ae39); Cbr-lin-29(ae75)/+,*

ME564 *aeIs15[rps-28pro>atTIR1(F79G)::P2A::GFP::His-11; Cbr-unc-119(+)] Cbr-lin-4(ae79); Cbr-mir-241 Cbr-mir-48(ae73); Cbr-hbl-1(aeIs12[Cbr-hbl-1::AID]) Cbr-mir-84 (ae70),*

ME565 *Cbr-lin-41(ae76); aeIs15[rps-28pro>atTIR1(F79G)::P2A::GFP::His-11; Cbr-unc-119(+)]; Cbr-hbl-1(aeIs12[Cbr-hbl-1::AID]),*

ME566 *Cbr-hbl-1(aeIs12[Cbr-hbl-1::AID]) Cbr-let-7(ae48); aeIs15[rps-28pro>atTIR1(F79G)::P2A::GFP::His-11; Cbr-unc-119(+)]*

ME567 *Cbr-lin-28(ae39); Cbr-mir-48(ae65)*

ME568 *Cbr-dpy-5(v234) +/+ Cbr-lin-28(ae80)*

### Development synchronization

To generate developmentally synchronized populations of worms, adults filled with eggs were washed from crowded plates into 15ml tubes, spinned down, the excess liquid then was removed leaving a worm pellet. 500-1000µl of household bleach solution was added to the tube and vortexed every 2 minutes until cuticles of worms were dissolved enough to release eggs. Tubes then were filled with sterilized distilled water to slow the bleach activity and centrifuged at 400g for 3 minutes, then the supernatant was discarded and eggs were washed twice with sterilized distilled water and then twice with M9. Then eggs were transferred into M9 and left on a shaker for 20-48 hours at room temperature. Then the liquid with larvae was centrifuged to concentrate larvae and they were transferred to plates with food in a small amount of M9.

### Microscopy

Animals were examined using DIC and fluorescence microscopy on a Zeiss Axioplan2 microscope with Zeiss objectives: Plan-NEOFLUAR 5x, Plan-NEOFLUAR 16x, Plan-NEOFLUAR 63x, alpha Plan-FLUAR 100x. Images were acquired using AxioCam with AxioVision software.

### Seam cell and gonad phenotype scoring

Fluorescent seam cell markers to facilitate the counting of seam cells are not yet available for *C. briggsae* as for *C. elegans*. Therefore, there was some variability in seam cell counts for animals of apparently identical age due to occasional difficulty distinguishing seam cells from lateral hypodermal nuclei by DIC microscopy. Seam cells were confidently identified when located along the lateral midline or slightly off and having a visible oval-shaped outline. Errors in counting may occur when seam cell nuclei or hypodermal syncytial nuclei lay away from or close to the apparent midline, respectively.

A disorganized gonad was scored if gonad or oocyte contents leaked into the pseudocoelom or oocytes or spermatozoa were found outside the gonad.

### CRISPR/Cas9

We followed general protocols described in Paix *et al*. 2017. gRNA was synthesized from PCR-amplified templates using Invitrogen™ MEGAshortscript™ T7 Transcription Kit (Catalog# AM1354) and purified with Invitrogen™ MEGAclear™ Transcription Clean-Up Kit (Catalog# AM1908). Purified gRNAs were mixed with Cas9 (EnGen^®^ Spy Cas9 NLS, Catalog# M0646T) and used in microinjections. Typical concentrations of the components in the injection mix: gRNA (up to 200ng/µl), Cas9 (250 ng/µl), co-injection *Cbr-myo::GFP* plasmid (35 ng/µl).

When making insertions, a hybrid dsDNA repair template was used as described in (Dokshin *et al*. 2018). Repair templates were melted and cooled before injections (Ghanta and Mello 2020). The repair template then was added to the CRISPR mix to the final concentration of 100-500 ng/µl of DNA.

### RNAi

RNA interference used to knockdown *Cbr-hbl-1* expression was performed as described (Hammell and Hannon 2016). Part of the *Cbr-hbl-1* ORF flanked by T7 promoters was amplified using these primers: 5’-GCGCGCTAATACGACTCACTATAGGTCCCAGCACCCCTACCACCAC-3’, 5’-GCGCGCTAATACGACTCACTATAGTGGTGACGCCGGCTCTCCTTT-3’

RNA was synthesized and purified with the in-vitro transcription kit mentioned above. Purified RNA was diluted with sterile distilled water to 200 ng/µl concentration and injected into gonads.

### Auxin-inducible degron system

We used a modified auxin-inducible system with *TIR1(F79G)*, using 5-Ph-IAA as the auxin analog (Zhang *et al*. 2015; Hills-Muckey *et al*. 2021). To express *TIR1*, wildtype *C. briggsae* were injected with a modified pCMH2074 plasmid (C. Hammell, pers. comm.) containing *TIR1(F79G)* mutation and GFP in place of mCherry. Bright fluorescing animals carried a stable array with a high inheritance rate (50-75%) were selected and a strain with stable extrachromosomal expression was established (*aeEx44*).

To integrate *TIR1(F79G)* into the genome, fluorescent L4 and young adult animals were transferred to a plate without bacteria and irradiated with UV in an UV-crosslinker set to an energy level of 13 kJ/cm^2 to generate chromosomal breaks and attach the array to a chromosome. Fluorescent F1 animals were isolated and then plates were screened for 100 or 75% transmission rates or for animals displaying uniform (vs. mosaic) fluorescence. Stable integrants were identified as animals with uniform fluorescence with 100% penetrance. This strain then was outcrossed at least 3 times.

To assess phenotypes using AID system, animals carrying alleles with fused degron tag and expressing *TIR1(F79G)* were grown on plates containing 50µmol of 5-Ph-IAA that was spread on standard NGM plates to approximately 0.005 µM concentration in the agar.

### Cbr-lin-28::GFP plasmid

The *Cbr-lin-28::GFP* expression plasmid was produced by PCR and restriction digestion and ligation techniques. Q5^®^ High-Fidelity DNA Polymerase (NEB #M0491S) and Phusion^®^ High-Fidelity DNA Polymerase (NEB #M0530S) were used for amplification. *Cbr-lin-28* with promoter region and 3’UTR were amplified from the *C. briggsae* (AF16) genome, and the *C. elegans*-optimized GFP sequence with introns was amplified from pVT221 (Moss *et al*., 1997). To mutate the plasmid, a Q5^®^ Site-Directed Mutagenesis Kit (NEB #E0552S) was used. The plasmid scheme and mutation sequences are shown in Fig. S10.

### Plots and statistics

Data was analyzed using Prism software. *P*-values were calculated using unpaired Welch’s t-tests for absolute values (seam cell and intestinal nuclei count averages) and Fisher’s exact tests for fractions (percents). Error bars in plots indicate 95% CI, asterisks indicate the following: * *p* ≤ 0.05, ** *p* ≤ 0.01, *** *p* ≤ 0.001, **** *p* ≤ 0.0001, ns - not significant (*p* > 0.05).

## Acknowledgements

We would like to thank members of Ellis lab and Kevin Kemper for valuable comments on the manuscript, Ronald Ellis for comments, advice and strains, Chris Hammell for TIR1 strains and plasmid.

## Conflict of Interest Statement

Authors have no conflicts of interest to declare.

## Data availability Statement

Strains are available upon request.

## Supplementary materials

### Supplementary tables

**Table S1:**
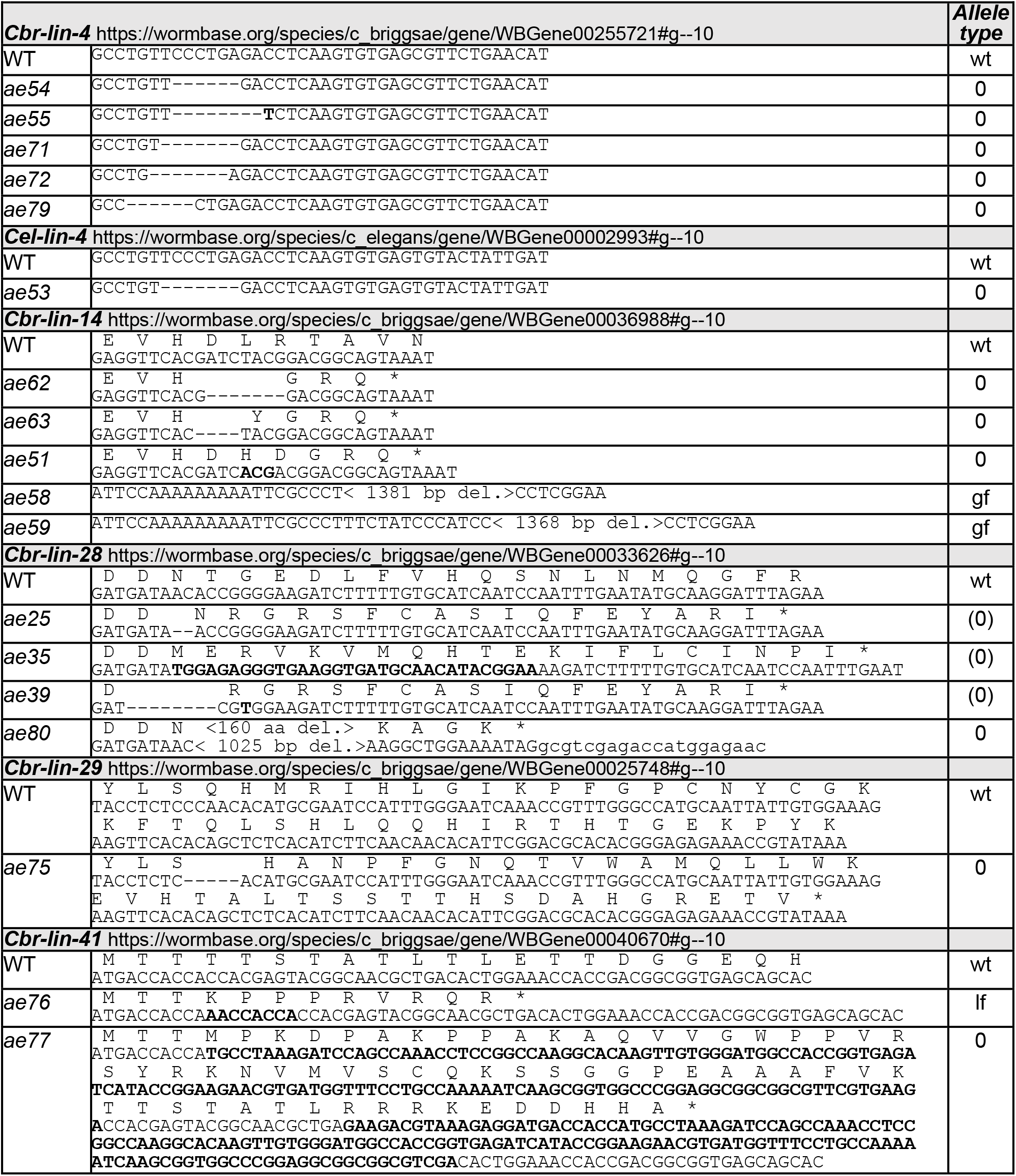
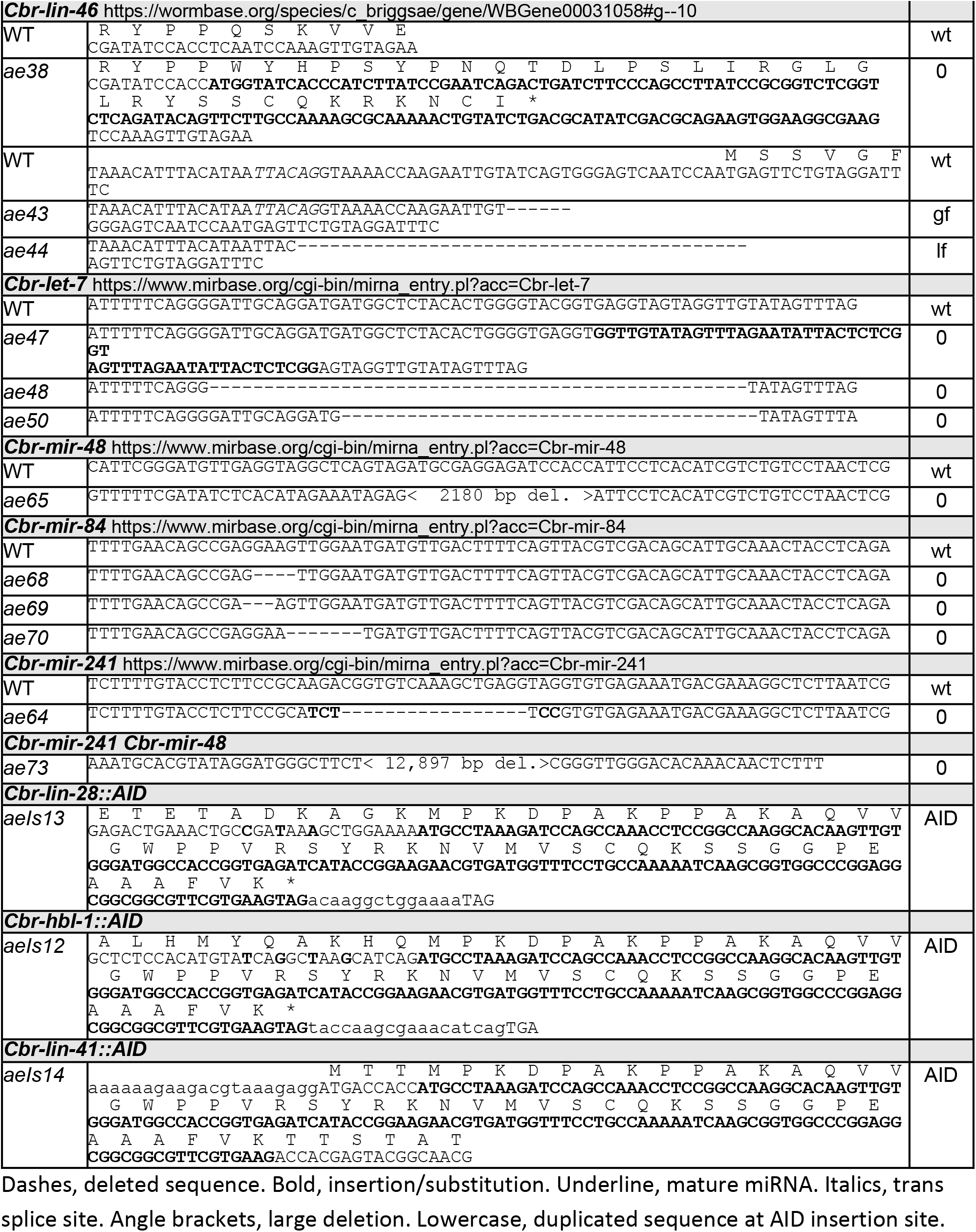
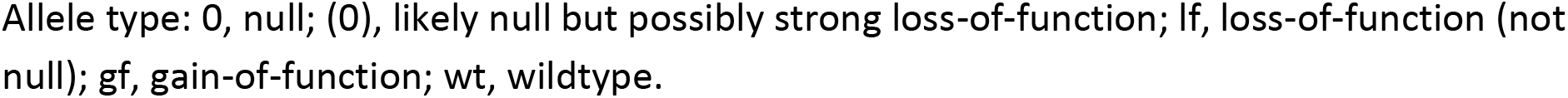
Genetic lesions.

**Table S2:** adult alae in *Cbr-lin-4(0)* and *Cbr-lin-14(gf)* strains.

| Alae completeness | *Cbr-lin-4(ae55)*<br>young adults<br>n = 12 | *Cbr-lin-4(ae55)*<br>egg-producing adults<br>n = 18 | *Cbr-lin-14(ae58) 3'UTR Δ*<br>egg-producing adults<br>n = 21 |
| --- | --- | --- | --- |
| Full | 8% | 17% |  |
| Gaps | 8% | 78% |  |
| Half | 17% | 6% |  |
| Patches | 50% |  | 48% |
| No alae | 17% |  | 52% |

Alae gaps mean that more than 50%, but less than 100% of seam cells generated adult alae, alae patches - more than 0% but less than 50%, half means that approximately 50% of seam cells generated adult alae.

**Table S3:** Intestinal nuclei number in strains expressing *TIR1(F79G)*.

| Genotype | Average number of intestinal nuclei at L3 and older |
| --- | --- |
| Wild type | 33.1+/-1.8 (n = 60) |
| *TIR1(F79G) II* without auxin | 25.7+/-3.3 (n = 17) |
| *TIR1(F79G) II*; *Cbr-hbl-1::AID* on 5-Ph-IAA | 22.6+/-2.5 (n = 10) |
| *TIR1(F79G) II*; 3let-7s, *Cbr-hbl-1::AID* on 5-Ph-IAA | 25.7+/-2.7 (n = 15) |

### Supplementary figures and legends

**Fig. S1.**
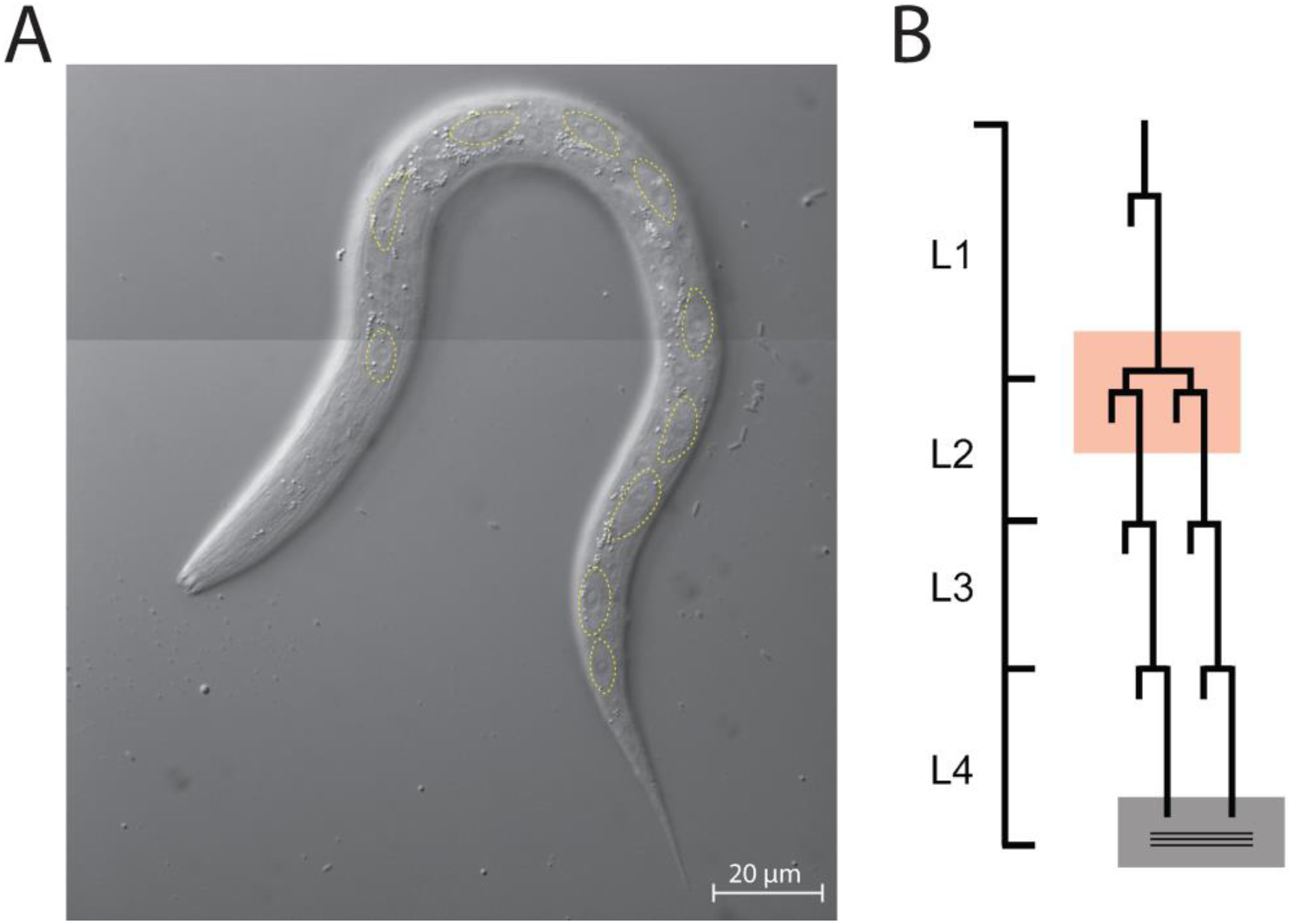
Seam cell lineages of *C. briggsae* are identical to those of *C. elegans*. (A) DIC microphotograph of an L1 worm less than 3 hours after hatching. Ten seam cells are outlined with the dotted line. (B) Cell lineage diagram summarizing the divisions of V1-V4 and V6 seam cells during the four larval stages. Vertical lines indicate the passage of time, and horizontal lines indicate a cell division, with those that stop indicating terminal differentiation, either by joining the hypodermal syncytium, or by differentiation into adult seam cells that produce alae (gray square with horizontal lines). The red square highlights symmetric cell divisions that each produce two daughters that remain at the seam and continue dividing. This cell lineage is identical for *C. elegans* and *C. briggsae*.

**Fig. S2.**
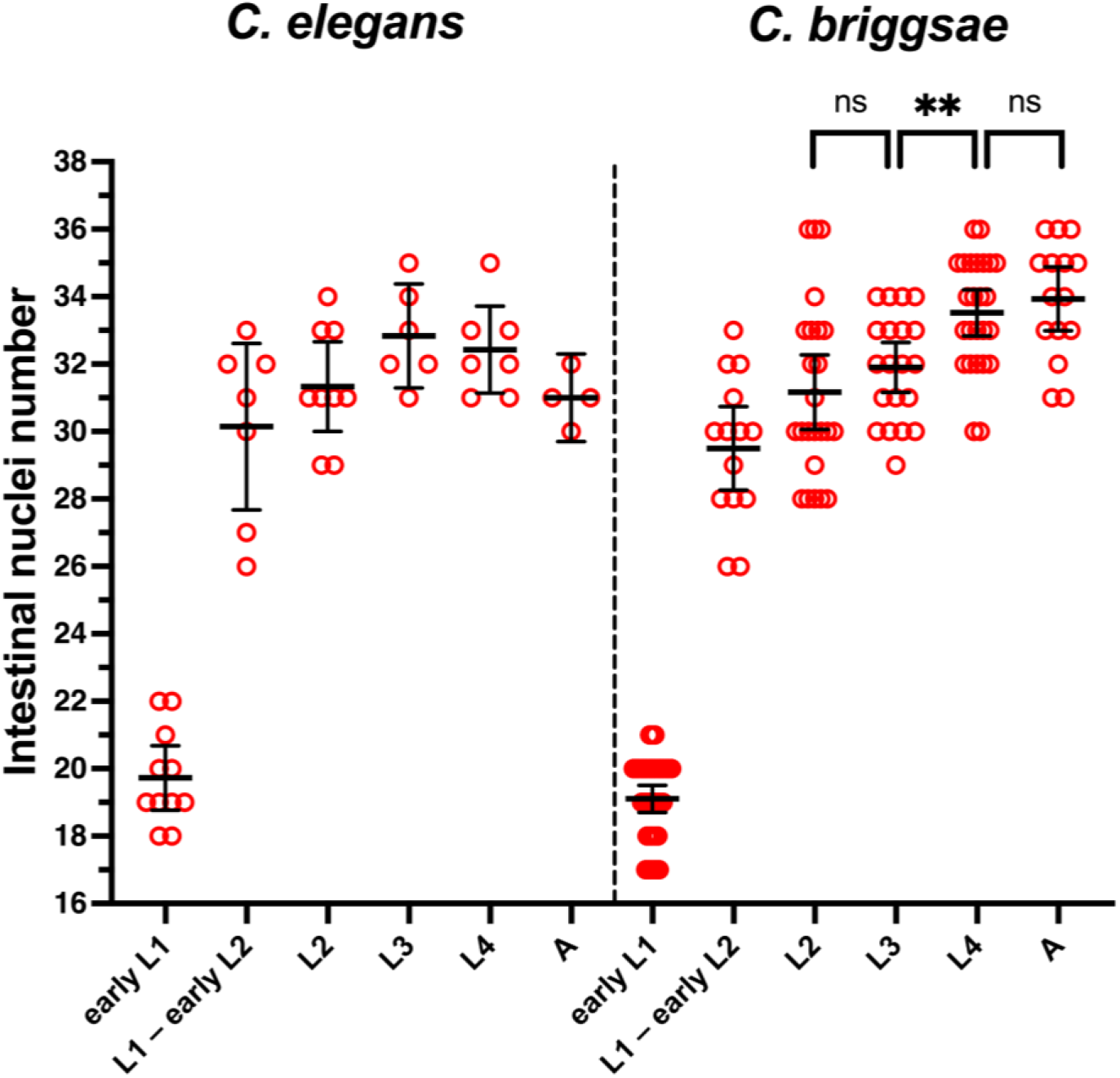
Intestinal nuclei divide after the L1 stage in *C. elegans* and *C. briggsae*. Division in *C. briggsae* occurs after the L2 stage, but not in *C. elegans*. “Early L1” refers to animals before the L1 seam cell divisions. “L1 – early L2” refers to animals during and after the L1 seam cells divisions but before the L2 (symmetric) seam cell divisions. “L2” refers to animals both during and after the L2 seam cell divisions. Statistical analysis is described in Materials and Methods.

**Fig. S3.**
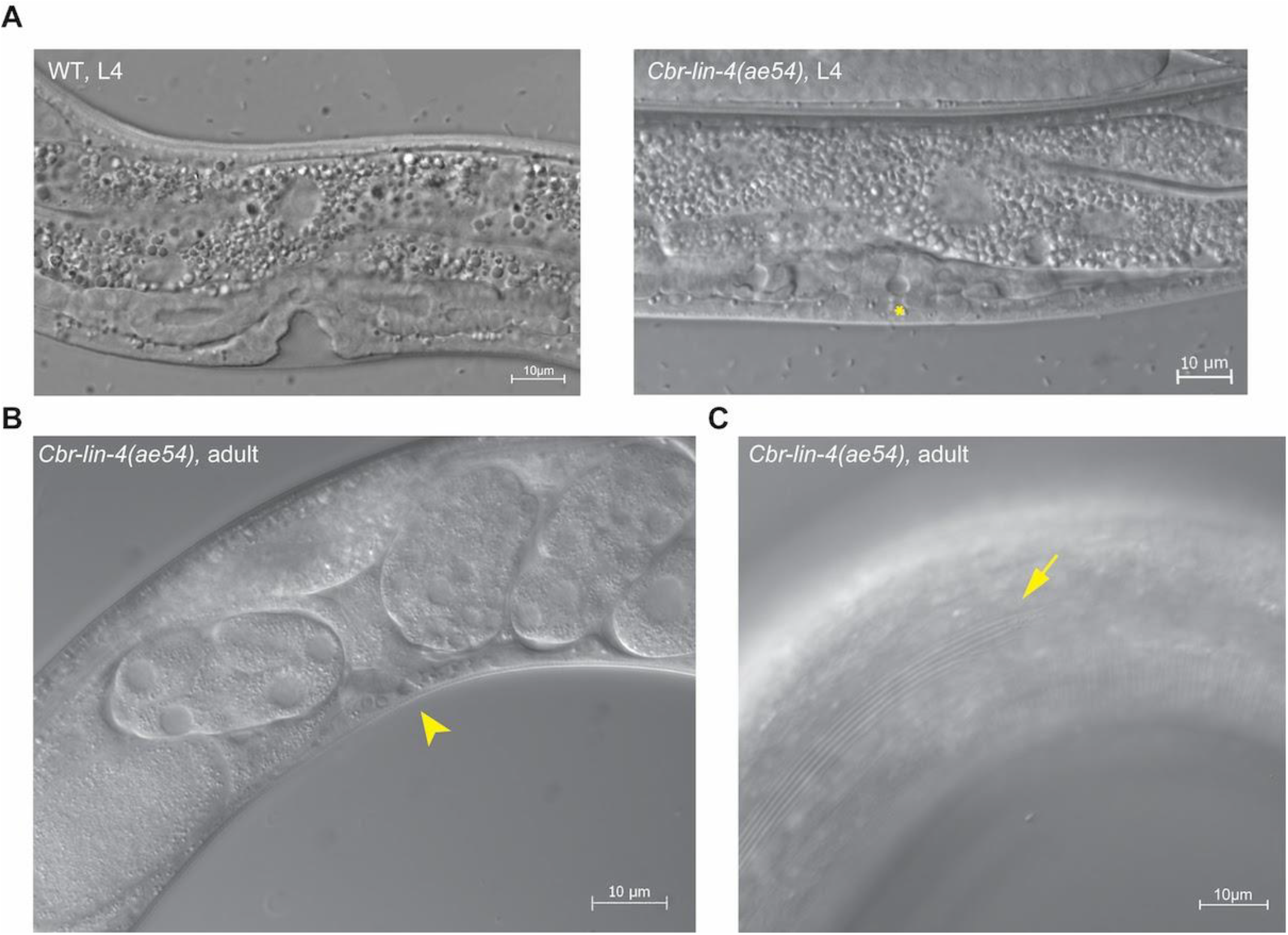
*Cbr-lin-4(0)* mutants have a reiterative phenotype. DIC micrographs showing (A) the vulval region in a *Cbr-lin-4(ae54)* L4 animal compared to a wild-type L4 animal. The vulval precursor cells failed to divide. Asterisk indicates the vulva area, and above the asterisk is the lysed anchor cell. (B) *Cbr-lin-4(ae54)* adult where the vulva failed to form resulting in the accumulation of embryos in the uterus. Arrowhead points at the vulva area. (C) Gapped alae in an adult *Cbr-lin-4(ae54)* animal; arrow points at the alae gap.

**Fig. S4.**
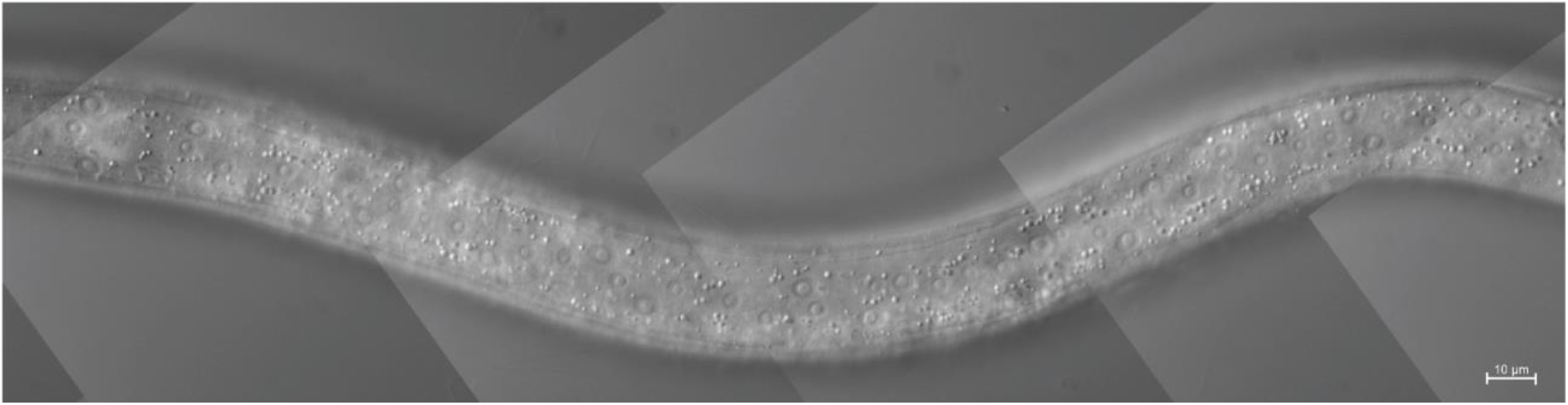
*Cbr-lin-4(0)* mutants reiterate L2 seam cell fates. DIC microphotograph of *Cbr-lin-4(ae54)* L3 larva with an increased number of seam cells (elongated cells located at the midline).

**Fig. S5.**
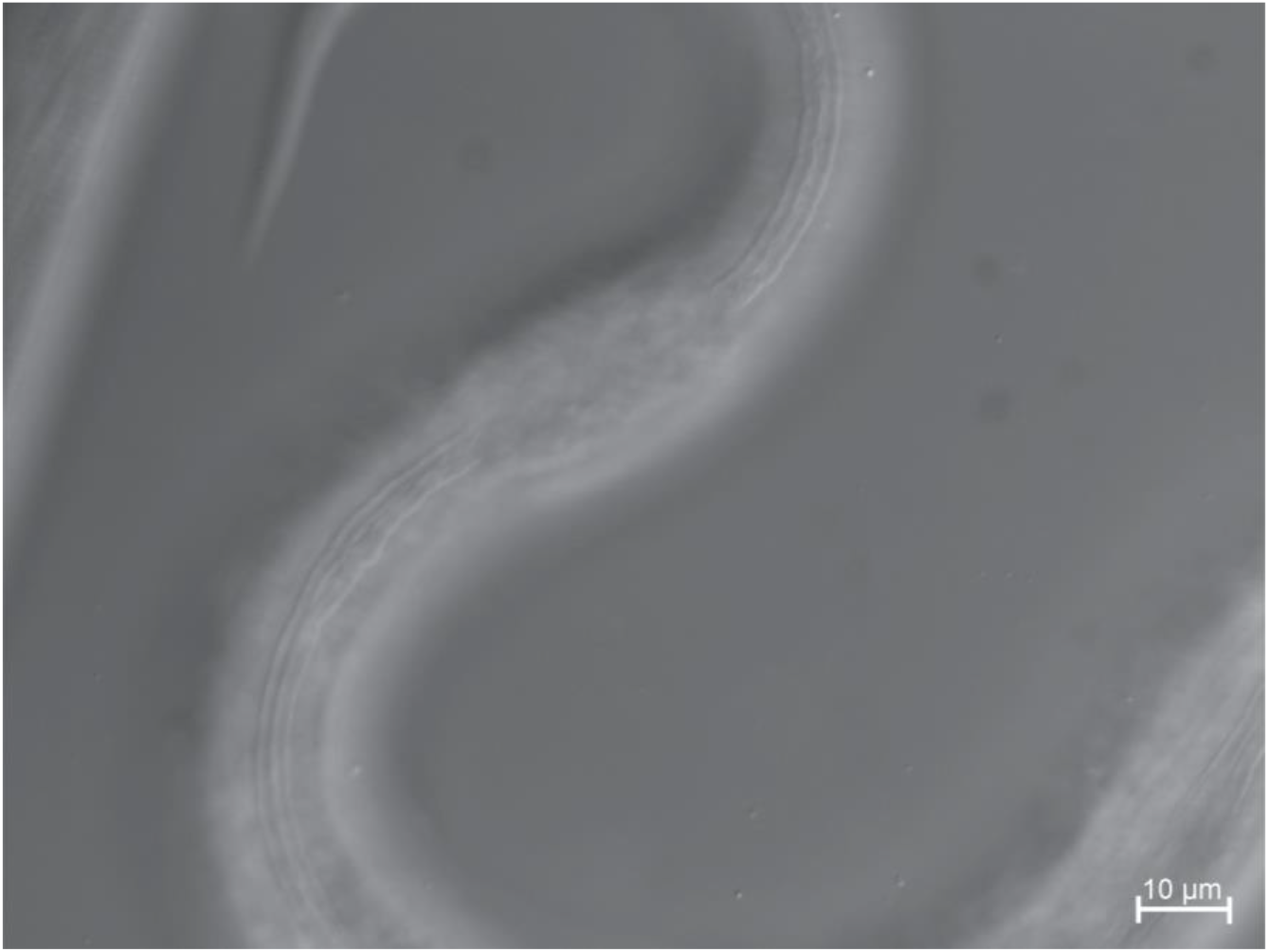
*Cbr-lin-4(0)* mutants can become dauers but incompletely. DIC microphotograph of the central body of a *Cbr-lin-4(ae54)* dauer larva. Part of its body is swollen and lacks the dauer cuticle.

**Fig. S6.**
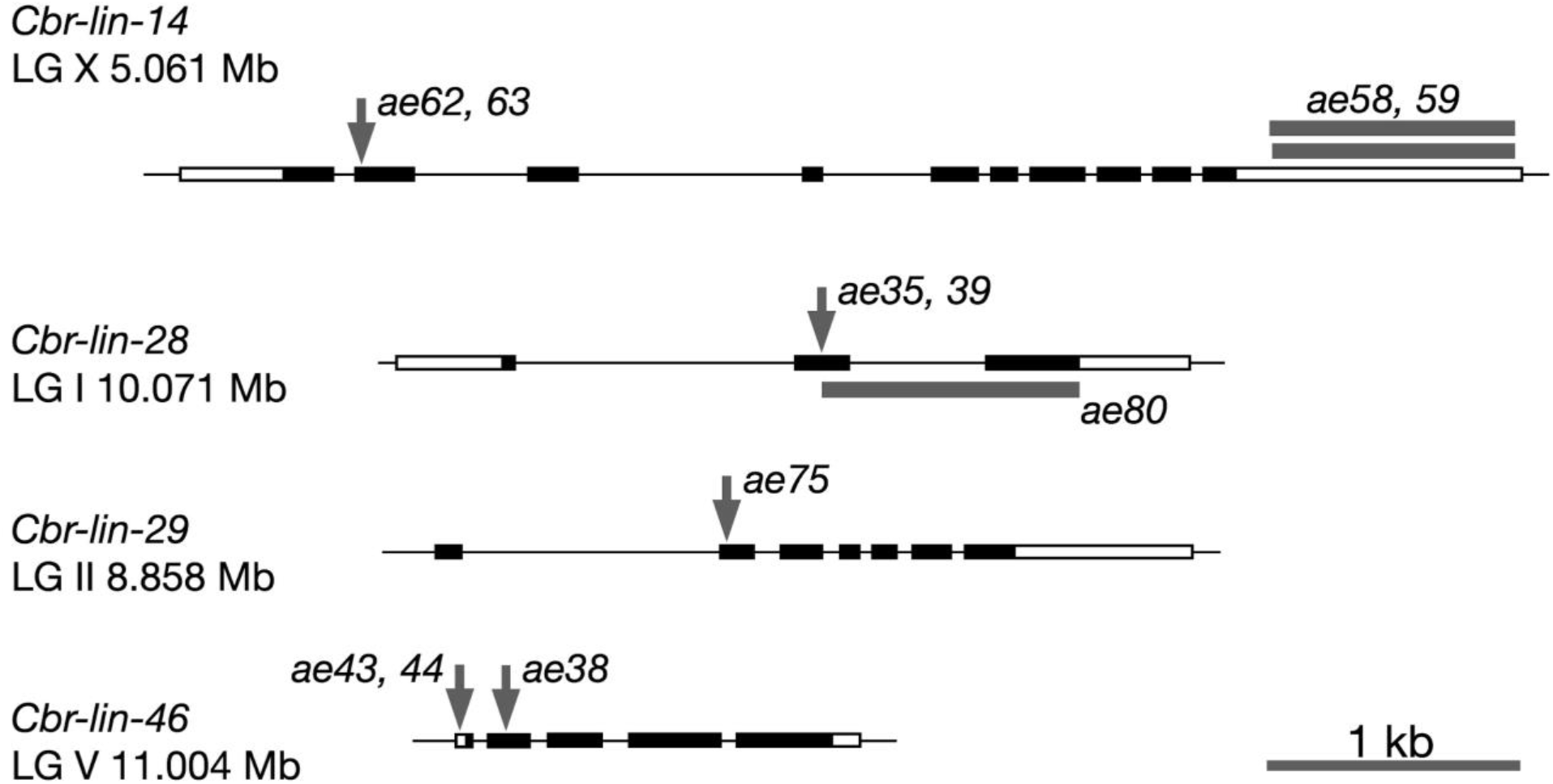
Genomic locations of genetic lesions listed in Table S1. Black rectangles, coding exons. White rectangles, non-coding regions. Arrows indicate lesions listed in Table S1. The bars for *Cbr-lin-14* alleles *ae58* and *ae59* show the extent of the deletions. Lesions and modifications not shown were at either the 5’ or 3’ end of the coding regions, as indicated in Table S1 and described in the text.

**Fig. S7.**
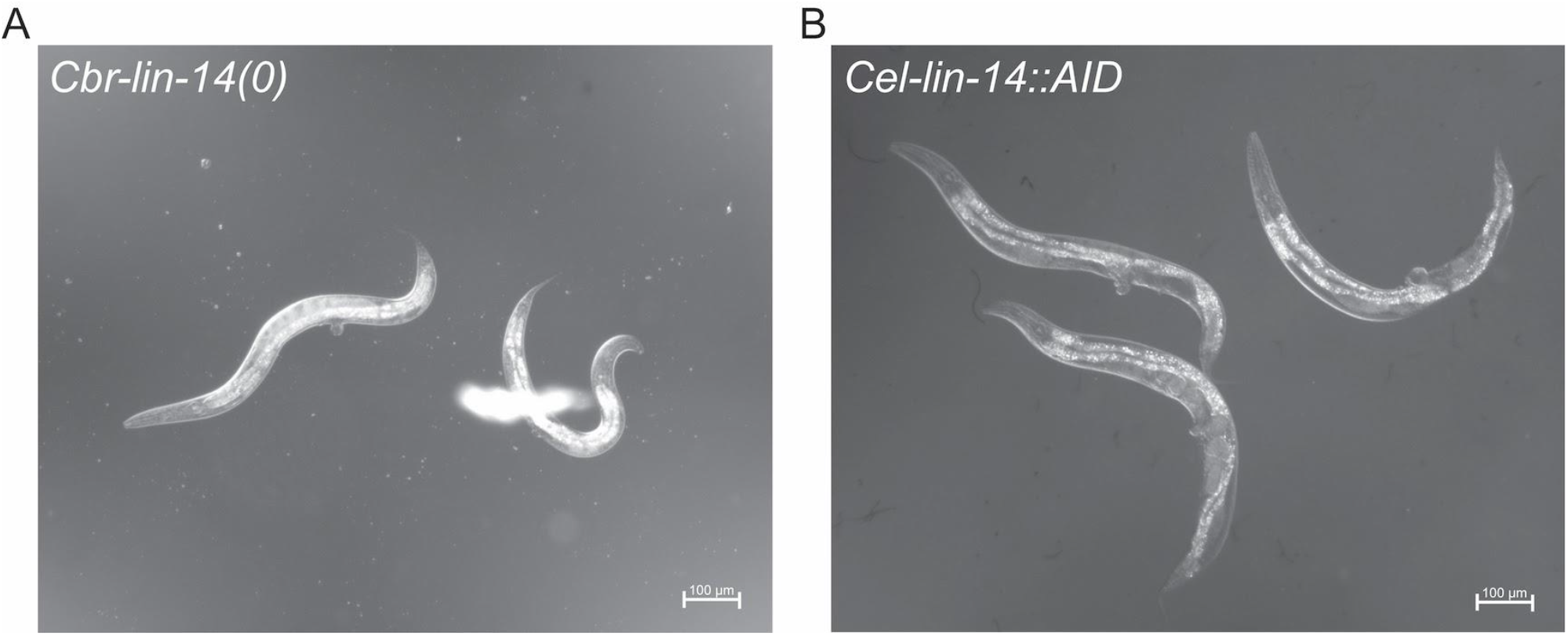
The overall morphology of *Cbr-lin-14(0)* mutants is almost identical to that of *Cel-lin-14(lf)* mutants. DIC microphotographs of *Cbr-lin-14(ae62)* mutant and *Cel-lin-14::AID* mutants on 5-Ph-IAA (20°C).

**Fig. S8.**
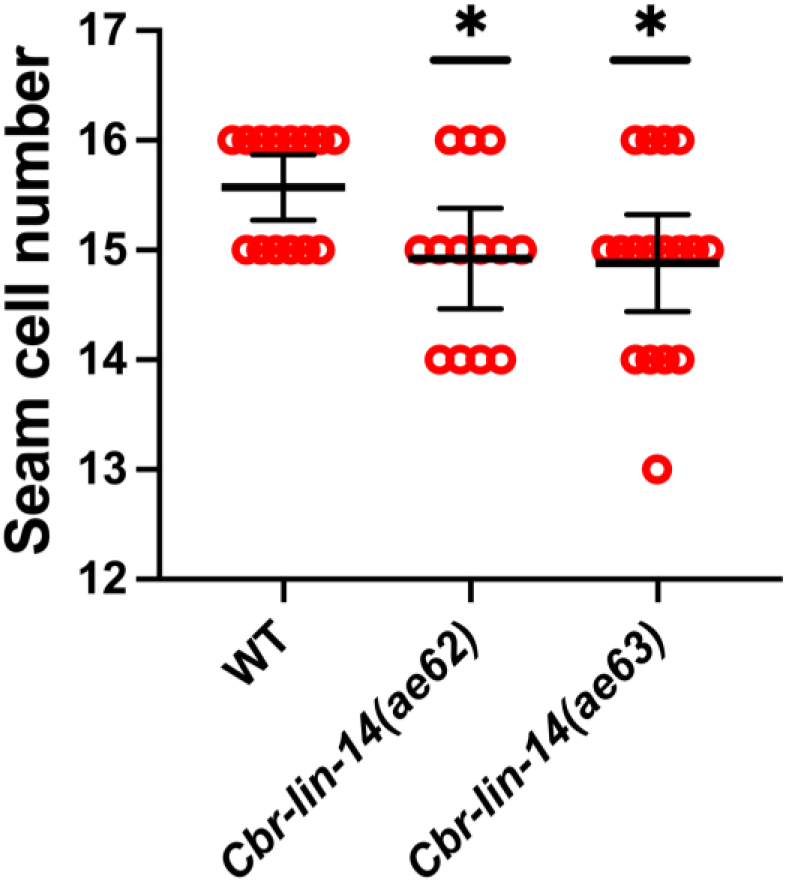
The number of seam cells in *Cbr-lin-14(0)* mutants is slightly reduced compared to the wild type. Seam cells were counted in L4 animals for two *Cbr-lin-14(0)* alleles, *ae62* and *ae63*. A small reduction in the seam cell number might be a result of skipping of the L2 divisions by some seam cells. Statistical analysis is described in Materials and Methods.

**Fig. S9.**
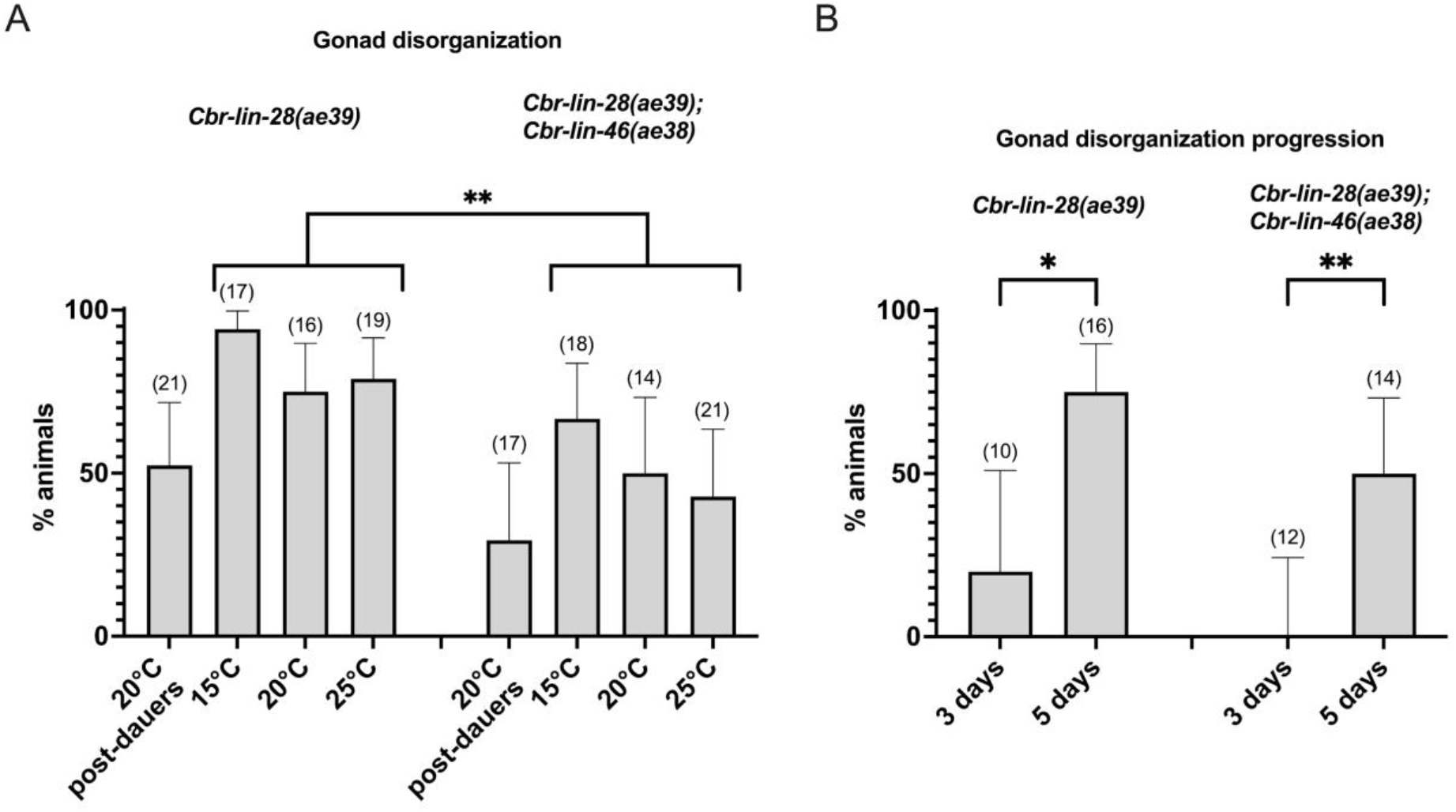
Gonad disorganization manifests with age and is slightly suppressed by *Cbr-lin-46(0)* in *Cbr-lin-28(0)* mutants. (A) Gonad disorganization in *Cbr-lin-28(0)* mutants is partially suppressed by the dauer pathway, although *p-value* was > 0.05 between post-dauers and normal animals at 20°C. *Cbr-lin-46(0)* mutation partly suppressed *Cbr-lin-28(0)* gonad disorganization, *p* < 0.005 when the data from all temperatures was combined in one data set in each strain. (B) Gonad disorganization manifests with age. In both *Cbr-lin-28(0)* and *Cbr-lin-28(0)*; *Cbr-lin-46(0)* double mutants, gonad disorganization was observed in few animals within 24 hours after they reached the adult or arrested L4 stage, 2 days later a higher fraction of animals had disorganized gonads (20°C). Animals’ age after hatching is indicated below the chart. Sample sizes are specified in parenthesis above the bars. Statistical analysis is described in Materials and Methods.

**Fig. S10.**
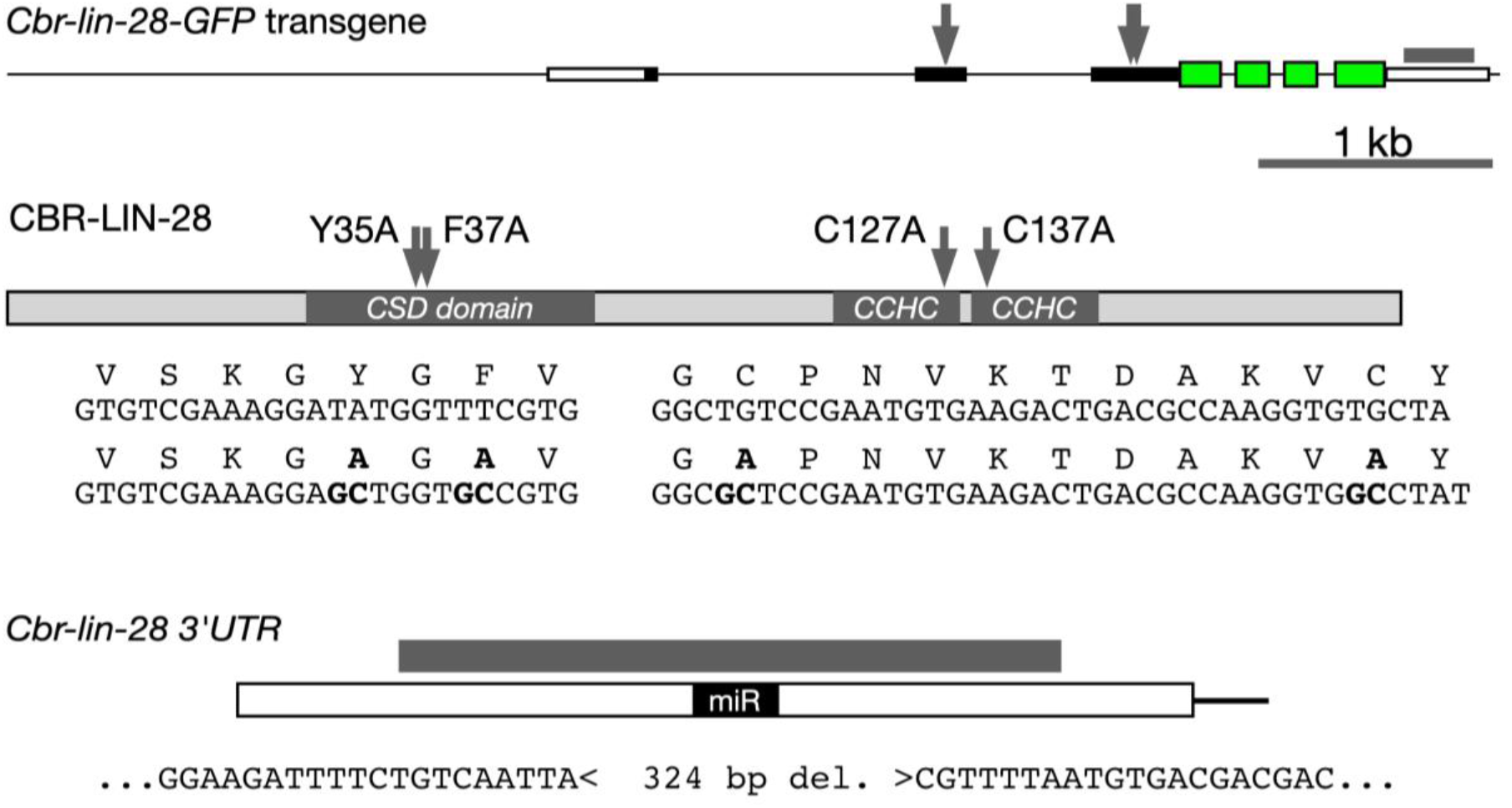
*Cbr-lin-28::GFP* transgenes. Top: An approximately 5.5 kb genomic sequence containing the *Cbr-lin-28* coding region and 5’ and 3’ regulatory regions was cloned into a plasmid. A nematode-optimized GFP was fused in-frame upstream of the stop codon (green). Arrows show locations of mutations in CSD and CCHC domains that inactivate the protein. Gray box above the 3’UTR shows the extent of the deletion that removes the miRNA sites. Middle: A schematic of the CBR-LIN-28 protein showing the locations of inactivating mutations in the CSD domain and CCHC motifs. Sequences of wild type and mutant transgenes are shown. Bottom: A schematic of the *Cbr-lin-28* 3’UTR showing the location of the recognition sites for let-7 and lin-4 miRNAs (miR). Grey bar shows the extent of the deletion removing the miRNA sites. The sequences flanking the deletion are shown.

**Fig. S11.**
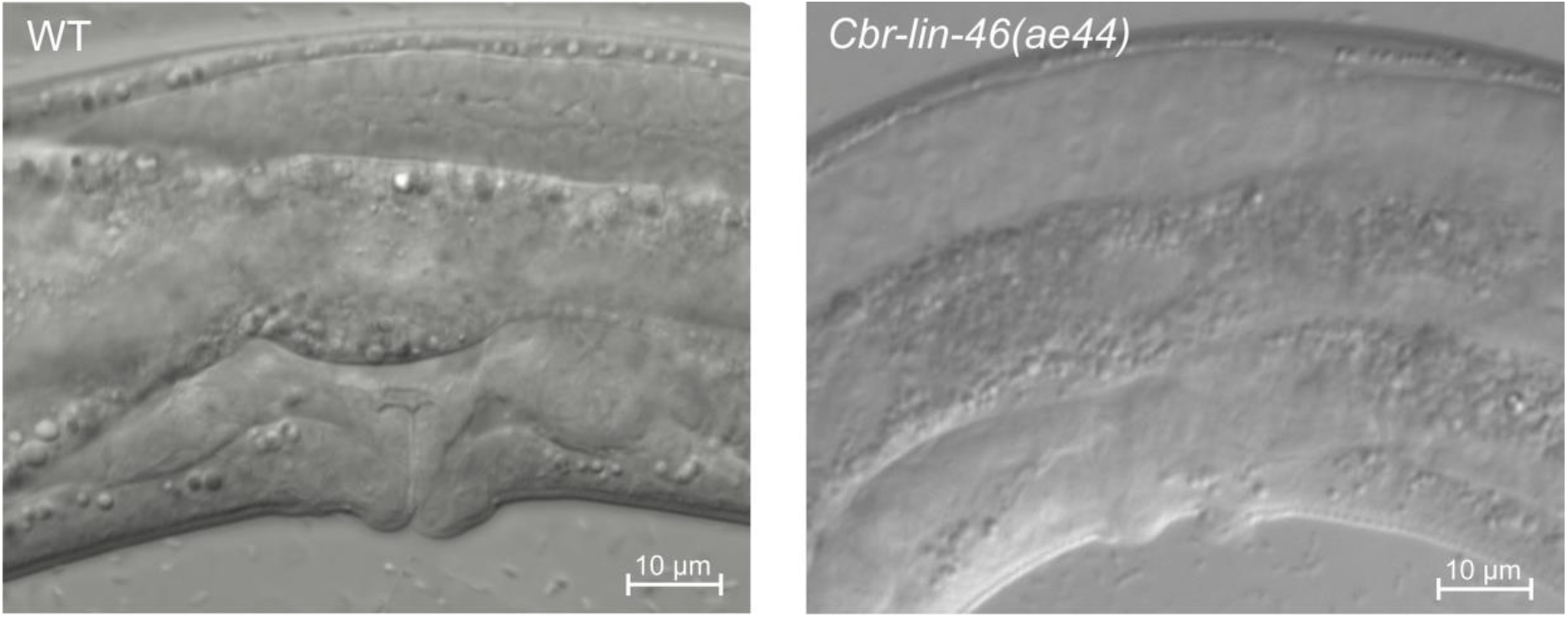
*Cbr-lin-46(0)* mutants have slight vulval developmental defects. A DIC microphotograph of a *Cbr-lin-46(ae44)* animal with vulval protrusions (20°C) compared to vulva of a wild-type animal.

**Fig. S12.**
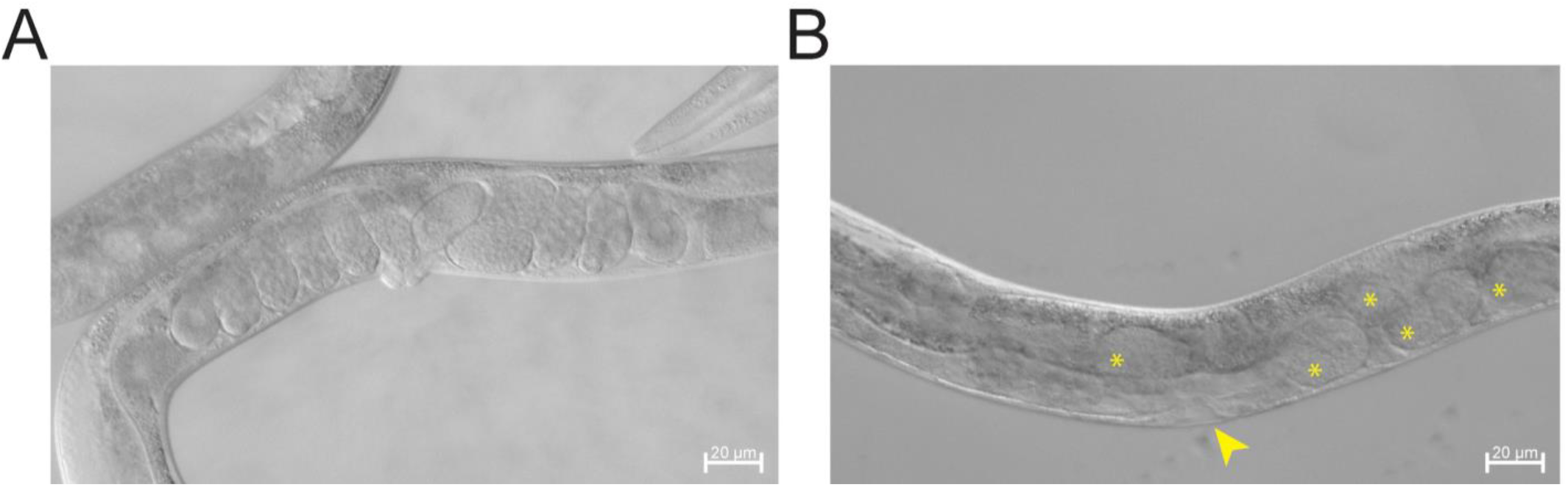
*Cbr-lin-28(0); Cbr-lin-46(0)* sometimes arrest at L4. DIC microphotographs showing (A) a successfully molted *Cbr-lin-28(ae39); Cbr-lin-46(ae38)* animal with adult vulva and eggs; (B) a *Cbr-lin-28(ae39); Cbr-lin-46(ae38)* animal with an arrested L4 phenotype. Arrowhead points at the L4-shaped vulva and asterisks indicate embryos.

**Fig. S13.**
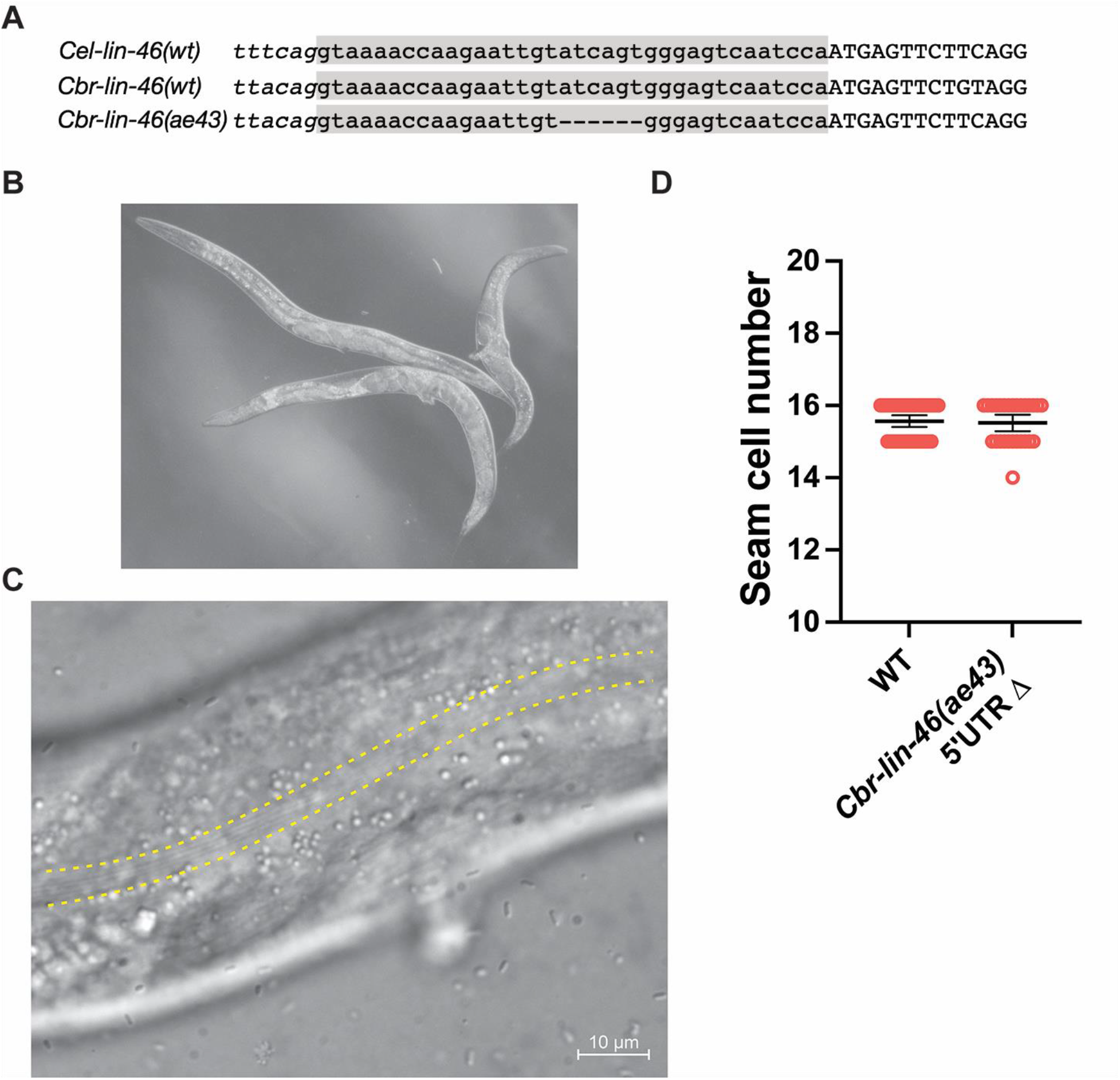
A deletion in *Cbr-lin-46*’s 5’UTR causes a precocious phenotype. (A) The 5’UTR of *C. elegans* and *C. briggsae lin-46* and the *Cbr-lin-46(ae43)* allele. The gray box highlights the conserved 36-nt 5’UTR, with a splice acceptor for a trans-spliced leader to the left and the start of the ORF to the right. (B) DIC microphotograph of *Cbr-lin-46* 5’UTR deletion mutants, showing protruding vulvae and accumulated eggs. (C) DIC microphotograph of a cuticle of *Cbr-lin-46(ae43)* L4 larva with precocious alae (outlined by a dotted line). (D) *lin-46* 5’UTR deletion mutants do not skip L2 stages (20°C).

**Fig. S14.**
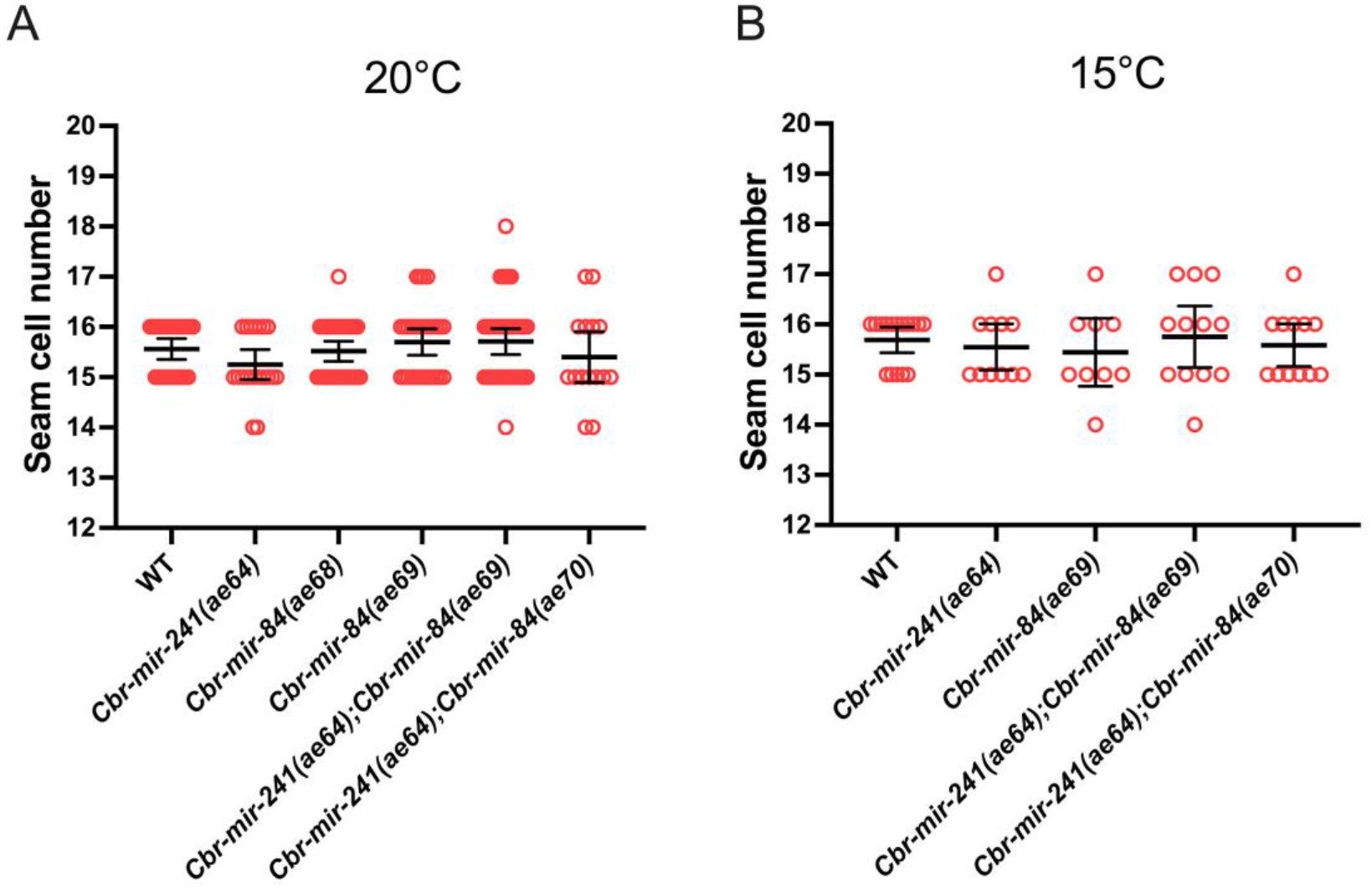
*Cbr-mir-241(0)*, *Cbr-mir-84(0)*, single and double mutants do not have reiterations of L2 stages at (A) 20°C or (B) 15°C. There was no statistically significant difference between WT and any other group at both temperatures (*p-value* > 0.05). Statistical analysis is described in Materials and Methods.

**Fig. S15.**
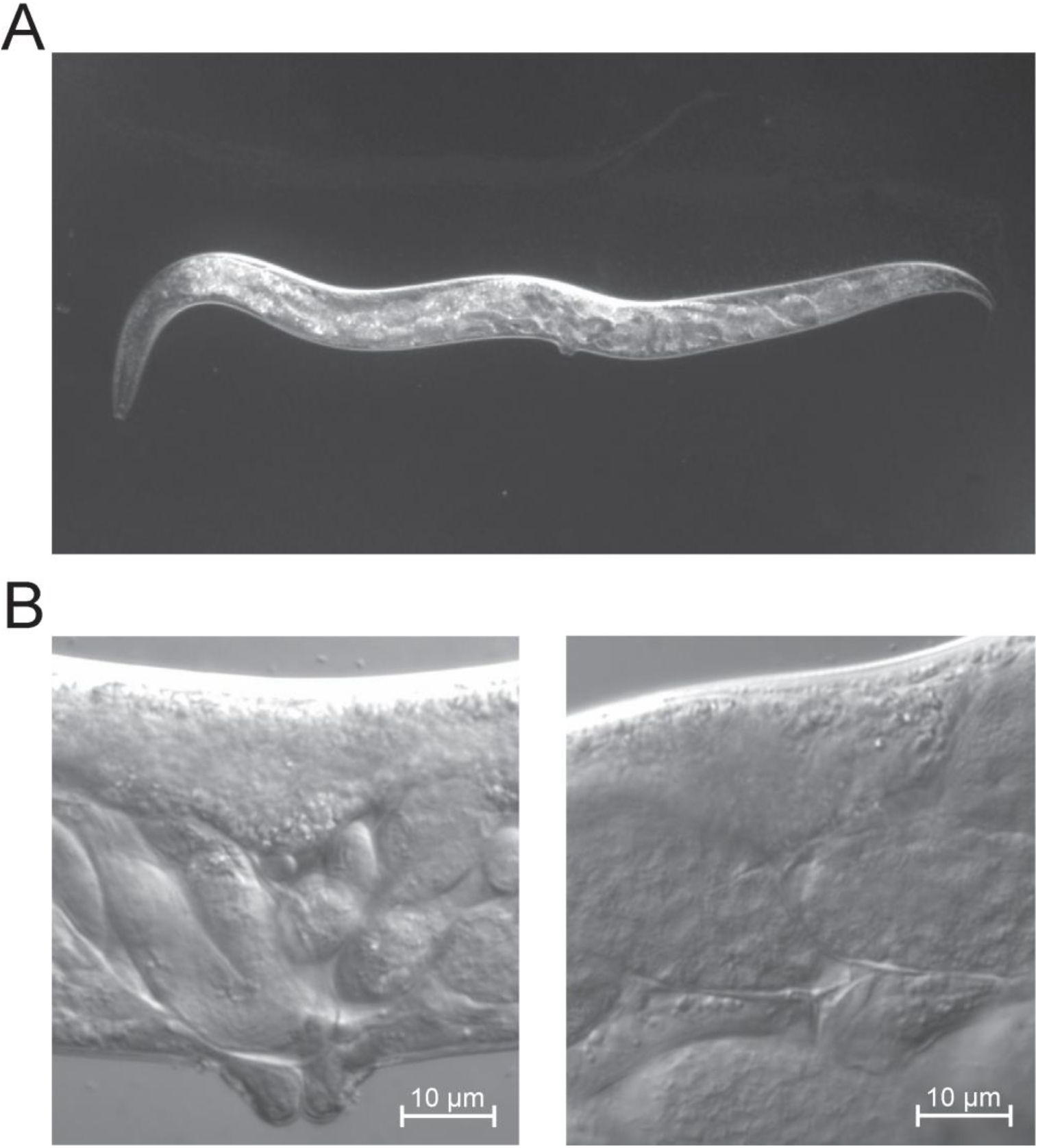
*Cbr-mir-241(0)*; *Cbr-mir-84(0)* double mutants occasionally develop egg-laying defects. DIC microphotographs showing (A) a *Cbr-mir-241(ae64); Cbr-mir-84(ae70)* double mutant with accumulated eggs; other *Cbr-mir-241(0), Cbr-mir-84(0)* single and double mutants had similar occasional phenotypes; (B) slightly abnormal vulvae shapes of the *Cbr-mir-241(ae64);Cbr-mir-84(ae70)* double mutants; other *Cbr-mir-241(0), Cbr-mir-84(0)* single and double mutants occasionally had similar abnormal vulvae. Compare to Fig. S11.

**Fig. S16.**
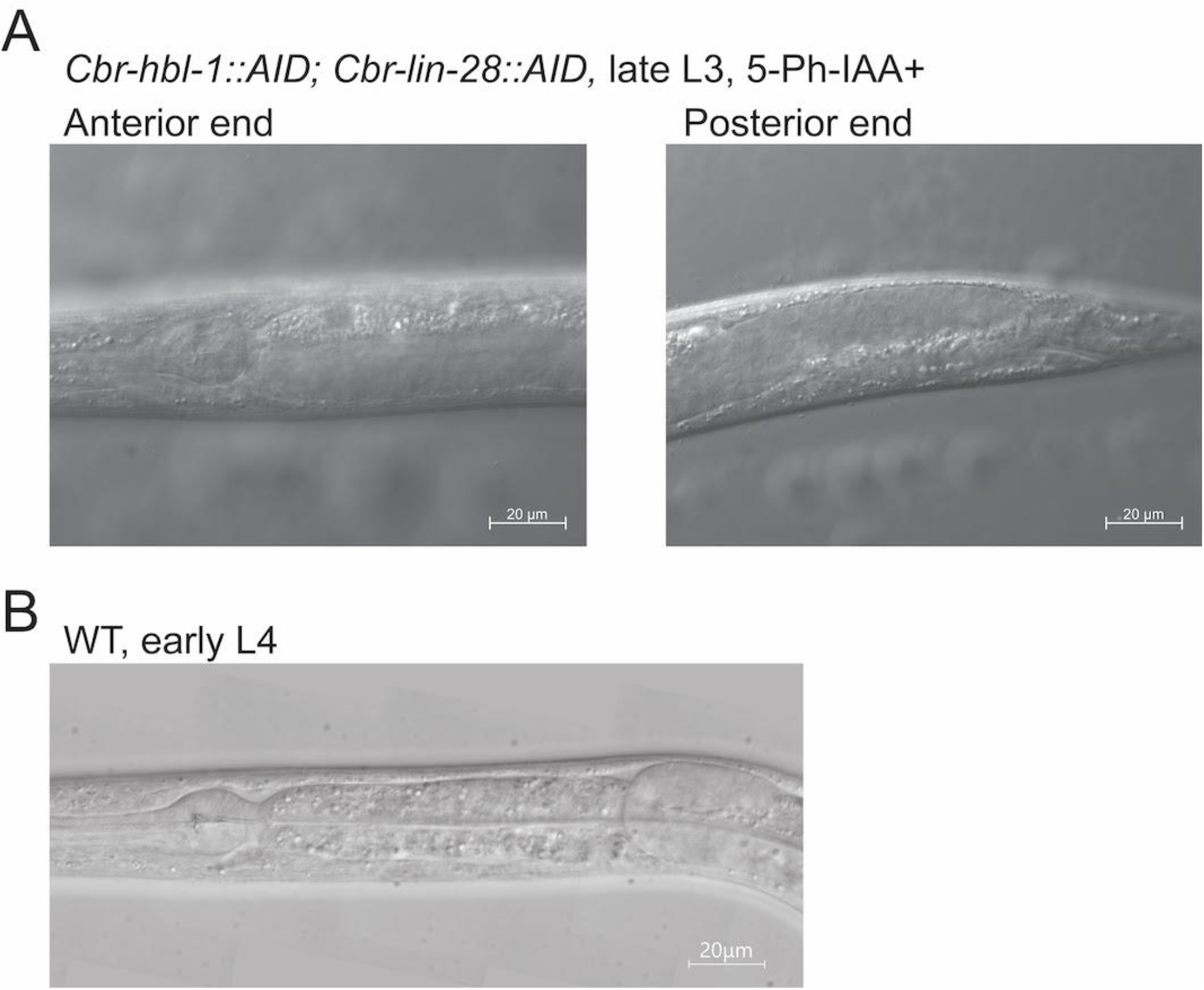
Simultaneous depletion of *Cbr-lin-28* and *Cbr-hbl-1* leads to gonad migration defects. DIC microphotographs showing (A) gonad arms in a *Cbr-hbl-1::AID; Cbr-lin-28::AID* late L3 worm on 5-Ph-IAA; gonad arms are extended towards the ends instead of reflexing compared to (B) wild-type early L4 animal with normal gonad reflexion at anterior end. Also, compare to Fig. 2A. Animals are oriented anterior end left, dorsal side up.

**Fig. S17.**
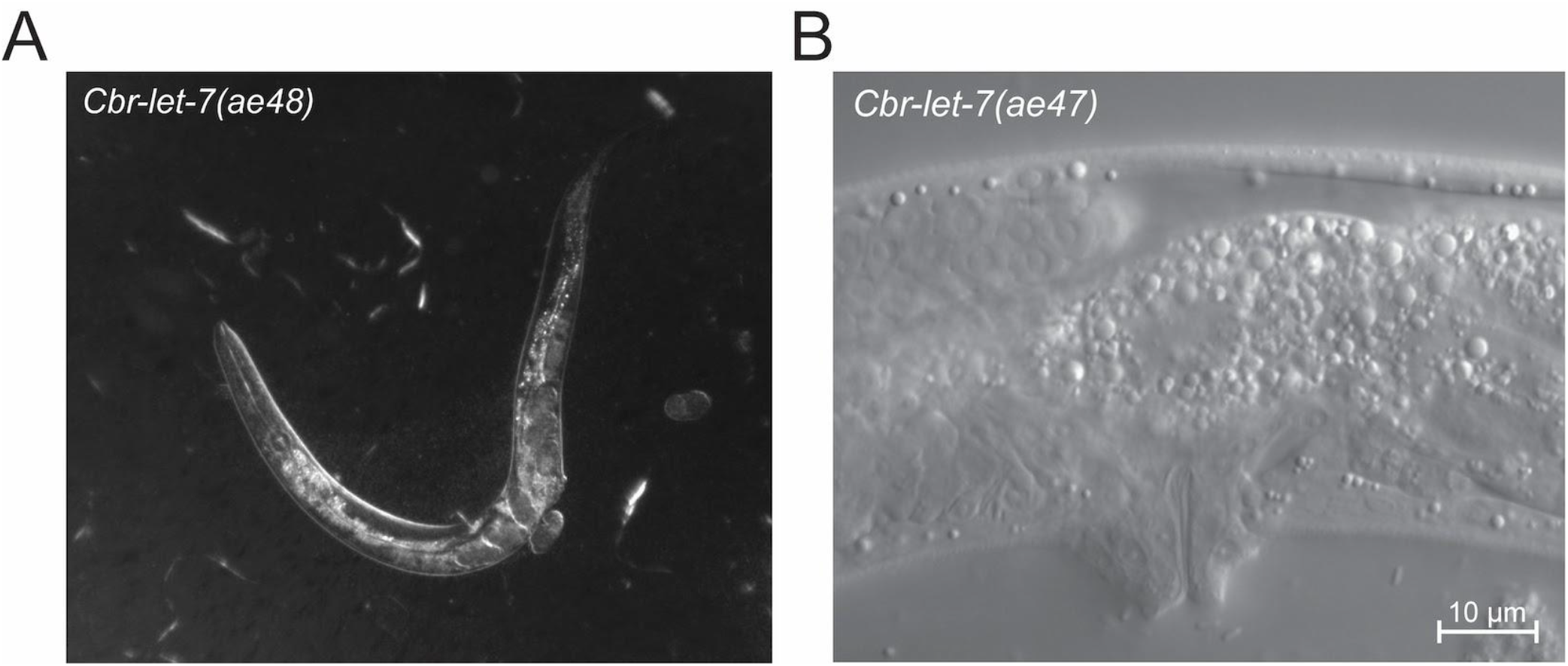
*Cbr-let-7(0)* mutants have egg-laying defects. DIC micrographs of (A) a *Cbr-let-7(ae48)* mutant with accumulated eggs. (B) Vulva of *Cbr-let-7(ae47)*. Compare to Fig. S11. Mutant vulvae protrude more. Animals are oriented anterior end left, dorsal side up.

**Fig. S18.**
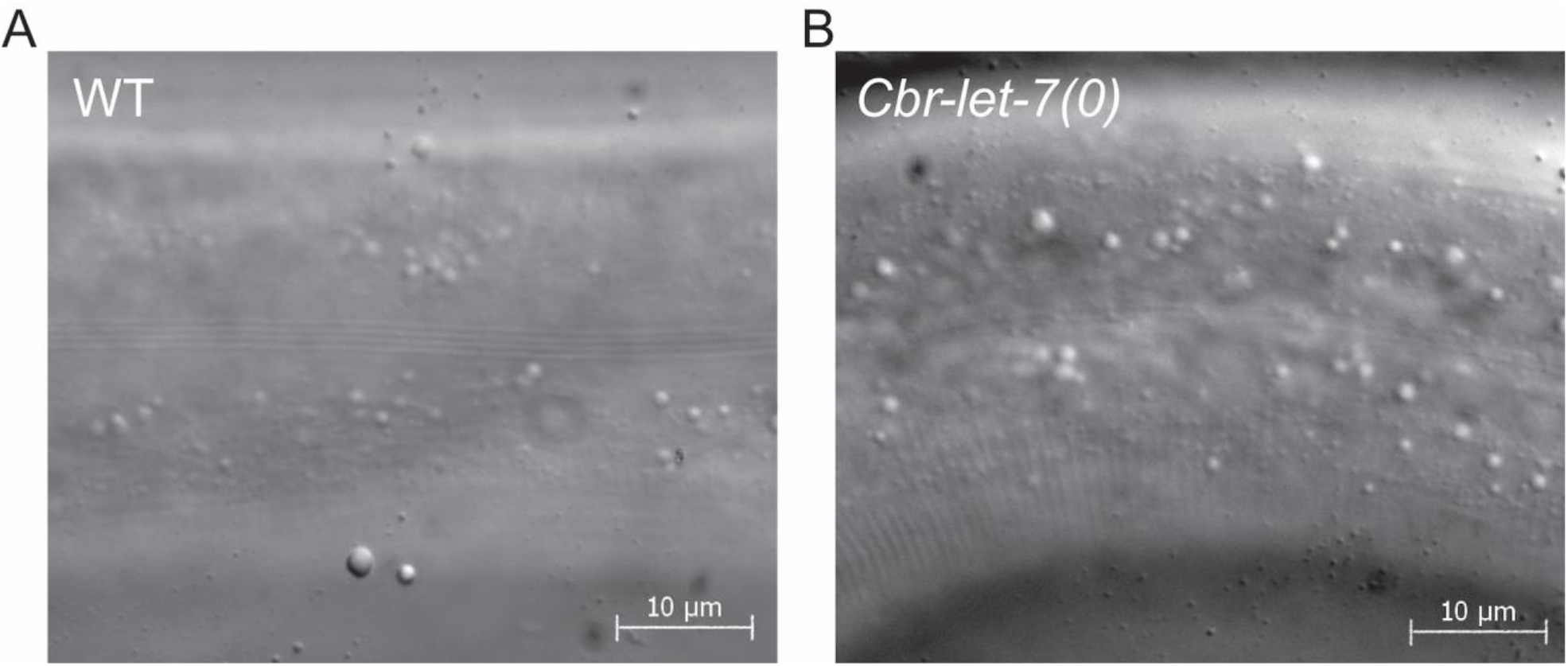
*Cbr-let-7(0)* mutants have thinner alae than the wild type. DIC micrographs of (left) Wild type *C. briggsae* adult alae, and (right) *Cbr-let-7(ae47)* adult alae in animals of approximately the same age. In *Cbr-let-7(0)* mutants adult alae appear to be thinner. Animals are oriented anterior end left, dorsal side up.

**Fig. S19.**
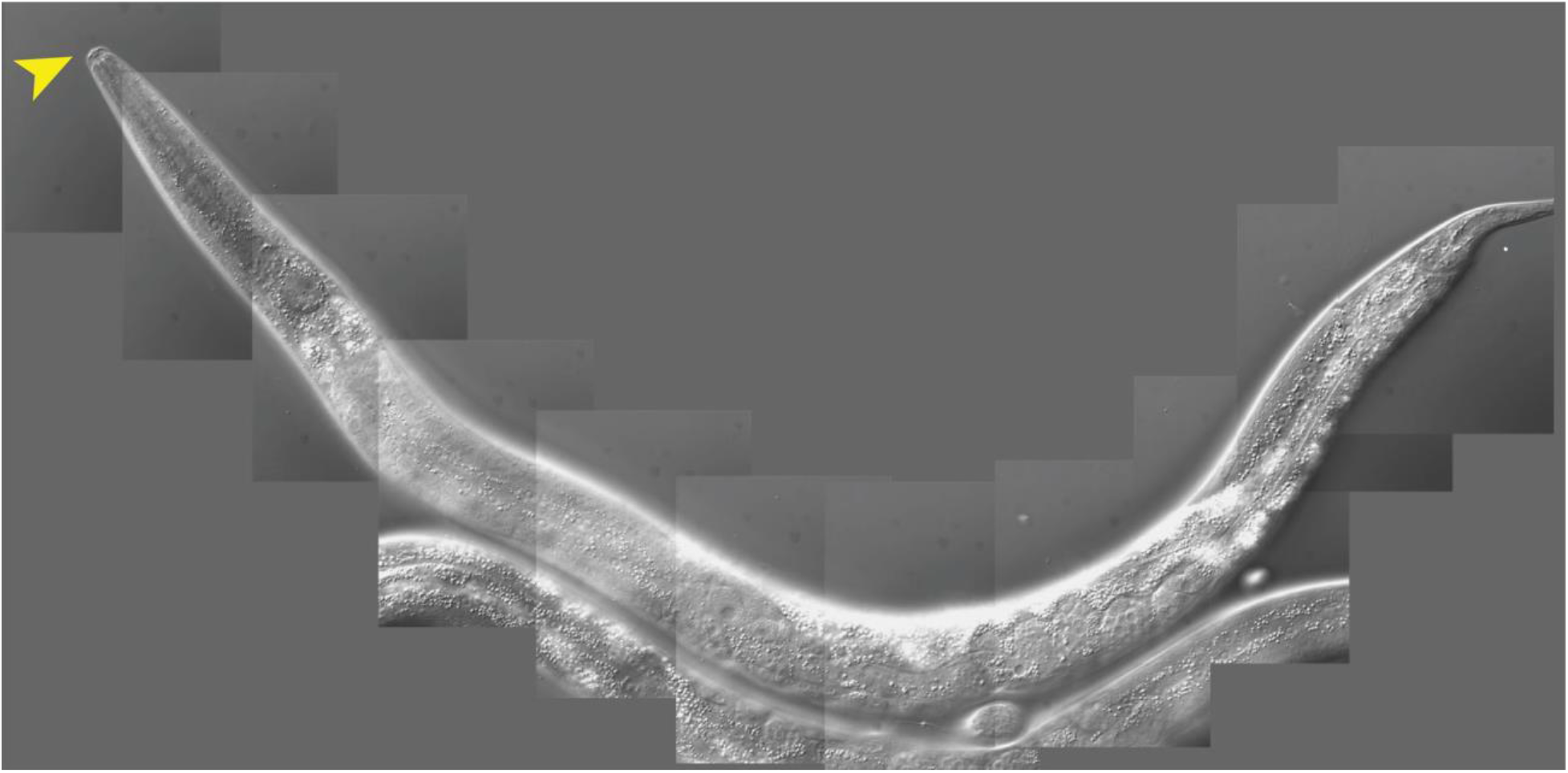
*Cbr-let-7(0)* mutants have extra molts. DIC micrograph of an adult *Cbr-let-7(ae47)* worm with cuticle coming off at the anterior end, indicated by the arrowhead.

**Fig. S20.**
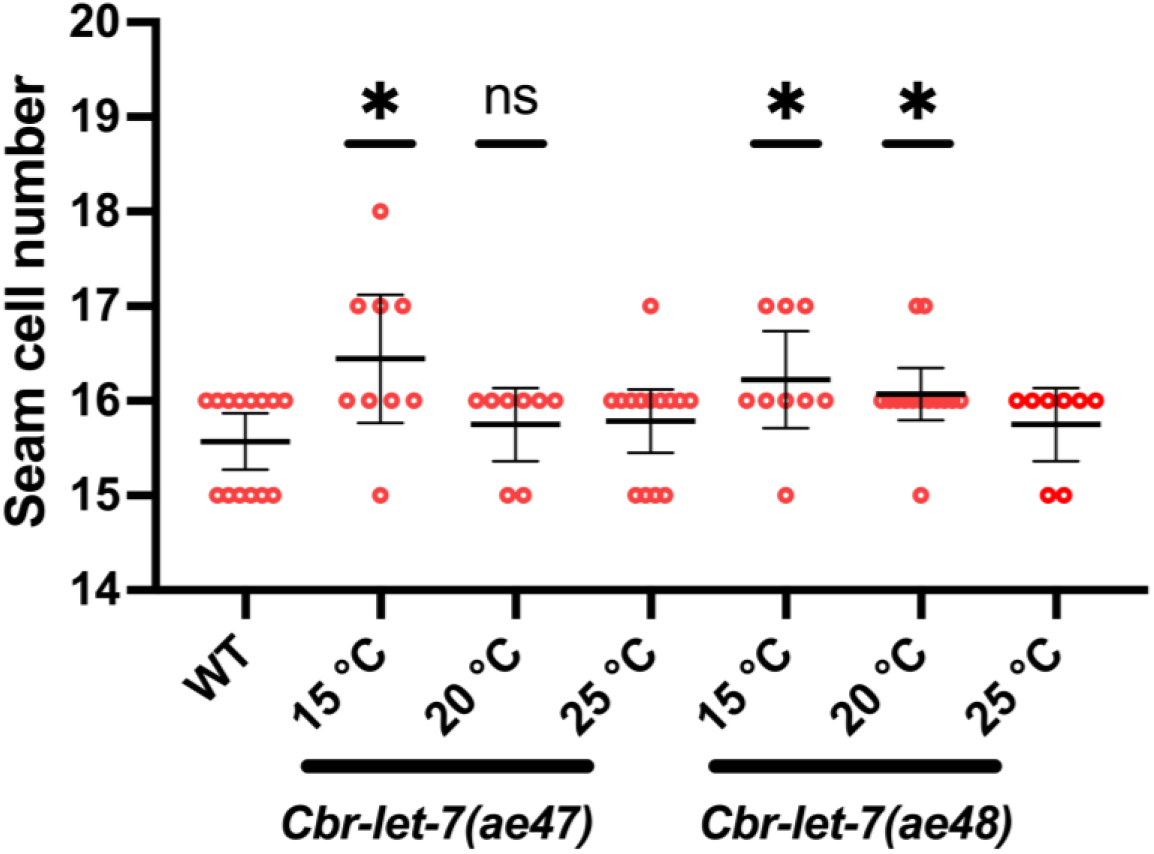
Seam cell number in *Cbr-let-7(0)* mutants is similar to wild type. Although there is an increase in the seam cell number in some animals at lower temperatures, we cannot conclude that this is due to reiteration of L2 cell fates as opposed to other reasons. Statistical analysis is described in Materials and Methods.

**Fig. S21.**
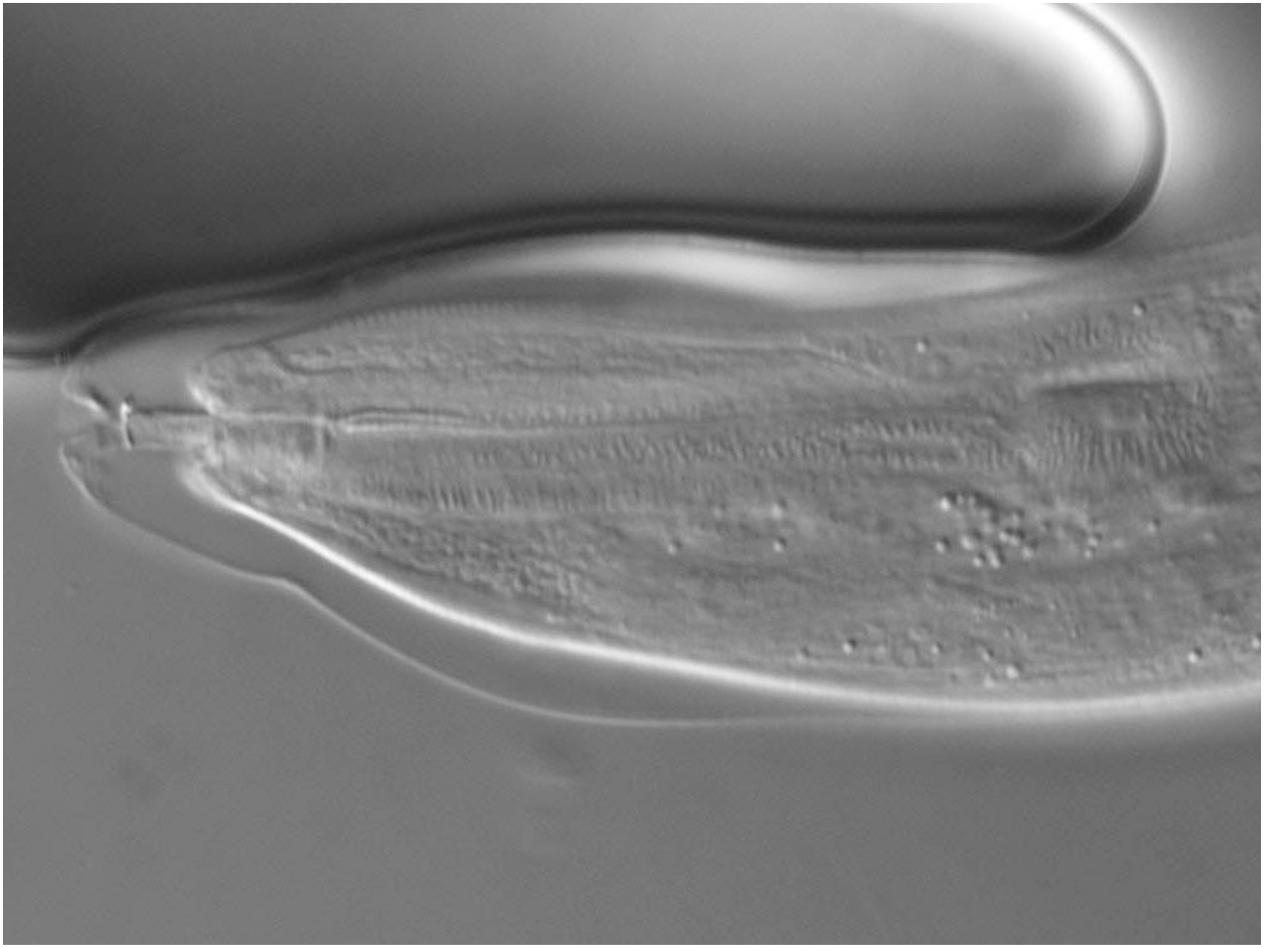
*Cbr-lin-28(0); Cbr-let-7(0)* double mutants have extra molts. DIC micrograph showing an *Cbr-lin-28(ae39); Cbr-let-7(ae47)* adult with a partly shed cuticle. Incomplete extra molts were common for this strain.

**Fig. S22.**
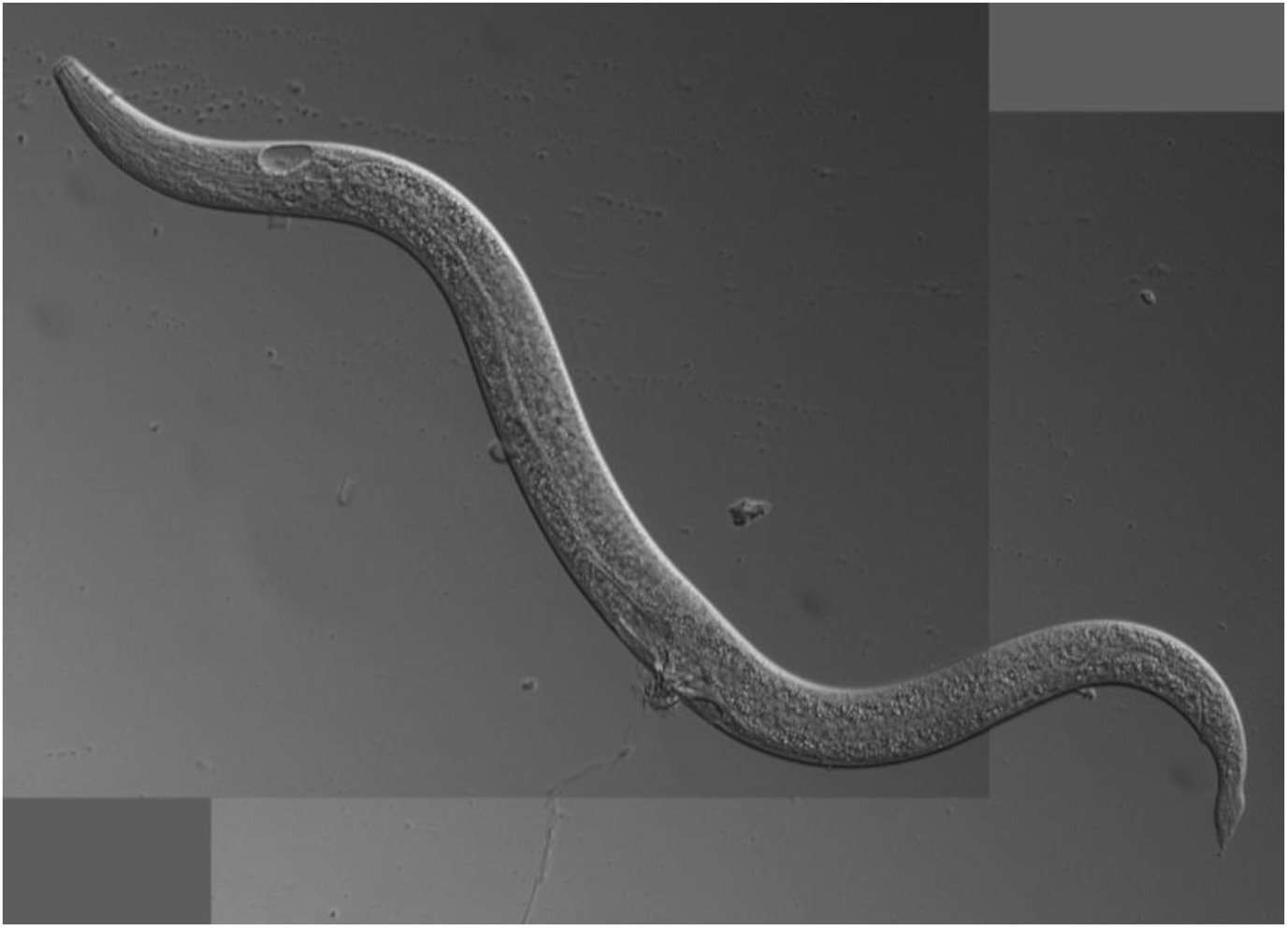
DIC micrograph of a typical *Cbr-lin-29(ae75)* adult. The abnormal vulva is typical for this mutant, but the tail is not.

**Fig. S23.**
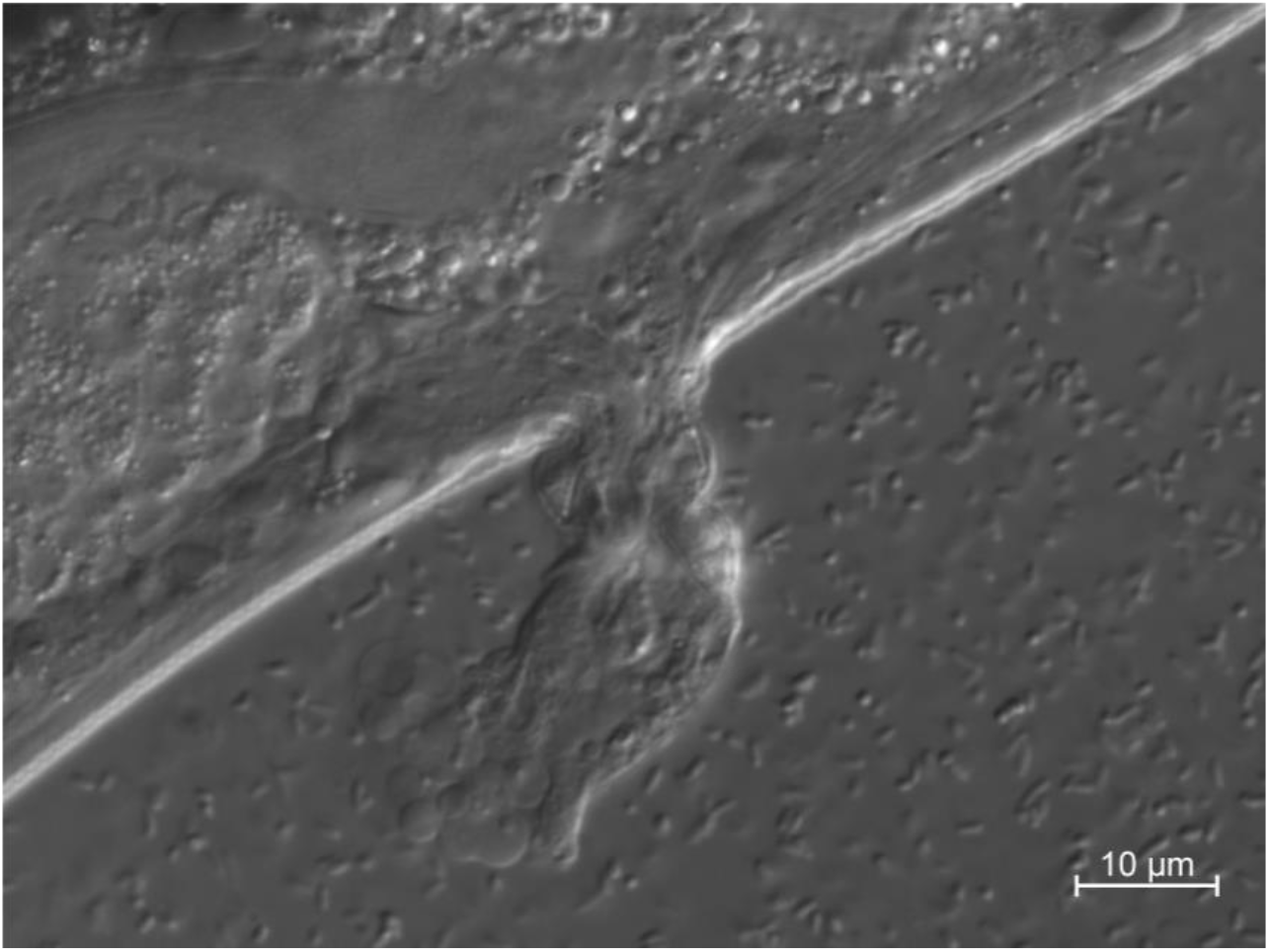
DIC micrograph showing a typical vulva of adult *Cbr-lin-28(ae39); Cbr-lin-29(ae75)* animals. The defect is similar to *Cbr-lin-28(ae39)* single mutant. Anterior end is to the left.

**Fig. S24.**
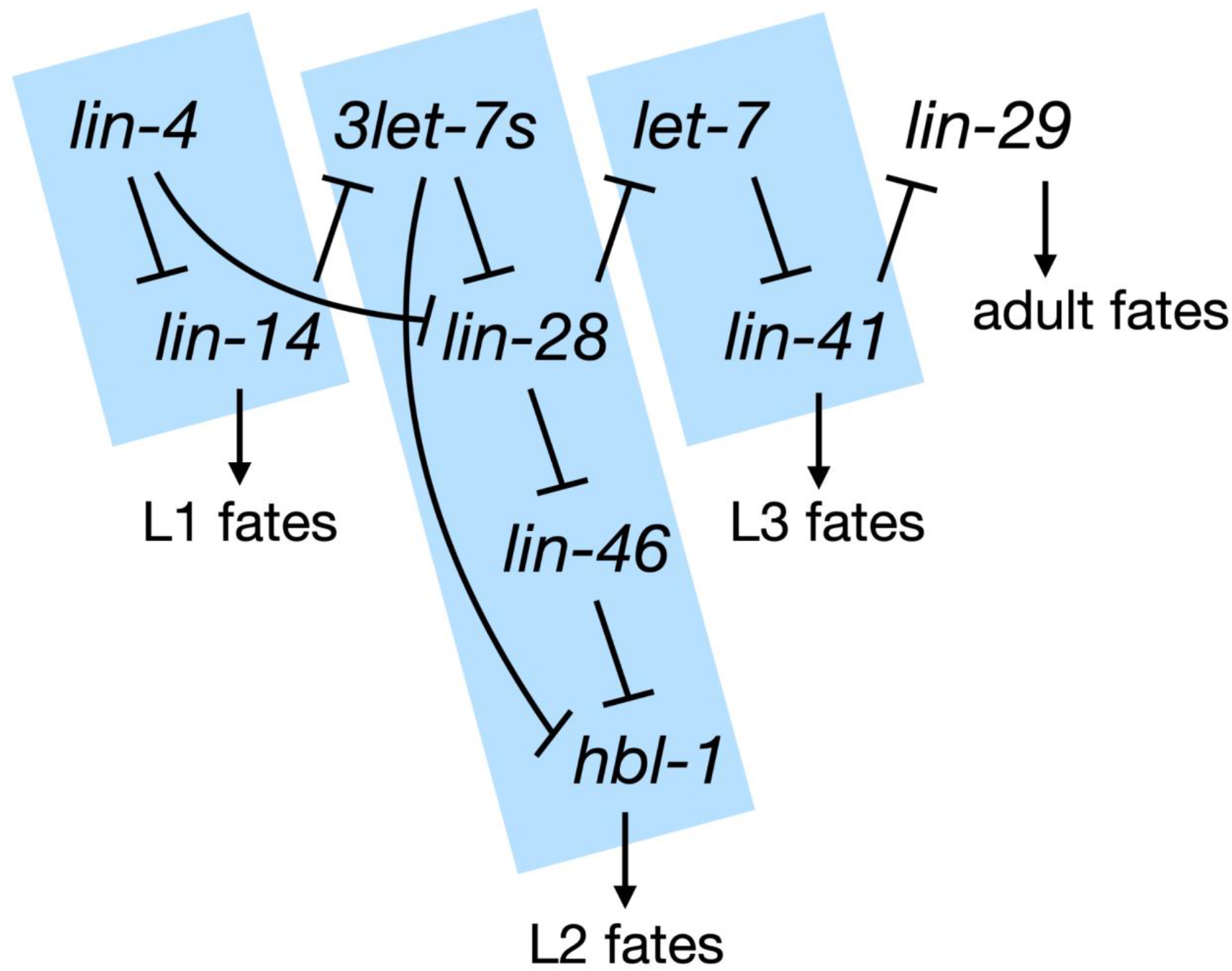
The heterochronic pathway of *C. elegans*. The scheme shows main components of the heterochronic pathway of *C. elegans* and their regulatory relationships. Bars indicate documented inhibitory activities. Blue boxes indicate the three regulatory modules. Arrows indicate the larval stages that the modules promote.

## Notes

### Competing Interest Statement

The authors have declared no competing interest.

### Summary of Updates

Added a new data on the cbr-lin-28(ae80) allele generated to confirm the null phenotype. Otherwise, minor edits, additions, and figure layout changes that improve the paper.

